# Unravelling the evolutionary relationships of hepaciviruses within and across rodent hosts

**DOI:** 10.1101/2020.10.09.332932

**Authors:** Magda Bletsa, Bram Vrancken, Sophie Gryseels, Ine Boonen, Antonios Fikatas, Yiqiao Li, Anne Laudisoit, Sebastian Lequime, Josef Bryja, Rhodes Makundi, Yonas Meheretu, Benjamin Dudu Akaibe, Sylvestre Gambalemoke Mbalitini, Frederik Van de Perre, Natalie Van Houtte, Jana Těšíková, Elke Wollants, Marc Van Ranst, Jan Felix Drexler, Erik Verheyen, Herwig Leirs, Joelle Gouy de Bellocq, Philippe Lemey

**Author notes:** Electronic address (Philippe Lemey).

## Abstract

Hepatitis C virus (HCV; genus *Hepacivirus*) represents a major public health problem, infecting about 3 % of the human population (*±* 185,000,000 people). Because no plausible animal reservoir carrying closely related hepaciviruses has been identified, the zoonotic origins of HCV still remain elusive. Motivated by recent findings of divergent hepaciviruses in rodents and a plausible African origin of HCV genotypes, we have screened a comprehensive collection of small mammals samples from seven sub-Saharan African countries. Out of 4,303 samples screened, 80 were found positive for the presence of hepaciviruses in 29 different host species. We here report 56 novel genomes that considerably increase the diversity of three divergent rodent hepacivirus lineages, which previously were almost exclusively represented by New World and European hepaciviruses. Furthermore, we provide undisputable evidence for hepacivirus co-infections in rodents, which remarkably, we exclusively but repeatedly found in four sampled species of brush-furred mice. We also point at hepacivirus co-infections indirectly in different animal hosts by demonstrating evidence for recombination within specific host lineages. Our study considerably expands the available hepacivirus genomic data and elucidates the relatively deep evolutionary history that these pathogens have in rodents compared to other mammalian hosts. Overall, our results emphasize the importance of rodents as a potential hepacivirus reservoir and as models for investigating HCV infection dynamics.

## Introduction

Diseases originating from animal sources represent a tremendous public health threat that requires sustained research effort and informed intervention measures. Owing to major advances in genome sequencing technologies, we are now able to characterize pathogen emergence and explore the interplay between viral evolution and host ecological dynamics in great detail. By obtaining viral genetic data and applying evolutionary analysis methods we can determine key factors for interspecies transmissions and successful epidemic spread in the human population. Therefore, harnessing such information may assist in effectively controlling pathogen outbreaks.

The current COVID-19 pandemic, the recent Ebola virus outbreaks, and the 2009 influenza H1N1 pandemic are just three examples that highlight the need to understand zoonotic disease emergence. To date, we still lack essential knowledge of how viruses evolve from their reservoir species, emerge into the human population and establish infections with epidemic and/or even pandemic potential. There is a pressing need to address these questions for recently emerged diseases, but it is also important to understand the origins of long-established human pathogens. An important example of the latter is Hepatitis C virus (HCV), which appears to have originated several hundred years ago (Pybus and Thézé, 2016) from a zoonotic source that remains enigmatic. Fundamental knowledge of the hepacivirus animal reservoir is essential for assessing the risk of future zoonotic spillovers into the human population.

Hepaciviruses (HVs) comprise a large group of positive-sense single-stranded RNA viruses belonging to the *Flaviviridae* family. Following its discovery in 1989 (Choo *et al*., 1989), HCV emerged as the type species of this genus. It is a blood-borne pathogen that can cause severe chronic liver disease and accounts for more than 185 million infections globally (Messina *et al*., 2015). If left untreated, HCV infection can lead to liver cirrhosis, hepatocellular carcinoma and liver failure. Because HCV infection usually produces no or only very mild symptoms during the early stages, diagnosing the infection remains challenging. As a consequence, by the time many infected individuals receive antiviral treatment, their livers can be severely damaged resulting in the need for organ transplantation. HCV infection therefore creates an enormous (economic) burden encapsulated in a major global health problem.

Although considerable research has been devoted to the optimization of curative antivirals, comparatively less effort has been put into unravelling the epidemic history and emergence of HCV. Many attempts to answer important questions about hepacivirus ecology, epidemiology and evolution remain inconclusive. For instance, it is still unclear what the zoonotic origins of this virus are, under which circumstances it crosses species barriers and how it emerged and adapted to the human population. Fundamental knowledge of how HCV has been transmitted to humans therefore remains lacking.

For a long time, HCV was the sole representative of the *Hepacivirus* genus. Since 2011, however, considerable efforts to fill the gaps in hepacivirus diversity led to the identification of HCV homologues in a wide range of animal hosts. To date, those include mammalian hosts such as: bats (Quan *et al*., 2013), cows (Baechlein *et al*., 2015; Corman *et al*., 2015), dogs (El-Attar *et al*., 2015; Kapoor *et al*., 2011), horses (Burbelo *et al*., 2012; Lyons *et al*., 2012), primates (Canuti *et al*., 2019; Lauck *et al*., 2013), possums (Chang *et al*., 2019), shrews (Guo *et al*., 2019), sloths (Moreira-Soto *et al*., 2020) and rodents (Drexler *et al*., 2013; Firth *et al*., 2014; Kapoor *et al*., 2013; Van Nguyen *et al*., 2018). Non-mammalian hosts harbouring hepaciviruses have also been identified including birds (Chu *et al*., 2019; Goldberg *et al*., 2019), fish and reptiles (Shi *et al*., 2018). Finally, divergent hepaciviruses were very recently detected in the first non-vertebrate hosts, specifically in a *Culex annulirostris* mosquito (Williams *et al*., 2020) and an *Ixodes holocyclus* tick species (Harvey *et al*., 2019).

Despite our expanding knowledge of the hepaciviral host range, the zoonotic origins of HCV still remain unresolved. The most closely related viral lineage to HCV is currently found in horses, which has also transmitted to dogs (Pybus and Thézé, 2016) and donkeys (Walter *et al*., 2017). Rodent hepaciviruses (RHVs) show the greatest genetic heterogeneity among mammalian host clades and were hypothesized to constitute the primary zoonotic source of mammalian hepaciviruses (Hartlage *et al*., 2016; Pybus and Gray, 2013; Pybus and Thézé, 2016). The idea of rodents harbouring ancestral reservoir virus species that subsequently infected other animals, has gained considerable momentum. Not only do rodents host extensive virus diversity, they also are abundant in nature providing ample ecological opportunity for spreading infectious diseases (Pybus and Thézé, 2016).

Although hepaciviruses have now been identified in a variety of hosts, our knowledge about the evolutionary dynamics of hepaciviruses is almost entirely based on HCV. It still remains unclear to what extent insights from HCV can be extended or applied to other hepaciviruses. Recombination is for example rare in HCV, but Thézé *et al*. (2015) suggested that it may have occurred in the ancestral history of different hepacivirus lineages. HCV is also known to escape immune responses during chronic infection (e.g. Gaudieri *et al*. (2006)), but whether similar virus-host interactions impact hepacivirus evolution in other hosts is currently unknown. Finally, while the HCV genome has been estimated to evolve at about 0.001 substitutions/site/year (Gray *et al*., 2011), a rate typical of RNA viruses, similar attempts to quantify the evolutionary rate of hepaciviruses in other hosts are lacking.

To scrutinize the role of small mammals as potential natural reservoir hosts of hepaciviruses, we screened 4,303 specimens from wild mammals corresponding to 161 species. We specifically focused on rodents that accounted for the majority of our sample collection. Complementary to previous research that mainly investigated rodents from Europe and the New World (Drexler *et al*., 2013; Firth *et al*., 2014; Kapoor *et al*., 2013), our sampling concentrated exclusively on Africa. The focus on sub-Saharan Africa as a source of HCV is critical because it harbours several endemic HCV genotypes and has, along with Asia, one of the highest number of cases globally. Using a comprehensive set of new rodent hepacivirus genomes, we characterize diversity, virus-host phylogenetic relationships and co-infection patterns. Finally, we take an important step towards exploring the evolutionary dynamics of hepaciviruses by examining recombination, selective pressure and temporal signal in specific host lineages.

## Materials and methods

### Sample collection and hepacivirus screening

We screened a large collection of small mammal samples previously assembled in various ecological and evolutionary studies by authors of this study and their collaborators (e.g. (Bryja *et al*., 2012, 2014; Goüy de Bellocq *et al*., 2010; Gryseels *et al*., 2015, 2017; Laudisoit *et al*., 2009; Makundi *et al*., 2015; Massawe *et al*., 2012; Mazoch *et al*., 2018; Meheretu *et al*., 2012; Petružela *et al*., 2018; Těšíková *et al*., 2017; Van de Perre *et al*., 2018, 2019b). As part of this investigation, a total number of 4,303 wild mammals (rodents, bats, shrews, elephant shrews, hedgehogs and moles) were captured in multiple localities of seven different countries across Central and East Africa between 2006 and 2013. The specimen collection used here consists of 894 animals originating from the Democratic Republic of the Congo (DRC), 426 from Ethiopia, 532 from Kenya, 30 from Madagascar, 399 from Mozambique, 1,798 from Tanzania and 224 from Zambia (Supplementary Table 1). For the majority of captured individuals (using various types of traps), whole blood was collected on pre-punched filter papers but also spleen, kidney and other organs were collected and stored in RNAlater (Qiagen) at −20°C or in ethanol at room temperature. For the hepacivirus screening, *n* = 4, 173 dried blood spots (DBS) were pooled by two and RNA was extracted using the RTP® DNA/RNA Virus Mini Kit (Stratec), according to the manufacturer’s instructions and using the maximum lysis incubation time. For *n* = 130 kidneys (all belonging to the collection from Mozambique), RNA was purified using the Nucleospin® RNA II Total RNA Isolation kit (Macherey-Nagel) following the manufacturer’s protocol. Complementary DNA (cDNA) was synthesized from either blood or tissue extracts using the Maxima Reverse Transcriptase (ThermoFisher Scientific) with random hexamers and 8 µL of RNA extract.

In order to screen for hepaciviruses, we employed a highly sensitive hemi-nested PCR assay targeting a 300-nt fragment of the conserved NS3 protease-helicase gene, as described by Kapoor *et al*. (2013). The first round of PCR was performed using the OneStep RT-PCR Kit (Qiagen) with primer pair AK4340F1 and AK4630R1 (Kapoor *et al*., 2013) and 5 µL of cDNA. The cycling conditions consisted of an additional reverse transcription step with a sequence-specific primer at 50 °C for 30 min, followed by an initial denaturation step at 95 °C for 15 min. The PCR cycle included 35 rounds of 95 °C for 30 s, 57 °C for 30 s and 72 °C for 1 min. The final extension step was performed at 72 °C for 10 min. For the second round of PCR, 1 µL of the amplified product was subjected to another PCR reaction using primer pair AK4340F2 and AK4630R2 (Kapoor *et al*., 2013) along with the DreamTaq DNA Polymerase (ThermoFisher Scientific). The PCR conditions comprised of 3 min of denaturation at 95 °C, 40 cycles of 95 °C for 20 s, 62 °C for 20 s, 72 °C for 30 s and a final extension step of 10 min at 72 °C. The quality of the PCR products was assessed visually through gel electrophoresis and in case of reasonable indication of hepacivirus presence, we subsequently purified the PCR product using the ExoSAP-IT PCR Product Cleanup Reagent (Applied Biosystems) and performed Sanger sequencing. Positive hepacivirus hits were confirmed through a similarity search against a custom viral database employing the tBLASTx algorithm.

We resolved the positive pools by individually extracting RNA from the separate DBS samples or from either kidney or spleen, depending on availability of the biological material for each individual. On these tissue extracts, an additional screening step was performed using the previously described PCR assay in order to confirm hepacivirus presence.

### Whole-genome sequencing and hepacivirus genome assembly

With available resources, we focused on obtaining viral genomic data from rodent individuals and attempted whole genome sequencing on the positive specimens when organ samples were available. Total RNA was purified from kidneys and spleens stored in RNAlater (Qiagen) at - 20°C or from other organs stored in ethanol at room temperature, using an optimised protocol of the RNeasy Mini Kit (Qiagen). In order to optimally prepare the specimens for whole genome sequencing, we adapted the original assay as detailed in Bletsa *et al*. (2019). Briefly, we introduced two freeze-thaw steps: before and after tissue homogenisation, followed by an intermediate on-column DNase treatment using the RNase-Free DNase Set (Qiagen) to remove any residual DNA prior to RNA purification. In order to increase the yield of viral RNA during extraction, we used the flow-through from the first elution to re-elute the column.

RNA extracts were subjected to RNA quantification using the RNA Quantifluor System (Promega) and their RNA profiles were assessed on an Agilent RNA 6000 Nano chip (Agilent Technologies). Prior to library preparation, a ribosomal RNA (rRNA) depletion step using the Ribo-Zero rRNA Removal Kit (Illumina) was applied to the total RNA to eliminate both cytoplasmic and mitochondrial rRNAs. For cDNA generation and construction of the sequencing libraries we used the NEXTflex Rapid Illumina Directional RNA-Seq Library Prep Kit (PerkinElmer) followed by paired-end sequencing on an Illumina NextSeq 500 at Viroscan3D (Lyon, France).

Demultiplexing was performed using Bcl2fastq v2.17.1.14 (Illumina) and low quality parts of the reads were trimmed using Trimmomatic v0.36 (Bolger *et al*., 2014). For digital host subtraction we used SNAP v1.0 (Zaharia *et al*., 2011) to map the trimmed reads against a list of 10 mammalian reference genomes (8 rodent species, shrew and human genome) coupled with PRINSEQ v0.20.4 (Schmieder and Edwards, 2011) as an additional filtering step prior to *de novo* assembly. SPAdes v3.12.0 (Bankevich *et al*., 2012) was used to generate contigs, which were subsequently analyzed using tBLASTx against a flavivirus and hepacivirus enriched database. To correct for any sequence polymorphism, we re-mapped all reads to our generated consensus sequences using Bowtie2 v2.2.5 (Langmead and Salzberg, 2012) and QUASR v6.08 (Watson *et al*., 2013). Coverage and sequencing depth were assessed by calculating the proportion of the mapped reads over the total numbers of reads.

To fill gaps in partial genomes, we designed strain-specific primers (see Supplementary Table 2) and generated overlapping amplicons using the OneStep RT-PCR Kit (Qiagen) and 5 µL of cDNA. PCR products were purified and Sanger sequenced in both directions. Open reading frames were predicted in Geneious Prime v2019.2.1 (Biomatters, 2019) based on previously characterized rodent hepaciviruses.

### Validation of co-infections

To exclude the possibility of *de novo* assembly artefacts in the newly discovered co-infections, we developed a strain-specific PCR validation assay for two different specimens (MOZ329 and TA100) that harboured three rodent hepaciviruses each. Outer and inner primer pairs were designed targeting the most variable region of the rodent hepacivirus genome (see Supplementary Figure 1 and Supplementary Table 2). All PCR reactions were performed using the OneStep RT-PCR Kit (Qiagen) with only very few differences compared to the detection assay. For the first round of PCR an annealing temperature of 52 °C was used, whereas 53 °C was the optimal temperature for the second round. Products were loaded on a 2 % agarose gel and the appropriate bands were excised and cleaned up with the Zymoclean Gel DNA Recovery Kit (Zymo Research). All purified amplicons were shipped to Macrogen Europe B.V. (Amsterdam, the Netherlands) and Sanger sequenced in both directions. Sequencing chromatographs were visually checked and the sequences were mapped to their corresponding strains. As an additional step to validate the co-infections, we meticulously tested for intra-specific recombination using SimPlot v3.5.1 (Lole *et al*., 1999) and BootScan methods (Martin *et al*., 2005; Salminen *et al*., 1995) with the default settings of a 200 bp window size and a step size of 20 bp.

**FIG. 1.**
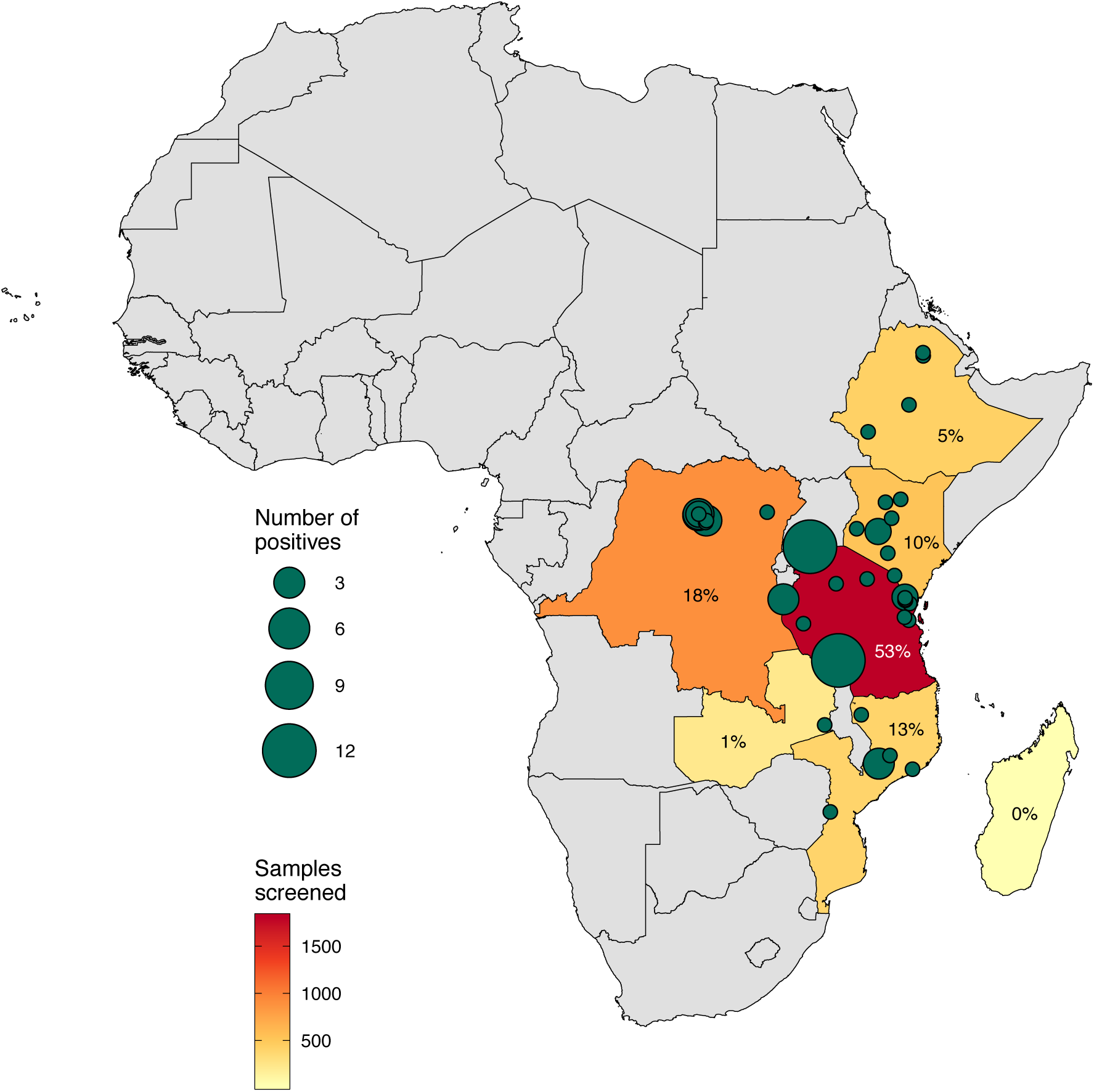
Spatial distribution of the hepacivirus-positive specimens. Map of Africa indicating sampling sites and the exact locations of our detected hepacivirus cases. Grey-coloured countries correspond to locations that were not included in this survey, while coloured countries represent our sampling focus. In those countries, the number of specimens screened is indicated by a continuous colour scale ranging from yellow (small sampling size) to red (large sampling size). Green circles denote the number of hepaciviruses detected in each locality. Circle size is proportional to the number of infected individuals, ranging from 1 to 12 positive specimens per site. For each country, we show the percentage of positive samples.

### Phylogenetic analysis and visualization

We analyzed our novel rodent hepacivirus genomes (*n* = 56) together with all available full-length hepaciviruses (*n* = 115) in GenBank (accessed on: 19/11/2019) along with information on their host, sampling location and collection date. Due to the vast number of sequences available for HCV, we only included one representative genome from each HCV genotype. This resulted in a final dataset of 171 genome-wide hepaciviruses (Supplementary Table 3). All 5’ and 3’ UTRs were removed to retain the polyprotein coding sequence for downstream analyses.

Upon translating the polyprotein sequences to amino acid sequences, we built a multiple alignment using MAFFT v7.407 (Katoh *et al*., 2009) and SeaView v4.6 (Gouy *et al*., 2010) in a stepwise approach. Firstly, we generated alignments for all main lineages defined by Thézé *et al*. (2015): equine, bovine, human, primate, bat and rodent virus lineages. Secondly, we manually edited the individual alignments in Aliview v1.18.1 (Larsson, 2014) to remove large gaps and then progressively incorporated the lineage-specific alignments into a single multiple host alignment using profile alignment. BMGE v1.12 (Criscuolo and Gribaldo, 2010) was used to eliminate poorly aligned regions and keep a fair amount of conservation within our dataset (171 sequences, 1,532 amino acids).

We used IQ-TREE v1.6.7 (Nguyen *et al*., 2015) to find the best-fitting amino acid substitution model according to the Bayesian Information Criterion (BIC), which was identified to be the LG + F + I + G4 model, and to reconstruct maximum likelihood (ML) phylogenies using this substitution model. We obtained bootstrap support using 1,000 pseudo-replicates and visualized trees as midpoint-rooted.

Amino acid alignments for the classification of hepaciviruses were prepared according to the methodology proposed by Smith *et al*. (2016), which resulted in a subset of 60 sequences. We estimated mean pairwise amino acid *p*-distances using MEGA7 v.0.1 (Kumar *et al*., 2016) for positions 1123 - 1566 in NS3 and 2536 - 2959 in NS5B. Phylogenetic trees were reconstructed for both regions with IQ-TREE v1.6.7 (Nguyen *et al*., 2015) using the LG + F + I + G4 substitution model and 1,000 bootstrap replicates.

To molecularly confirm the host species of the positive specimens, we recovered cytochrome b gene sequences in the samples subjected to whole genome sequencing by directly mapping the deep sequencing data to a list of reference sequences from various African rodent species. Some of these species have not yet been formally named, but we used expert opinion to delineate the different species. In addition to those, we downloaded cytochrome b sequence data of the 12 rodent species from which hepacivirus genomes were sequenced in previous studies (see Supplementary Table 4).

Phylogenetic trees based on the alignment of 21 cytochrome b sequences were estimated with IQ-TREE v1.6.7 (Nguyen *et al*., 2015) using the TIM2 + F + I + G4 nucleotide substitution model (identified as the best model according to the BIC) and clade support was assessed using 1,000 bootstrap replicates.

To visualize and annotate phylogenies we made use of *ggtree* and *treeio* R packages (Wang *et al*., 2019; Yu *et al*., 2018). In order to investigate the relationships between rodents and hepaciviruses we created a co-phylogenetic plot (or “tanglegram”). This visual representation plots the host phylogeny opposite to the virus phylogeny and draws lines between the taxa of the two trees, as a function of their topological distance. Here, we focused on highlighting the evolutionary relationships of rodent-borne hepaciviruses and their hosts only, as those mammals were exclusively associated with multiple circulating hepacivirus strains (co-infections). Briefly, for the tanglegram we constructed a viral phylogeny based on a subset of all rodent hepaciviruses (*n* = 78) from the the large dataset using the approach described above. The association matrix between the host and viral phylogeny was computed using the *ape* R package (Paradis and Schliep, 2018) and patristic distances were calculated using the *adephylo* R package (Jombart *et al*., 2010).

### Recombination, selective pressure and temporal signal in host-specific lineages

For comparative evolutionary analyses we selected viral genomes representative for specific host lineages from the complete hepacivirus phylogeny that are roughly similar in diversity (see Supplementary Figure 2). This includes the entire collections of bovine (*n* = 16) and equine (*n* = 34) strains and subsets of two rodent lineages (*n* = 11 and *n* = 36 respectively). To represent HCV, we collected representative genome data sets of similar sizes for HCV genotype 1a (*n* = 35), genotype 1b (*n* = 34) and genotype 3a (*n* = 34) by applying phylogenetic diversity analyzer (Chernomor *et al*., 2015) to the large number of genomes publicly available in Genbank. Multiple sequence alignments were constructed for the amino acid translations of the polyprotein coding sequence using Muscle v3.8.31 (Edgar, 2004) and back-translated to the nucleotide sequences. A relatively short but highly diverse part in the 3’ region of one of the rodent lineage aligments was removed by manual editing. To ensure comparable data in the evolutionary analyses, the equivalent part was removed from the other host-lineage alignments.

We tested for recombination in those host-specific lineages using the PHI-test (v4.15.1) (Bruen *et al*., 2006) and confirmed the evidence of significant recombination using RDP4 v4.97 (Martin *et al*., 2015). The RDP analysis employed the following individual methods: 3SEQ (Lam *et al*., 2017), RDP (Martin and Rybicki, 2000), Bootscan (Martin *et al*., 2005), Chimaera (Martin *et al*., 2005) and SisScan (Martin *et al*., 2005). For RDP, Bootscan and SisScan a window size of 200 bp was selected, while for Chimaera we allowed for a number of 20 variable sites per window. Apart from specifying linear genomes and recombination events to be listed by more than two methods, all other parameters were kept to their default settings.

Due to the detection of a significant amount of recombination in the viral genomes from animal hosts, we estimated selection pressure at the molecular level using the population genetics approach implemented in omegaMap v0.5 (Wilson and McVean, 2006). This method was specifically designed to estimate the relative excess of nonsynonymous (dN) over synonymous (dS) substitutions in the presence of recombination. Selection analysis was performed on the same data sets (see Supplementary Figure 2) that have been used to screen for recombination. In our estimates we allowed for variation in dN/dS ratio according to a block-like model with length of 30 sites. We set the codon frequencies to the values obtained by multiplying the four empirical nucleotide frequencies that are obtained separately for the three codon positions.

To test for temporal signal, we focused on the genomes in the aforementioned lineage-specific data sets for which sampling time was available. This information was retrieved for all HCV genomes, the rodent virus genomes (exact sampling dates), and for 14 out of 16 bovine virus genomes, as well as for 22 out of 34 equine virus genomes. To avoid the impact of recombination, minor recombinant regions were masked based on the RDP4 analysis, keeping only the major non-recombinant regions in the lineage-specific alignments. Temporal signal was assessed in a visual manner and using a Bayesian testing procedure. For the visual assessment, we plotted root-to-tip divergences as a function of sampling time using TempEst v1.5.3 (Rambaut *et al*., 2016) based on ML trees inferred by IQ-TREE under a GTR + G4 nucleotide substitution model. A more formal test of temporal signal was performed by comparing marginal likelihood estimates for a model with dated tips and a model that assumes all sequences are contemporaneous (Duchene *et al*., 2019).

## Data availability

Genomic sequences generated in this study were deposited in GenBank under the following accession numbers:

- 56 rodent hepacivirus genomes (accession numbers: MN555567, MN564789 - MN564794, MN587650 - MN587698)
- 25 rodent cytochrome b sequences (accession numbers: MN616976 - MN616980, MN616983 - MN616986, MN616989 & MN616990, MN616993 - MN616995, MN616998 - MN617000, MN617005 - MN617007, MN617010 - MN617014)

## Results

### Hepaciviruses are present in a wide range of rodent host species

We screened a comprehensive set of small African mammals (*n* = 4, 303), collected between 2006 and 2013, for the presence of hepaciviruses. The capture efforts culminating in our sample collection were undertaken in Central and East Africa and included 61 different small mammal genera that further classify into 161 potential species. The vast majority of specimens were from rodents (*n* = 3, 788) constituting 88% of the whole collection, which to our knowledge, is the most extensive hepacivirus screening of African rodents to date. Our rodent sample set consists of 38 genera that are further divided into 116 potential rodent species with the *Mastomys* (*n* = 888) genus being the most frequent followed by the *Praomys* (*n* = 641), *Rattus* (*n* = 335) and *Lophuromys* (*n* = 303) genera. In addition to rodent specimens, *n* = 515 samples from shrews, bats, elephant shrews, hedgehogs and moles were screened for hepaciviruses (see also Suppl. Table 1).

The molecular screening resulted in a total number of 80 positive specimens from 29 extant host species: 78 were identified across 28 potential species of *Rodentia* and 2 were found in bat individuals from one single host species. Interestingly, we did not detect any positive shrew samples, although these mammals have been previously reported to host hepaciviruses (Guo *et al*., 2019). In Supplementary Fig. 3a, we summarize how our work substantially extends the species range of rodents as hepacivirus hosts. Particularly in the Muridae family, which harbours more than 834 extant species (Burgin *et al*., 2018), we identified 24 new rodent hosts of hepaciviruses, and four additional rodent species in the Bathyergidae (suborder Hystricognathi), Spalacidae, Nesomyidae and Gliridae families have been detected to harbour hepaciviruses (Suppl. Fig. 3a). Therefore, we do not only expand the current rodent sampling at a species level, but we also contribute to a higher host taxonomic hepacivirus detection. The percentage of positive samples varied considerably by geographic region ranging from no positive individuals in Madagascar (but only 30 samples were tested) to 2.51% in Mozambican rodents (Suppl. Fig. 3b). For a summary of our screening results by country we refer to Supplementary Figure 3; Supplementary Table 5 provides a detailed list of all hepaci-positive specimens.

We mapped our screening results to further investigate the geographic distribution of the positive specimens (Fig. 1). This highlights three distinct localities with an exceptionally high rate of rodent hepacivirus infections: two sites in Tanzania and one in Mozambique. A considerable number of hepaciviruses were discovered at the Minziro Forest Reserve which lies in the northwestern part of Tanzania (coordinates: 1°01′ 51.6”*S*31°34′ 19.2”*E*). In this location, we detected a total of12 small mammals with hepaciviruses: 4 *Lophuromys stanleyi* and 6 *Praomys jacksoni* rodents and 2 bat specimens of the *Glauconycteris atra* species out of 23 samples tested (52% positives). The second spot is situated in Mount Ngozi (coordinates: 9°02′ 26.7”*S*33°34′ 23.5”*E*), where 12 rodents were identified to be hepaci-positive, all belonging to the species *Lophuromys machangui*. We tested a total number of 47 samples from this particular location (26% positives), out of which 17 belonged to the *Lophuromys* genus. In addition to those, 6 *Lophuromys machangui* individuals carried hepaciviruses in Mount Mabu out of the 43 mammals screened (14% positives), which is located at the northeastern part of Mozambique (coordinates: 16°18′ 20.5”*S*36°25′ 26.8”*E*).

### Evolutionary relationships of hepaciviruses with a focus on rodent hosts

We generated 56 complete hepacivirus genomes originating from 9 rodent species using an untargeted deep sequencing procedure (see also Supplementary Table 3). The mean read depth for the 56 genomes was 355.3 with more than 91.6% of each genome having 20x coverage.

Phylogenetic analysis of a data set that combines all available hepacivirus genomes with our new data confirms an extensive virus diversity that is to some extent structured by vertebrate class, order and family into host-specific clades (Fig. 2). All mammalian hepaciviruses form a monophyletic cluster, but with low bootstrap support. Bird, fish and reptile hepaciviruses form lineages basal to to the mammalian-borne clade and exhibit long branches, which suggests long-term circulation across those hosts. Although birds occupy a relatively basal position in the evolutionary history of vertebrates, hepaciviruses present in these hosts appear well confined to this vertebrate class (Fig. 2).

**FIG. 2.**
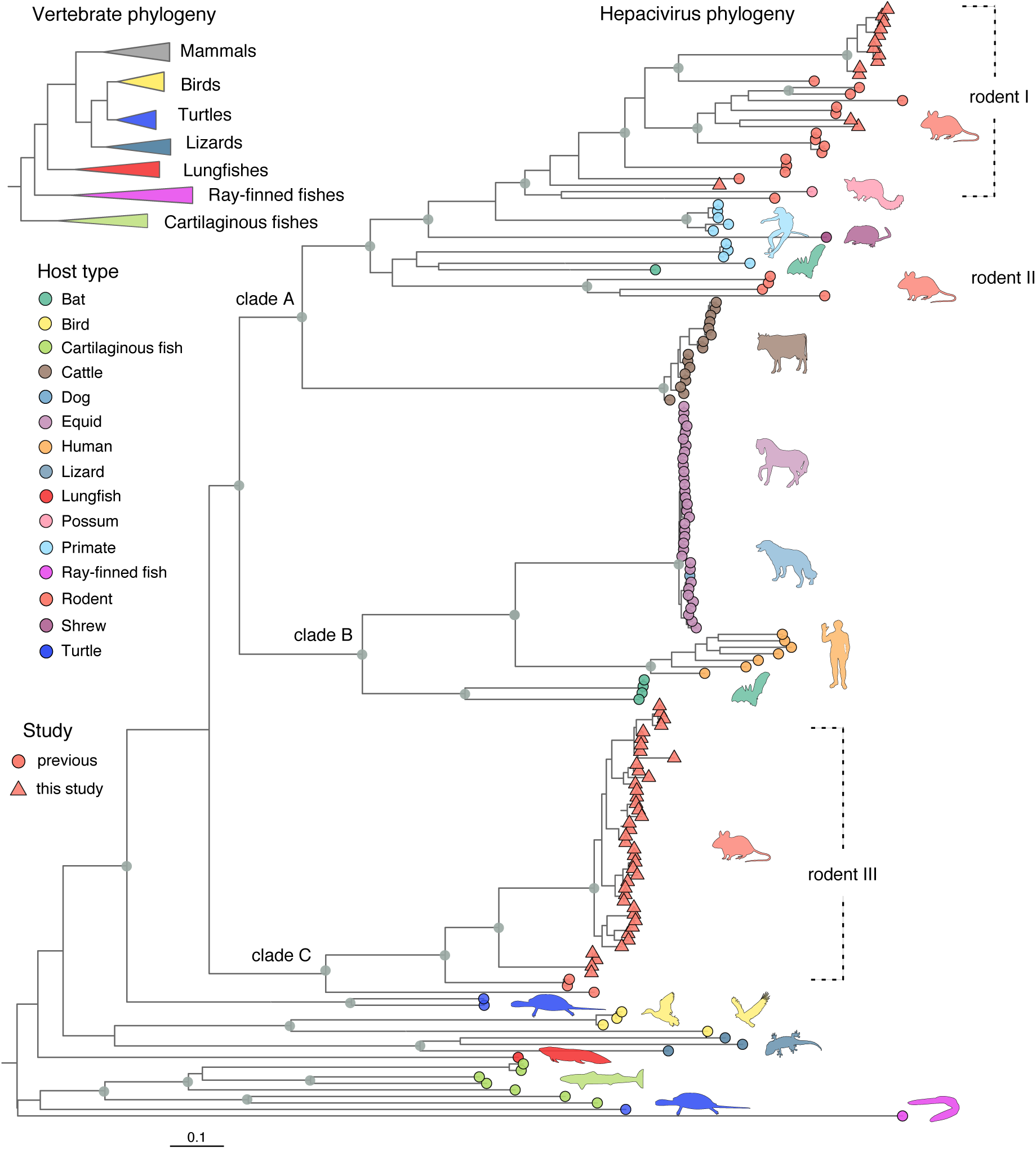
Genome-wide phylogenetic reconstruction of hepaciviruses. ML tree of all available (*n* = 115) and novel (*n* = 56) hepacivirus genomes. Silhouettes indicate hosts and are coloured according to their broader host type: bats (green), birds (yellow), cartilaginous fishes (lime green), cattles (brown), dog (grey blue), equids (lilac), humans (peach orange), lizards (steel blue), lungfish (red), possum (pastel pink), primates (light blue), ray-finned fish (fuchsia), rodents (salmon), shrew (plum), turtle (marine blue). Grey circles indicate internal nodes with bootstrap support *≥*70, while circles at the tips represent previously published sequences. Salmon triangles represent the novel genomes generated in this study. A simplified version of the vertebrate phylogeny is reproduced from (Irisarri *et al*., 2017) and has been minimally adapted to reflect their evolutionary clustering. Clades A, B and C have been provisionally named for the purpose of discussing the mammalian hepacivirus lineages in the main text.

We distinguish three well supported clades (provisionally named A, B and C) in mammals for the purpose of description. From the perspective of mammalian orders, the clustering of viral lineages does not adhere to the established host clustering (Fig. 2). In particular, clade A represents the most heterogenous group and is characterized by an intermingling of viruses found in a wide range of taxonomically diverse animals such as rodents, bats, shrews, non-human primates, possums and cattle. Clade B exhibits some degree of clustering by species and contains viruses originating from equine, canine, human and bat hosts, while clade C comprises a strictly monophyletic group of rodent hepaciviruses. Bat hepaciviruses (BHVs) form two distinct lineages with few representatives that cluster in both clades A and B, while RHVs fall in three divergent lineages in the phylogeny (Fig. 2). In particular, two separate RHV lineages can be identified in clade A while the remaining lineage is responsible for the entire C clade. In further analyses below, we refer to the rodent-borne lineages in clade A as ‘rodent I’ and ‘rodent II’, containing viruses that are paraphyletic with respect to non-human primate, bat, shrew and possum viruses, and to the third RHV lineage as ‘rodent III’ (equivalent to clade C).

According to the mean pairwise patristic amino acid distances, primates and rodents exhibit the highest hepacivirus diversity followed by bats (Fig. 3), although the last-mentioned are represented by fewer samples compared to the other two orders. Despite the overall pairwise divergence being similar between primates and rodents, the pairwise divergence within many sampled rodent species is substantially greater than within primate and bat species. For the non-human primates and for the bat species one could argue that perhaps this lack of within-host-species diversity is due to only limited samples available per species (*n* = 3 *–* 5), which are derived from the same locality. However, some of the rodent species that were sampled also have very few samples available (*n* = 3 *–* 6) and these are also derived from the same locality, yet are still divergent. This is particularly noticeable when we compare HV diversity of the non-human primate host species *Propithecus diadema* or *Colobus guereza* to the RHV diversity within the *Lophuromys stanleyi* species. Especially interesting is the comparison between the limited diversity of human hepaciviruses, which have been extensively sampled across multiple localities globally, and the much higher levels of HV diversity hosted by rodent species with narrower geographic range limits (e.g. *Myodes glareolus* or *Lophuromys stanleyi*).

The ever-expanding genomic characterisation of hepaciviruses is challenging to subject to systematic characterisation. To illustrate this, we attempted to apply the classification criteria by Smith *et al*. (2016), which focus on two relatively conserved subgenomic regions: part of NS3 and of NS5B (Supplementary Table 6). This resulted in an impractically large number of different virus ‘species’, many of which were represented by only a single taxon, and a large degree of inconsistency between the two genome regions despite congruent tree topologies (Supplementary Figure 4). It therefore remains more practical and coherent to describe lineages by strongly supported evolutionary units, which is also in line with the interspecific level previously used by Thézé *et al*. (2015).

As demonstrated in Supplementary Figure 5, our novel hepaciviruses fall into the major rodent lineages that were previously, disproportionally represented by New-World and Asian viruses. More specifically, the diverse rodent I cluster did not include any African viruses, while the rodent II lineage contained only 2 African genomes. In addition, rodent III clade was exclusively represented by European and Asian viruses. The large number of African hepacivirus genomes that contribute about 70% of the current RHV genomes increases the Old-World diversity of those lineages and substantially broadens the known host spectrum of RHVs, especially in the Muridae family. Of particular interest is an isolate that originated from a *Graphiurus kelleni* sample collected in the DRC (GenBank accession number MN564789). This rodent species belongs to the Gliridae familly, which has not been included in any of the previous screening efforts. The closest relative of this strain is a hepacivirus from a *Rattus norvegicus* sample that was isolated in the USA, indicating the deep evolutionary trajectory of RHVs.

In terms of geographic structure, the rodent I and II lineages show an intermingling of hepaciviruses from different continents without any clear patterns of co-divergence between the viruses and their rodent hosts. Contrary to the other two lineages, rodent III lineage (or clade C) exhibits some degree of confinement to specific rodent taxa since one subclade of this lineage is restricted to the *Lophuromys* genus and another subclade is restricted to the Praomyini tribe (Lecompte *et al*., 2008). Hepaciviruses in the rodent III lineage are exclusively sampled from Old-World locations and demonstrate geographic clustering by continent, but with mixing by country in Africa (Supplementary Figure 5). The large diversity and wide distribution of RHVs as well as the lack of a clear geographic and host structure suggest a relatively long-term circulation history with little boundaries to transmission among different rodent hosts.

### Hepacivirus co-infections within *Lophuromys* rodents

We identified a remarkably large proportion of RHV-positive samples harbouring multiple divergent strains, suggesting a high degree of hepacivirus co-infections among those rodents. Specifically, 11 *Lophuromys* individuals were found to carry from two up to five different hepaciviruses, while co-infections were not found in any other sampled genus (see also Suppl. Table 7). To exclude the possibility that these multiple genomes could have been generated by assembly artefacts, we performed additional molecular assays and computational analyses. The *in silico* validation included different assembly algorithms to *de novo* reconstruct our sequencing data. The majority of algorithms resulted in multiple hepacivirus strains per sample, albeit with variabilities in the length of the generated scaffolds. As *in vitro* validation test, we developed a strain-specific PCR assay to examine two cases harbouring three hepaciviruses each. Sanger sequencing after strain-specific PCR assays targeted to the hypervariable regions confirmed the presence of three distinct hepacivirus strains in each of the two tested specimens for which co-infections were inferred from metagenomic sequencing. This supports that our metagenomic sequencing protocols and our bioinformatic pipeline indeed reliably retrieved distinct hepacivirus genomes (and thus co-infections) within single specimens. Phylogenetic relationships among those RHVs are depicted in (Fig. 4), while a summary of the multiple isolates per individual can be found in Suppl. Table 7.

**FIG. 3.**
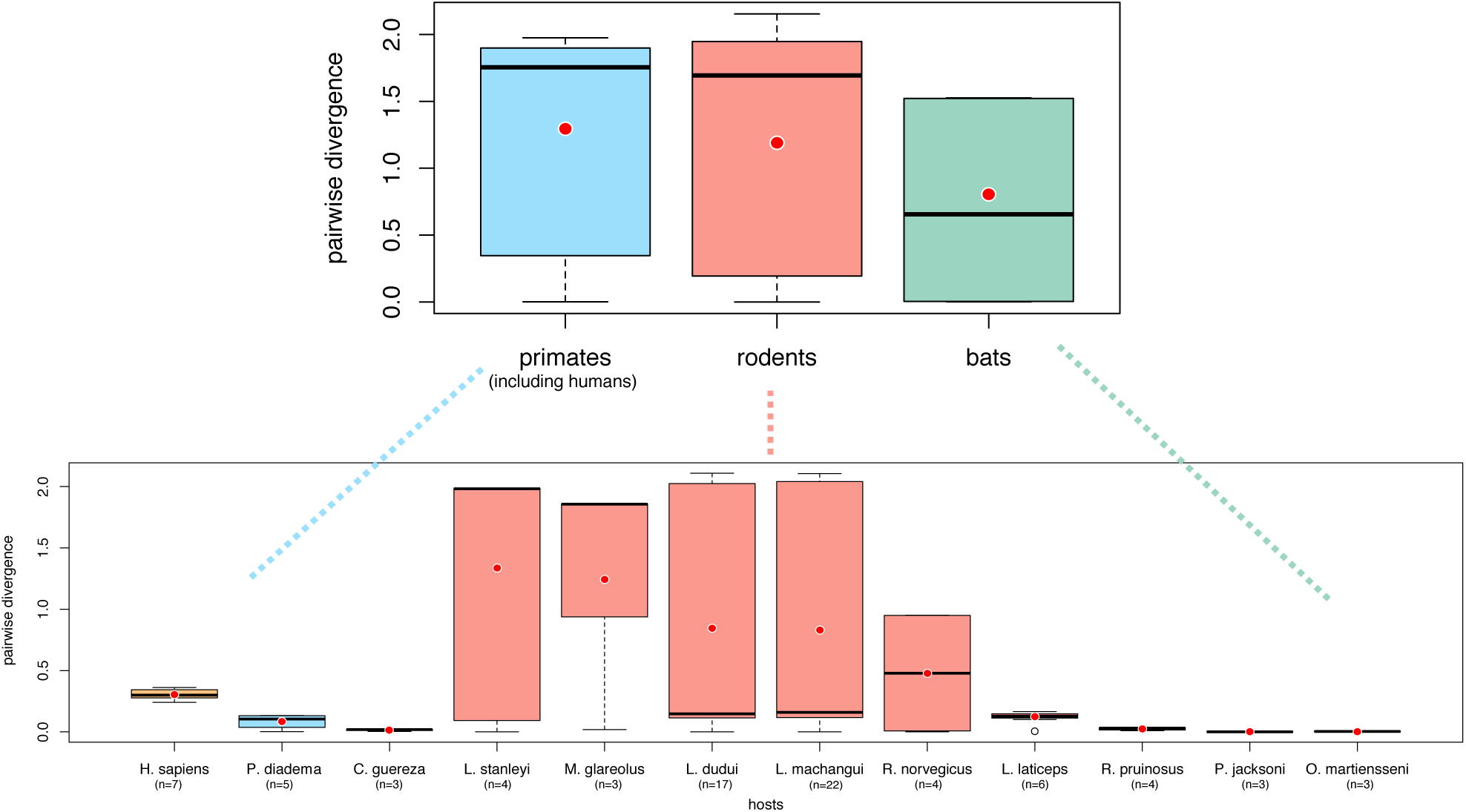
Hepacivirus diversity in primates, rodents and bats. Patristic pairwise distances were estimated from the genome-wide phylogeny of all available (*n* = 115) and novel (*n* = 56) hepacivirus genomes for the host orders that constitute the major possible hepacivirus reservoirs. Diversity of hepaciviruses by host species was estimated only for the mammalian species having more than two available hepacivirus genomes. Boxplots summarize the pairwise amino acid distances within the different hosts and are coloured according to (Fig. 2). The horizontal black lines inside the box correspond to the median patristic distance, while the red dots denote the mean values, which corresponds to a standard measure of diversity.

**FIG. 4.**
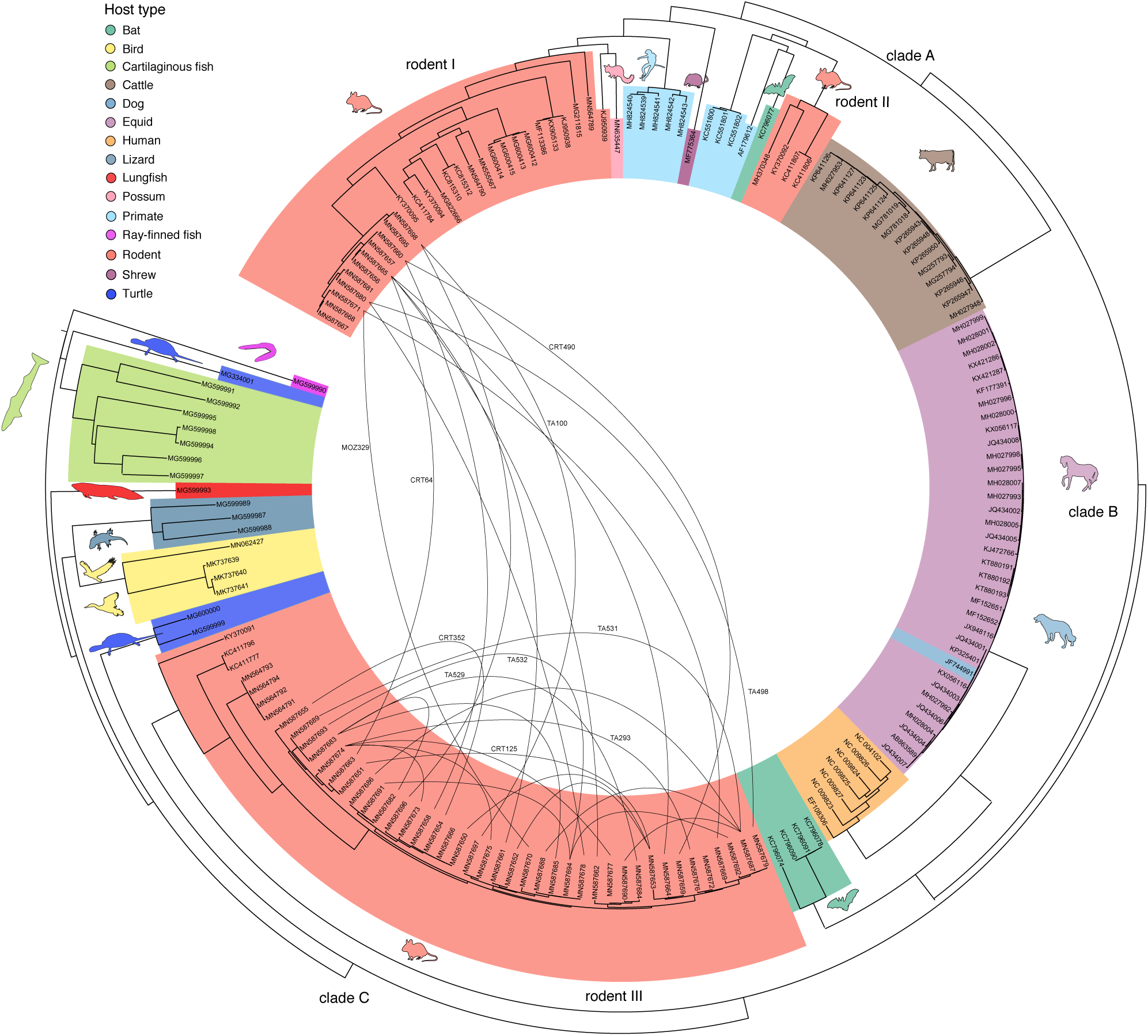
Hepacivirus co-infections in rodents. Circular ML tree of all available (*n* = 115) and novel (*n* = 56) hepacivirus genomes. Silhouettes indicate hosts and are coloured according to their broader host type: bats (green), birds (yellow), cartilaginous fishes (lime green), cattles (brown), dog (grey blue), equids (lilac), humans (peach orange), lizards (steel blue), lungfish (red), possum (pastel pink), primates (light blue), ray-finned fish (fuchsia), rodents (salmon), shrew (plum), turtle (marine blue). Interspersed lines connect the RHV genomes obtained from the same animals (co-infections). For example, rodent individual CRT352 harboured two hepaciviruses with GenBank accession numbers: MN587654 and MN587655.

Molecular identification of rodent hosts resulted in a wide species spectrum that can be broadly divided into three groups, as highlighted in Fig. 5: a) Those belonging to the Deomyinae subfamily: *Acomys wilsoni, Lophuromys dudui, Lophuromys laticeps, Lophuromys machangui* and *Lophuromys stanleyi*, b) those belonging to the Murinae subfamily: *Mastomys natalensis, Praomys jacksoni* and *Stenocephalemys albipes* and c) the one belonging to the *Graphiurus kelleni* species within the Gliridae family. Despite the broad host spectrum elucidated by our screening, a substantial proportion (55%) of hepaci-positive individuals were identified in the four species of *Lophuromys* rodents, and only these rodents were found to harbour more than one RHV strain. To our knowledge, this is the first time that hepacivirus co-infections have been described in any of the identified non-human host species. Fig. 5 highlights the viruses found in co-infections (blue lines) in a virus-host phylogenetic comparison, indicating that all co-infections were associated with only four potential species of the *Lophuromys* genus. Comparison of the rodent host phylogeny to the corresponding RHV phylogeny does not demonstrate any appreciable co-divergence patterns (Fig. 5), suggesting again that these viruses have frequently jumped rodent hosts throughout their evolutionary history and that they may transmit relatively easily between different rodent species and genera under the appropriate ecological opportunity.

**FIG. 5.**
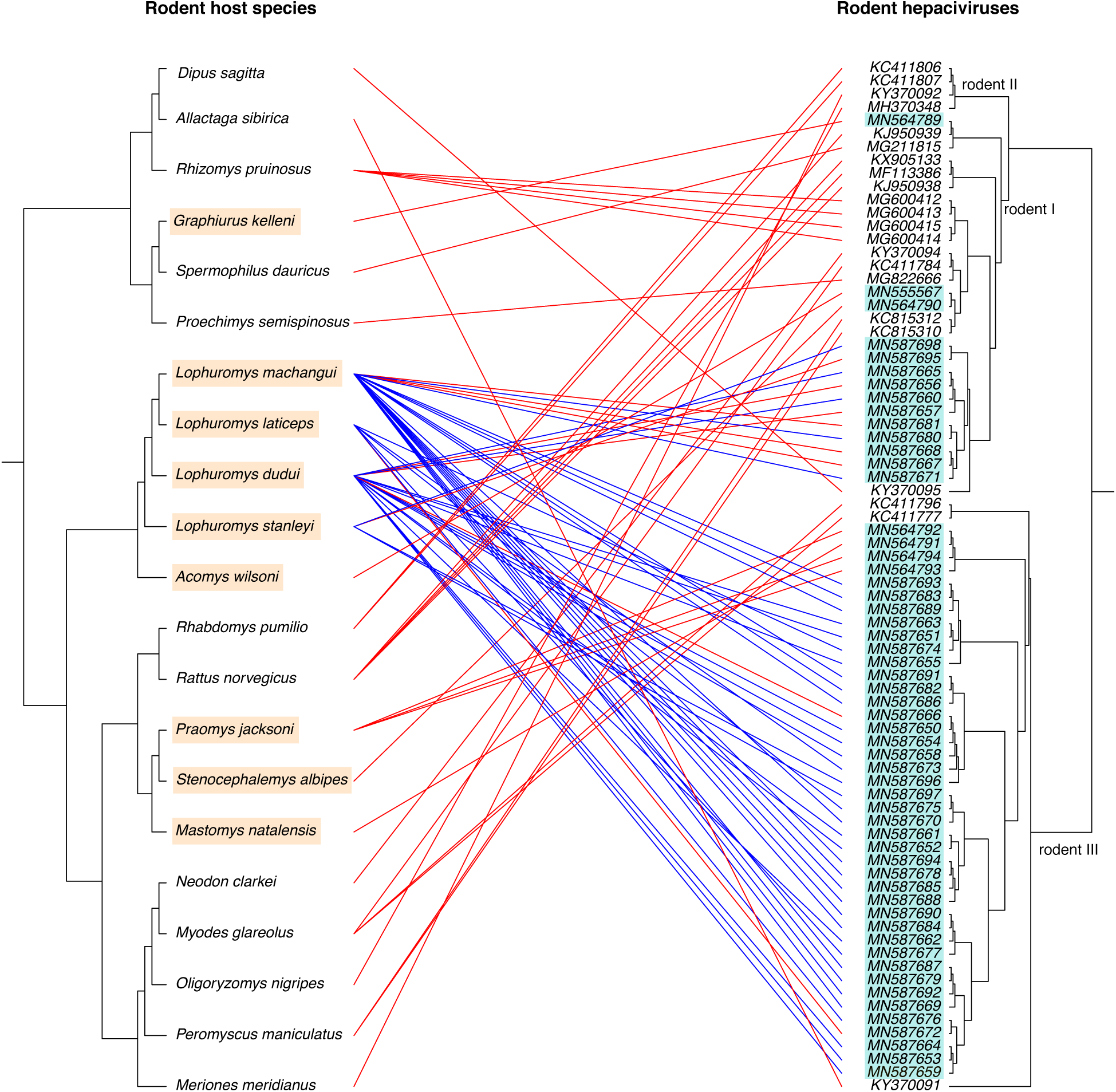
Tanglegram of rodent hosts and their hepaciviruses. The topology of the host tree was reconstructed using the cytochrome b gene from 21 rodent species (left phylogeny). For the viral reconstruction we used the rodent subset of our genome-wide alignments (right phylogeny) and we highlighted the novel RHV genomes in turqoise. Lines connecting the two phylogenies represent an association between the rodent host species and their identified hepaciviruses. Blue lines correspond to individuals harbouring multiple hepaciviruses, while rodent species highlighted in a caramel colour represent the novel hosts found carrying hepaciviruses in our study.

### Intraspecific recombination, prevailing negative selection and absence of temporal signal

With our additional RHV sampling, we assess recombination within host lineages (those lineages specific to a host type), where co-infections are more likely as indicated by our findings in the *Lophuromys* genus, and where high sequence divergence is less of a cofounding factor. We performed comparative analyses on host-specific data sets with relatively limited and comparable genetic diversity (Supplementary Figure 2). Formal testing using the PHI-test (Bruen *et al*., 2006) provided significant evidence for recombination in the bovine, equine, and the rodent I and III lineages (*p <* 0.01), but not in the three HCV data sets (1a, 1b and 3a) we included for comparison.

A substantial number of intraspecific recombinants were identified in rodents, with the highest proportion in strains circulating in the rodent III lineage. However, we did not detect any significant evidence for recombination among RHV genomes within any of the co-infections we identified. These results were also confirmed by a variety of methods implemented in RDP4 (Martin *et al*., 2015). For more details on specific recombinants and mosaic patterns found in each host lineage, we refer to the Supplementary information.

By focusing on specific host lineages, we can also perform genome-wide comparative analyses of selective pressure. At the interspecific level, such analysis would only be able to focus on conserved parts in which third codon positions may still suffer from saturation (Thézé *et al*., 2015). Because the presence of intraspecific recombination complicates widely-used phylogenetic codon substitution methods, we adopted a population genetic approach to estimate the ratio of nonsynonymous (dN) over synonymous (dS) substitutions in the presence of recombination (Wilson and McVean, 2006) (Fig. 6). The genome-wide estimates of dN/dS ratio (or ω) indicate a generally strongly negative selective pressure with average values ranging from 0.015 to 0.035 in the non-human hosts and 0.055 to 0.067 in the human host (grey horizontal bars in Fig. 6 with a Y-axis on a log-scale). The bovine data set was the only non-human data set for which the site-specific estimates provide evidence for two sites with an ω value significantly larger than 1. In contrast, a non-negligible number of positively selected sites (ranging from 20 to 25) was consistently identified in the HCV data sets, primarily located in the antigenically-important E1/E2 gene region.

**FIG. 6.**
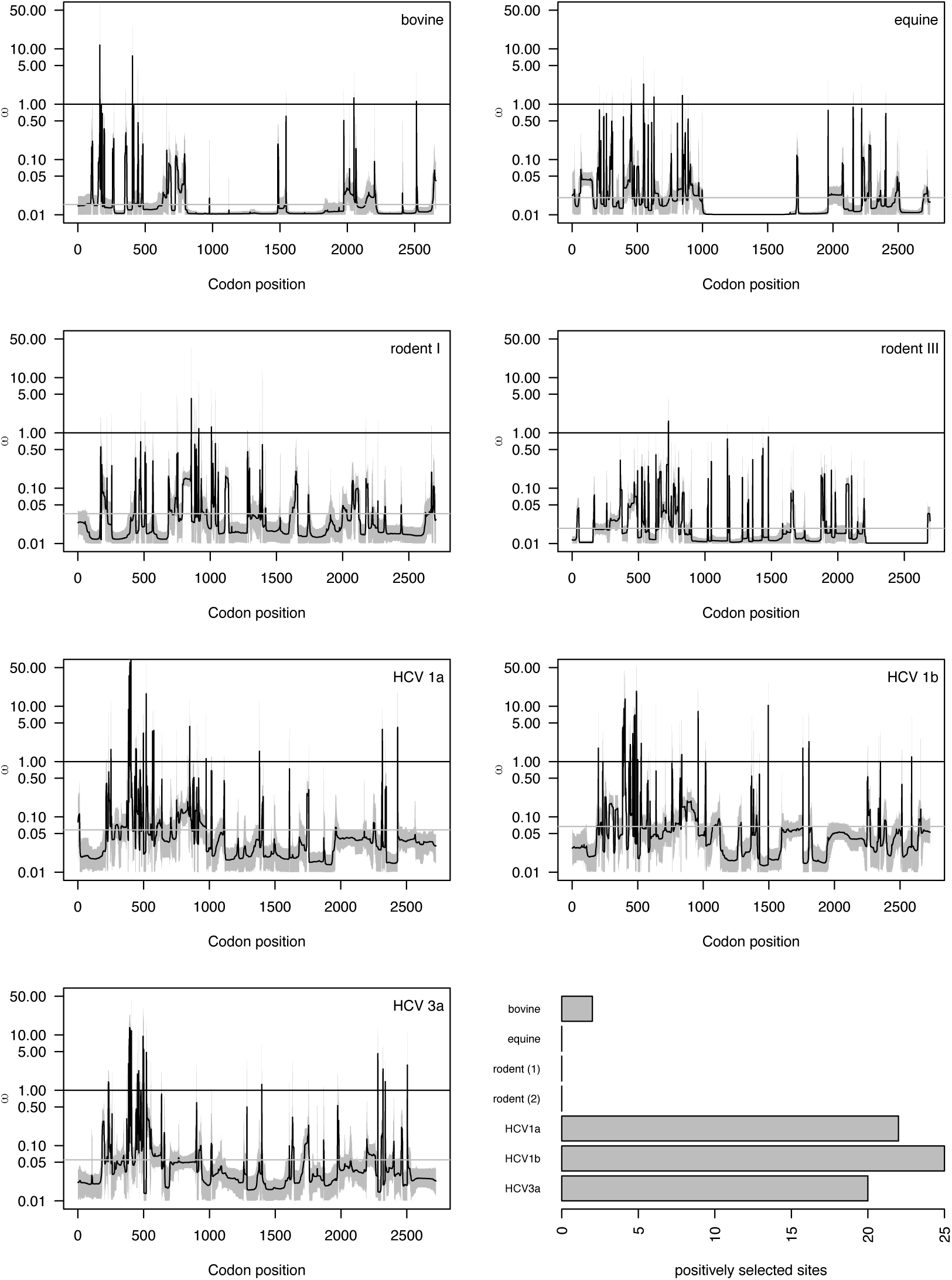
Site-specific variation of selection pressure in hepaciviruses. Estimates of ω in different animal hosts of hepaciviruses using omegaMap (Wilson and McVean, 2006). Equine and rodent hepaciviruses show no positively selected sites across their genome. For bovine hepaciviruses only two sites evolve under positive selection. HCV genotypes 1a, 1b and 3a indicate statistically significant positive selection pressure in 22, 25 and 20 sites respectively.

Because hepacivirus evolutionary rates have only been estimated for HCV, we here explore how informative current sampling in other host lineages is about the tempo of hepacivirus evolution while accounting for recombination (cfr. Methods). Using a recently developed test that compares the fit of a model that incorporates sampling time (the ‘dated tip’ model) to a model that assumes sampling time is uninformative (all sequences are sampled contemporeanously) (Duchene *et al*., 2019), we provide formal evidence that there is no sufficient temporal signal in bovine, equine and the two rodent lineages tested (Tab. 1) In the different HCV data sets on the other hand, temporal signal is consistently supported by a log Bayes factor support *>* 3. This discrepancy is likely explained by the difference in sampling time ranges for most data sets. Although the equine lineage has the broadest sampling time range, it is highly unbalanced with three closely related donkey viruses sampled in 1979 and all other viruses sampled between 2011 and 2016 (Supplementary Figure 6).

| Lineage | # of dated sequences | Sampling time range | BETS ln Bayes factor |
| --- | --- | --- | --- |
|  |  |  | (dated vs contemporaneous) |
| Bovine | 14 | 2013 - 2017 | 0.26 |
| Equine | 22 | 1979 - 2016 | -5.6 |
| Rodent I | 11 | 2010 - 2013 | 0.1 |
| Rodent III | 36 | 2010 - 2013 | -0.28 |
| HCV1a | 35 | 1997 - 2014 | 6.19 |
| HCV1b | 34 | 1990 - 2015 | 37.08 |
| HCV3a | 35 | 2002 - 2014 | 5.11 |

## Discussion

In this study, we performed the most comprehensive screening for hepaciviruses in African small mammals with a strong focus on rodents. We detect hepaciviruses in 29 animal species that had not been screened and found to carry hepaciviruses before, and therefore, considerably expand the RHV host spectrum. In line with previous research (Drexler *et al*., 2013; Kapoor *et al*., 2013; Van Nguyen *et al*., 2018), we demonstrate that rodents constitute an important source of hepaciviruses and that the evolutionary history of those pathogens has been largely shaped by host switching events. Finally, we identify a high rate of hepacivirus co-infections among *Lophuromys* rodents and conduct evolutionary analyses within specific host lineages.

While bats have received much attention as important pathogen reservoirs of infectious diseases, equally large-scale surveillance efforts have focused on rodents and, to a lesser extent, other small mammals. Rodents are generally considered as major transmitters of zoonoses carrying more than 66 pathogens that have crossed species barriers and infected humans (Han *et al*., 2015; Woolhouse and Gaunt, 2007). The number of virus lineages carried by vertebrate orders appears correlated mainly with the number of species present in these orders (Mollentze and Streicker, 2020). Therefore, species rich orders such as bats and rodents can be expected to host a higher number of viruses with zoonotic potential (Mollentze and Streicker, 2020). Identifying which animal was the source of hepaciviruses transmission to humans is of utmost importance, since this information can be used to unravel the mechanisms of HCV epidemic emergence and spread (Hartlage *et al*., 2016; Pybus and Thézé, 2016) as well as to assess whether we are still at risk of other emergence events.

In our sampling, 1.86% was positive for HCV homologues, a prevalence consistent with previous rodent screening efforts performed by Drexler *et al*. (2013). That study detected hepaciviruses in 1.8% of the *Myodes glareolus* population tested and a prevalence of 1.9% in *Rhabdomys pumilio* species. Although we detected three localities with a considerably higher prevalence of RHVs, these location-specific hotspots could also be correlated with variation in sample preservation, as they reflect distinct sampling sessions and therefore distinct ways in which the samples were preserved. Our 56 novel genomes represent new virus lineages and complement earlier efforts to uncover the diversity of RHVs. (Drexler *et al*., 2013; Kapoor *et al*., 2013; Van Nguyen *et al*., 2018; Wu *et al*., 2018).

A hepacivirus nomenclature has been proposed that consists of 14 species: *Hepacivirus A - N* (Smith *et al*., 2016); a classification made based on the amino acid divergence in distinct parts of the hepacivirus polyprotein. As more information accumulates on the genetic diversity of those pathogens, it becomes extremely challenging to define specific criteria for their classification (Simmonds *et al*., 2017). The current demarcation criteria do not adequately accommodate the high genetic diversity of hepaciviruses because they lead to discrepancies in the number of assigned species, as is demonstrated in our analysis (Supplementary Figure 4). This calls into question the current demarcation criteria and leaves hepacivirus classification as an open issue for further discussion.

To date, hepacivirus homologues in horses (EHV) and dogs (CHV) remain the closest relatives of HCV. Nevertheless, there is substantial genetic divergence between the equine/canine lineage and HCV, which casts considerable doubt on the hypothesis that Hepatitis C virus may jumped directly from horses to humans (Pybus and Thézé, 2016; Scheel *et al*., 2015). As Pybus and Gray (2013) and Hartlage *et al*. (2016) argue, there are currently two plausible scenarios for the origin of HCV. On the one hand, one can speculate that a single spillover event from a zoonotic reservoir established the infection in the human population. This ancestral HCV virus subsequently evolved and diversified in humans, resulting in the current genomic variability among genotypes. This is in agreement with a monophyletic HCV cluster, but it seems implausible that this resulted from the introduction of an ancestral EHV/CHV virus because of its substantial genetic divergence as a sister lineage and its relatively shallow diversity. On the other hand, it is conceivable that hepaciviruses have jumped to humans on multiple independent occasions and subsequently gave rise to the highly diverse HCV genotypes. Although there is currently no data to support this hypothesis, this may be attributed to sparse sampling of potential hosts (e.g. additional primate species or lagomorphs) and geographic gaps in surveillance. Based on the currently available sampling, primates, rodents and bats accommodate the highest hepacivirus genetic diversity among the mammalian hosts. Therefore, it is plausible that they represented an ancestral zoonotic source for transmission to other mammals (including humans) independently, of whether they crossed the species barrier once or multiple times (Moreira-Soto *et al*., 2020). While it is possible that viral lineages more closely related to HCV have gone extinct in their specific hosts, primates, rodents and bats deserve further attention as potential reservoirs. Surveillance in other hosts is also critical for mapping the broad host range of these viruses and to study their ecology and evolution.

Despite many host switches, there is still some non-random clustering of hepaciviruses based on rodent taxon. All *Lophuromys* hepacivirus clades form monophyletic groups exclusive to *Lophuromys* species, despite the fact that they have been sampled thousands of kilometers away. Furthermore, hepaciviruses sampled from other rodent taxa much closer geographically to some of these brush furred rat samples belong to different hepaci-lineages. This strongly supports that the hepacivirus evolutionary history has, at least to some extent, been driven by confinement to specific rodent taxa. These observations fit with an ancient evolutionary history constrained by the genetic background of the hosts. Furthermore, it is clear that early on in the evolution several lineages wound up in the same rodent taxa and have evolved in parallel with other hepacivirus lineages in the same rodent taxa.

Characterizing hepaciviruses in rodents may also prove relevant for HCV vaccine research. While treatment with direct-acting antiviral compounds has considerably advanced the past few years, a prophylactic vaccine is still lacking due to the absence of an *in vivo* model to study virus-host interactions within the liver. This has been an active field of research that made considerable progress in the development of surrogate rat models of chronic HCV infection (Billerbeck *et al*., 2017; Hartlage *et al*., 2019). Our work may motivate further biological characterization of RHVs, and the evidence of hepacivirus co-infections in specific rodents may have immunological implications to consider.

Remarkably, we only observed co-infections in a particular genus of the Muridae family, the brush furred rats, even though various genera from the same family have been found to carry hepaciviruses. To date, relatively little is known about the behavioural ecology of the four *Lophuromys* species that harboured co-infections. These four rodent species are phylogenetically closely related (they belong to the so-called *L. flavopunctatus* complex) (Verheyen *et al*., 2002, 2007) and they diverged in Pleistocene in different forest fragments (Komarova et al., submitted). Most of the species from this complex are endemic in relatively humid habitats of central and east Africa (Sabuni *et al*., 2018; Van de Perre *et al*., 2019a). Although *Lophuromys* tend to be solitary and show antagonist behavior to conspecifics, they can sometimes live in very high population densities. In captivity they may fight until death (Kingdon et al., 2013) and if such conflicts occur in natural circumstances, it may represent a mode of transmission that could help to explain the elevated RHV detection and co-infection rate. Furthermore, these rodents can be occasionally infested with blood-sucking fleas depending on the location and the specific flea index. Flea sharing between sympatric species of rodents has been previously described (Laudisoit *et al*., 2009) and could possibly support a scenario of RHV mechanical transmission.

Interestingly, a co-infection of two divergent paramyxovirus lineages was also found in a *Lophuromys* specimen, and in no other paramyxovirus host (Vanmechelen *et al*., 2018). Whether the apparent propensity of brush furred rats to be co-infected with multiple lineages of the same virus family is due to a common physiological background of the closely related species that enhances their susceptibility or tolerance of multiple hepacivirus/paramyxovirus infections, or because of behavioral characteristics that increases the transmission probabilities, is still unknown. Further research is needed into the heterogeneous viral detection and co-infection rate in rodents and how those are shaped by specific transmission dynamics.

Prior to this study, hepacivirus co-infections have to our knowledge only been identified for HCV in humans (Blackard and Sherman, 2007; Morel *et al*., 2010). While preliminary, the fact that only one RHV strain from the rodent I cluster co-circulates with multiple strains from the rodent III lineage appears to support a pattern of dominance of rodent I viruses over the rodent III variants. Whether this dominant strain hypothesis denotes any significant mode of infection needs further biological testing, ideally using an experimental mouse model. This may help to define critical elements of hepacivirus persistence, especially in the presence of multiple RHV co-circulating strains. Knowledge of the frequent establishment of co-infections within the *Lophuromys* mice may open new experimental horizons and offer more insights into the pathogenesis and immunity against hepaciviruses.

Although the blood-borne transmission route is well documented for HCV, other potential modes of infection for hepaciviruses are poorly characterized. Recently, divergent hepaciviruses were also discovered in two non-vertebrate hosts including a *Culex annulirostris* mosquito (Williams *et al*., 2020) and an *Ixodes holocyclus* tick species (Harvey *et al*., 2019). Phylogenetic analysis has grouped the mosquito hepacivirus with viruses present in birds (Williams *et al*., 2020), while the tick hepacivirus clusters within the rodent I lineage (Harvey *et al*., 2019). Both analyses, however, provided strong evidence that those invertebrate hosts were feeding on a bird species and a long-nose bandicoot, respectively. Whether mosquitoes or ticks act as intermediate hosts or vectors of hepacivirus transmission is currently speculative and additional surveillance is required to verify this infection route.

As part of our evolutionary analyses, we focused on recombination as an important driver of genetic diversity. Recombination is relatively uncommon in the extensively studied HCV population (González-Candelas *et al*., 2011; Karchava *et al*., 2015; Raghwani *et al*., 2012; Susser *et al*., 2017) and while some evidence for interspecific hepacivirus recombination has been found (Thézé *et al*., 2015), the authors indicated that a clear interpretation of this result is hampered by high genetic divergence and undersampling. We focused on recombination within specific host lineages and detected significant signal in the bovine, equine and two rodent lineages. This implies that co-infections, for which we found evidence in specific rodent hosts, also occur in other animal hosts.

Using selection analyses that account for recombination, we estimate an overall negative selection pressure on the virus population in each host providing evidence for a process of evolution under predominantly purifying selection. However, this does not exclude the possibility of episodic molecular adaptation in the evolutionary history of these viruses for example following a cross-species transmission to a new host. Unfortunately, the extensive interspecific genetic divergence hampers uncovering such events in codon sequences. We consistently identify a similar fraction of positively selected sites in three HCV genotype data sets, in particular in the E1/E2 region, while such sites are rare or absent in hepaciviruses in animal hosts. It is therefore interesting to speculate that differences in immune responses may, together with differences in transmission intensity, underly some variability in hepacivirus co-infections and hence also differences in recombination rates.

Although the hepacivirus discovery phase is still ongoing, the tremendous advances in genomics technologies allow us to start characterizing the evolutionary dynamics of these viruses beyond what is known from HCV research. For rapidly evolving RNA viruses, evolutionary rates can be estimated based on the sequence divergence that accumulates between genome samples obtained at different time points. We demonstrate that the current sampling time range is insufficient for calibrating a hepacivirus molecular clock in the different animal hosts. This calls for further characterization of hepacivirus genomes, both from old samples as well as from more recent samples, in order to capture sufficient temporal signal. This will provide the ability to estimate divergence times in the hepacivirus evolutionary history as well as to study the biological factors underlying evolutionary rate variation.

In conclusion, we show that viral genomic studies provide important information about the diversity, transmission history within and among different hosts, and evolutionary dynamics of hepaciviruses. We hope that screening efforts guided by ecologists will not only target wild animals but also commensal species that live in close proximity to residential areas. Characterizing possible routes of transmission among those hosts and/or between different hosts may prove particularly interesting as it may provide insights into the ecological barriers for viruses at the rodent-human interface. Hopefully, the expanding hepacivirus diversity will motivate further biological studies aimed at elucidating hepacivirus transmission routes and modes of infection.

## Acknowledgements

The research leading to these results has received funding from the European Research Council under the European Union’s Horizon 2020 research and innovation programme (grant agreement no. 725422-ReservoirDOCS). The Artic Network receives funding from the Wellcome Trust through project 206298/Z/17/Z. BV (grant number 12U7118N) and SL were supported by post-doctoral grants of the FWO (Fonds Wetenschappelijk Onderzoek – Vlaanderen, grant number for BV). PL acknowledges support by the Research Foundation – Flanders (‘Fonds voor Wetenschappelijk Onderzoek – Vlaanderen’, G066215N, G0D5117N and G0B9317N). FVdP was supported by a Ph.D. fellowship from the Research Foundation-Flanders.

Most samples provided by the Czech team were collected and processed within the projects of the Czech Science Foundation, no. P506/10/0983 and P502/11/J070. The fieldwork in the DR Congo was supported by the COBIMFO Project (Congo Basin integrated monitoring for forest carbon mitigation and biodiversity; contract no. SD/AR/01A) and was funded by the Belgian Science Policy Office (Belspo). The research permits and authorizations were granted by the University of Kisangani Biodiversity Surveillance Center (CSB [Centre de Surveillance de la Biodiversité] – in French) which has the scientific authority to do so. All “movement of personal” (called “ordre de Mission” were signed and approved by the local authorities at each field site. Material transfer agreements (MTA) were issued by the University of Kisangani, Biodiversity Surveillance Center (CSB [Centre de Surveillance de la Biodiversité]). None of the animal samples are listed on the CITES Red List of protected species nor on the national list of protected fauna of the DRC; hence no CITES permit was needed prior to shipping.

Finally, we gratefully acknowledge Zafeiro Zisi from Rega Institute, KU Leuven for her valuable technical assistance and Andrea Rasche from Charite - Universitatsmedizin Berlin for useful discussions and suggestions during the initiation of this project. The authors would also like to thank Viroscan 3D for their contribution and excellent technical service in this project.

## Notes

### Competing Interest Statement

The authors have declared no competing interest.

