## Supplementary Information for "Unravelling the evolutionary relationships of hepaciviruses within and across rodent hosts"

Magda Bletsa[1], Bram Vrancken[1], Sophie Gryseels[1,2], Ine Boonen[1], Antonios Fikatas[1], Yiqiao Li[1], Anne Laudisoit[3], Sebastian Lequime[1], Josef Bryja[4], Rhodes Makundi[5], Yonas Meheretu[6], Benjamin Dudu Akaibe[7], Sylvestre Gambalemoke Mbalitini[7], Frederik Van de Perre[8], Natalie Van Houtte[8], Jana Těšíková[4,9], Elke Wollants[1], Marc Van Ranst[1], Jan Felix Drexler[10,11], Erik Verheyen[8,12], Herwig Leirs[8], Joelle Gouy de Bellocq[4] and Philippe Lemey[1]

[1]*Department of Microbiology, Immunology and Transplantation, Rega Institute, KU Leuven – University of Leuven, Leuven, Belgium*

[2]*Department of Ecology and Evolutionary Biology, University of Arizona, Tucson, USA*

[3]*EcoHealth Alliance, New York, USA*

[4]*Institute of Vertebrate Biology of the Czech Academy of Sciences, Brno, Czech Republic*

[5]*Pest Management Center – Sokoine University of Agriculture, Morogoro, Tanzania*

[6]*Department of Biology and Institute of Mountain Research & Development, Mekelle University, Mekelle, Ethiopia*

[7]*Department of Ecology and Animal Resource Management, Faculty of Science, Biodiversity Monitoring Center, University of Kisangani, Kisangani, Democratic Republic of the Congo*

[8]*Department of Biology, Evolutionary Ecology Group, University of Antwerp, Antwerp, Belgium*

[9]*Charite - Universitatsmedizin Berlin, Berlin, Germany*

[10]*German Center for Infection Research (DZIF), Germany and*

[11]*OD Taxonomy and Phylogeny - Royal Belgian Institute of Natural Sciences, Brussels, Belgium*

(Dated: October 7, 2020)

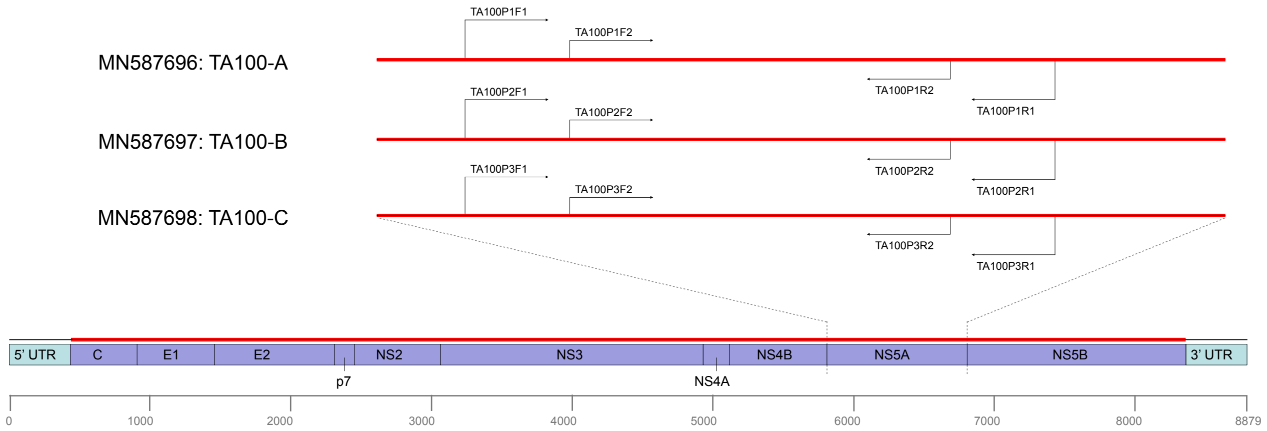


FIG. 1 **Schematic overview of the primers used in our co-infection validation assay.** Purple and light blue boxes correspond to the full genome organization of rodent hepaciviruses. Numbering of positions is relative to GenBank accession number NC_021153. Primers P1, P2 and P3 denote the distinct PCR assays designed on the three divergent hepaciviruses of specimen TA100. F1 and R1 represent the outer forward and reverse primers, respectively. Accordingly, F2 and R2 refer to the inner forward and reverse primers. For sample TA100 for example, we will have primer pair TA100P1F1 – TA100P1R1, which represents the outer primer pair designed specifically to amplify a fragment of the NS5A region of TA100-A (accession number: MN587696).

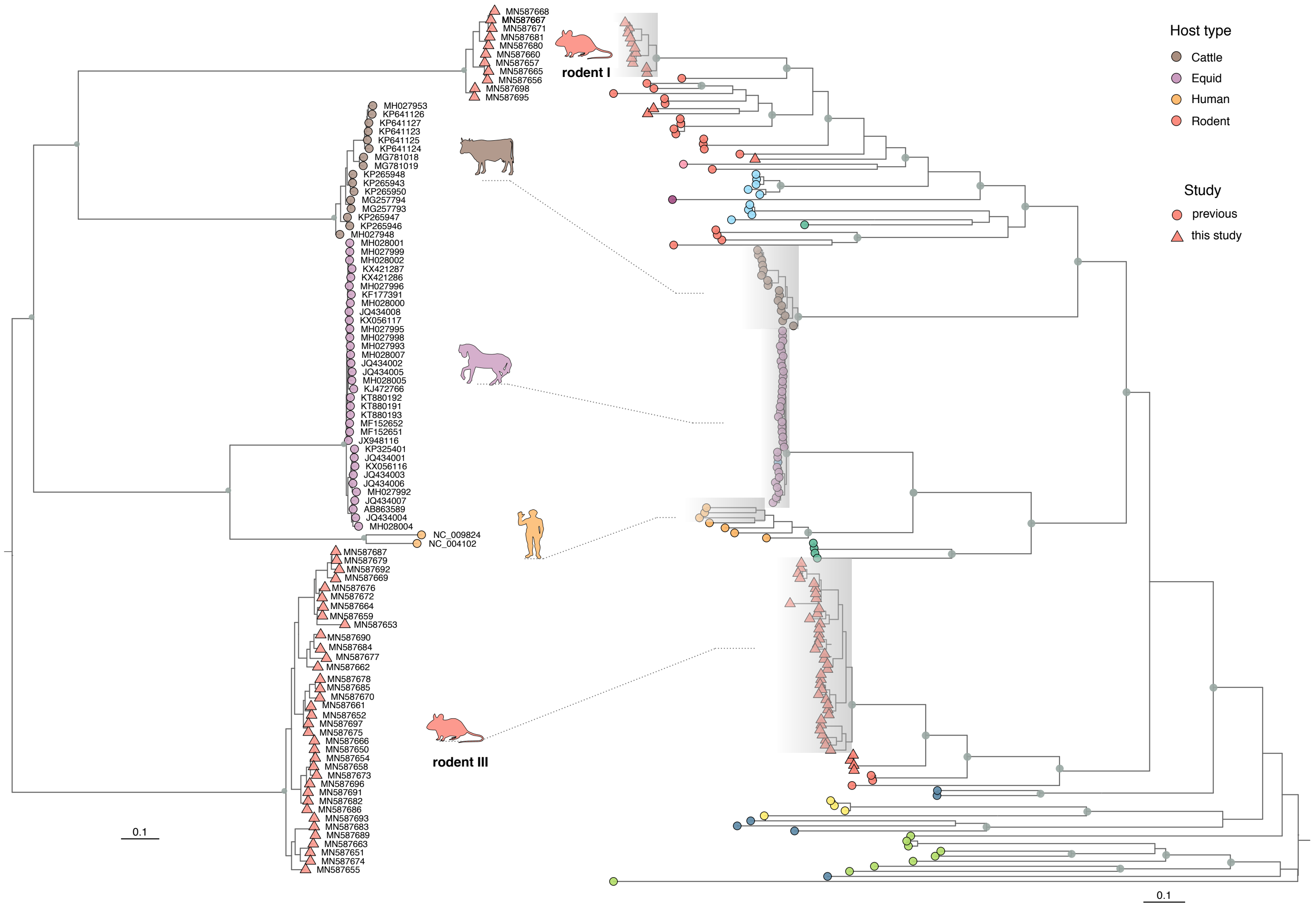


FIG. 2 **Host-specific lineages used in the recombination and selection analyses.** ML subtree of the genome-wide phylogenetic reconstruction depicted in main Figure 4. Five host-specific lineages were analyzed (left phylogeny) that showed limited genetic diversity compared to all the other hepacivirus lineages (right phylogeny). The colouring scheme follows the host assignment from main Figure 4.

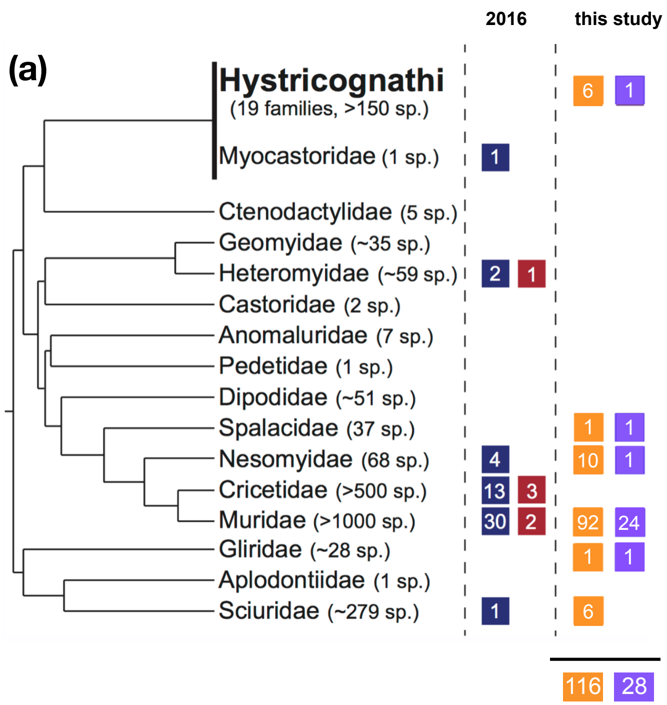


(b)

| Country | Samples | Positives (%) |
|---|---|---|
| DRC | 894 | 16 (1.79) |
| Ethiopia | 426 | 4 (0.94) |
| Kenya | 532 | 8 (1.50) |
| Madagascar | 30 | 0 |
| Mozambique | 399 | 10 (2.51) |
| Tanzania | 1,798 | 41 (2.28) |
| Zambia | 224 | 1 (0.45) |
| | **4,303** | **80 (1.86)** |

FIG. 3 **Sampling and screening summary of hepaciviruses in rodents. (a)** The phylogeny of the extant rodent families reproduced from (Pybus and Thézé, 2016) and adapted to include our current rodent sampling. The large rodent suborder Hystricognathi (in bold) has been collapsed into a single lineage for clarity. Boxes next to each family name demonstrate the estimated number of species that has been previously screened (blue box) and those found to harbour hepaciviruses in earlier studies (red box). The number of potential rodent species in our sampling is shown in orange boxes and the number of species within which we have detected hepaciviruses is indicated in the purple boxes. **(b)** The capture efforts were performed at multiple localities of seven African countries: the Democratic Republic of the Congo (DRC), Ethiopia, Kenya, Madagascar, Mozambique, Tanzania and Zambia. The number of sampled individuals is shown next to each country, along with the number of specimens that were found to be positive for hepaciviruses. In brackets we summarize the percentage of positives detected within the different countries.

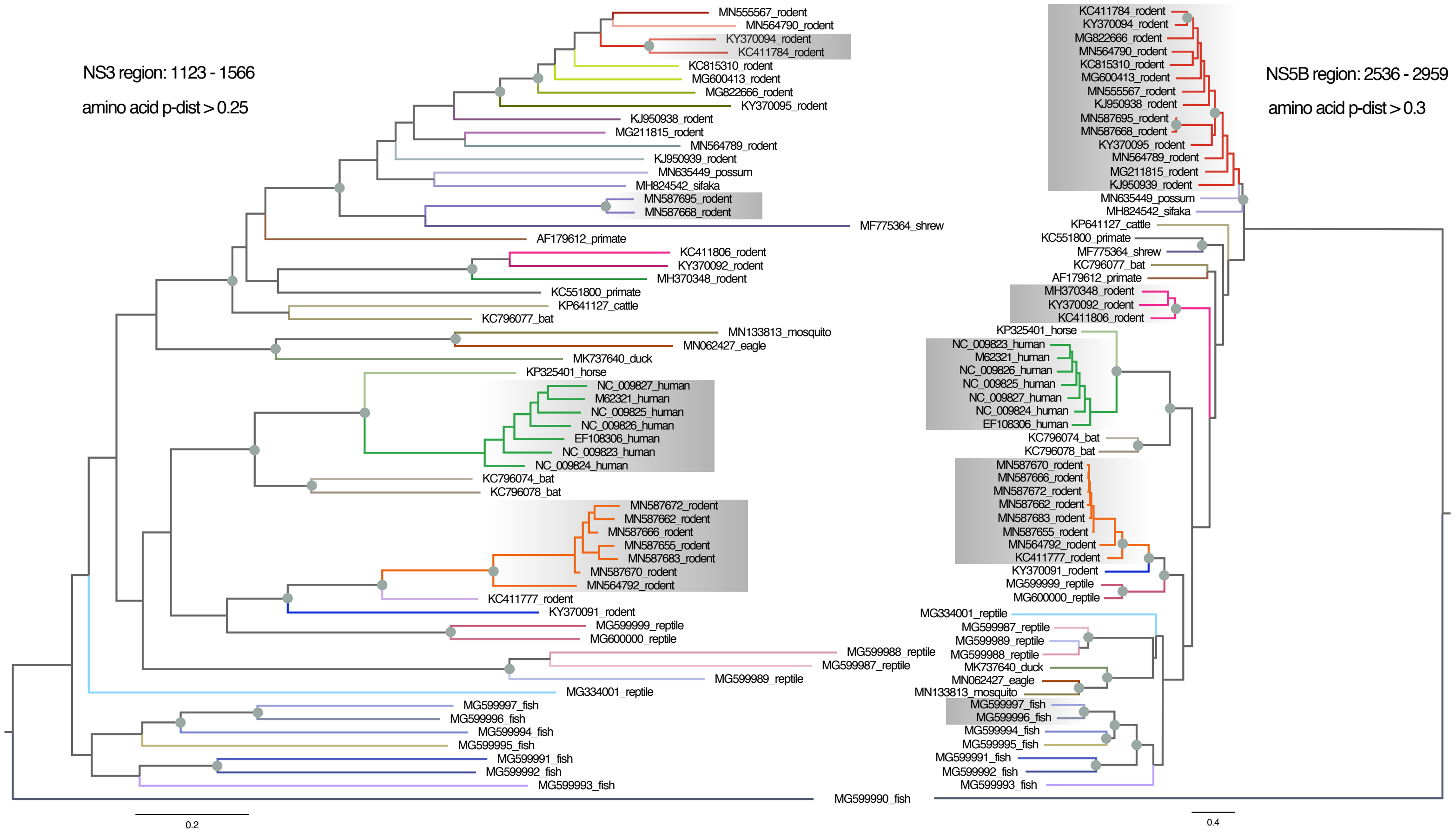


FIG. 4 ***Hepaciviruses* species classification following the taxonomic proposal by Smith *et al.* (2016).** For this analysis we used a reduced data set of 60 genomes differing over their complete coding sequence by amino acid p-distances greater than 0.1 (more details in Supplementary Tables 6 and 7). ML trees were generated for amino acid regions 1123 - 1566 and 2536 - 2959 (reference sequence: M62321). Demarcation between species occurs when amino acid p-distances in region 1123 - 1566 are greater than 0.25 and greater than 0.3 in region 2536 - 2959. Grey circles denote internal nodes with bootstrap support ≥70, while grey shaded areas indicate virus species with more than one representatives.

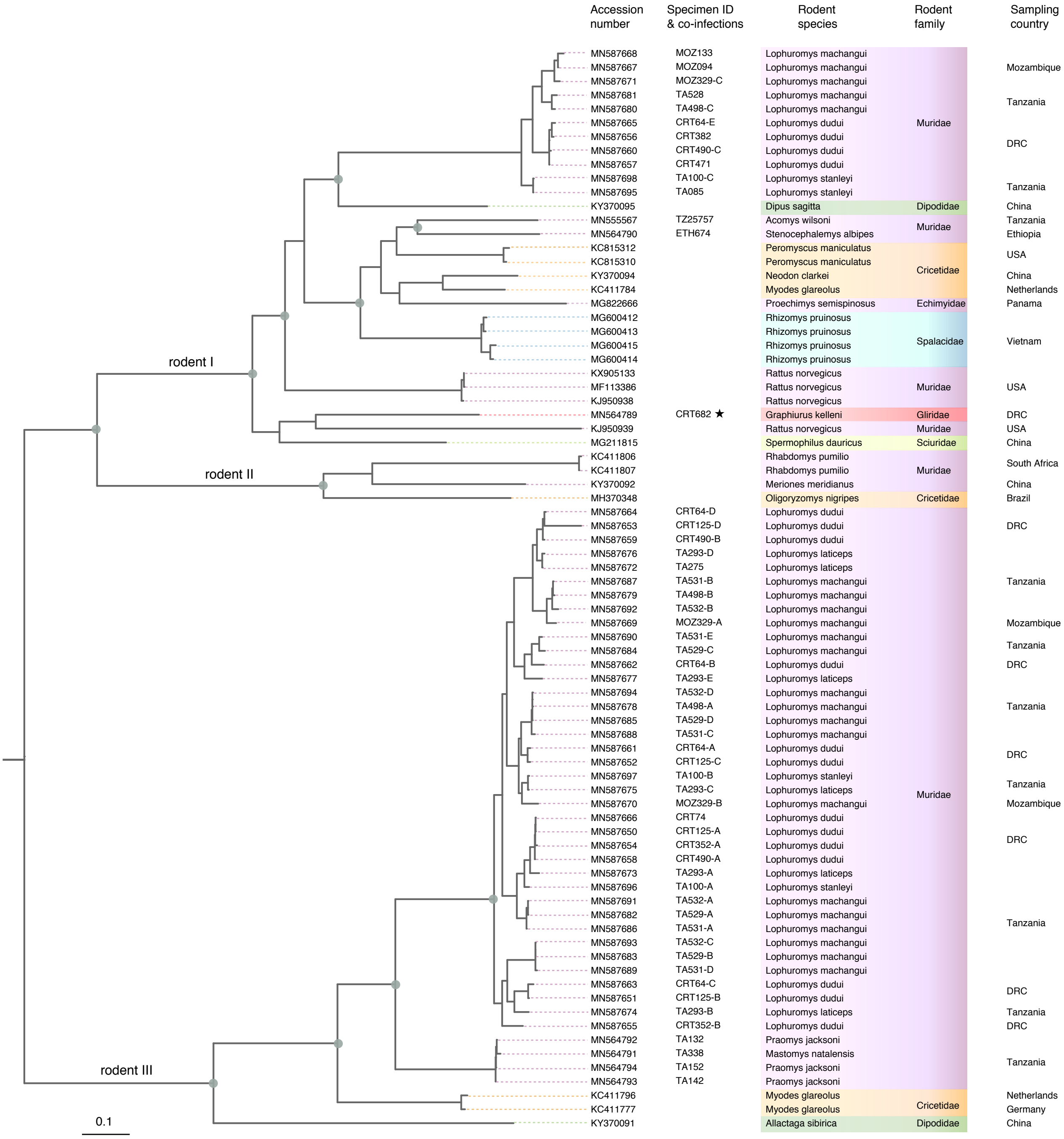


FIG. 5 **Phylogenetic reconstruction of rodent hepaciviruses.** ML tree of all available ($n = 22$) and novel ($n = 56$) hepacivirus genomes. Columns next to the phylogeny indicate GenBank accession numbers, specimen IDs and their associated RHV strains, the exact rodent species and family and the country of origin. Grey circles indicate internal nodes with bootstrap support $\geq$70. The star denotes isolate CRT682, that originated in a *Graphiurus kelleni* individual collected in the DRC.

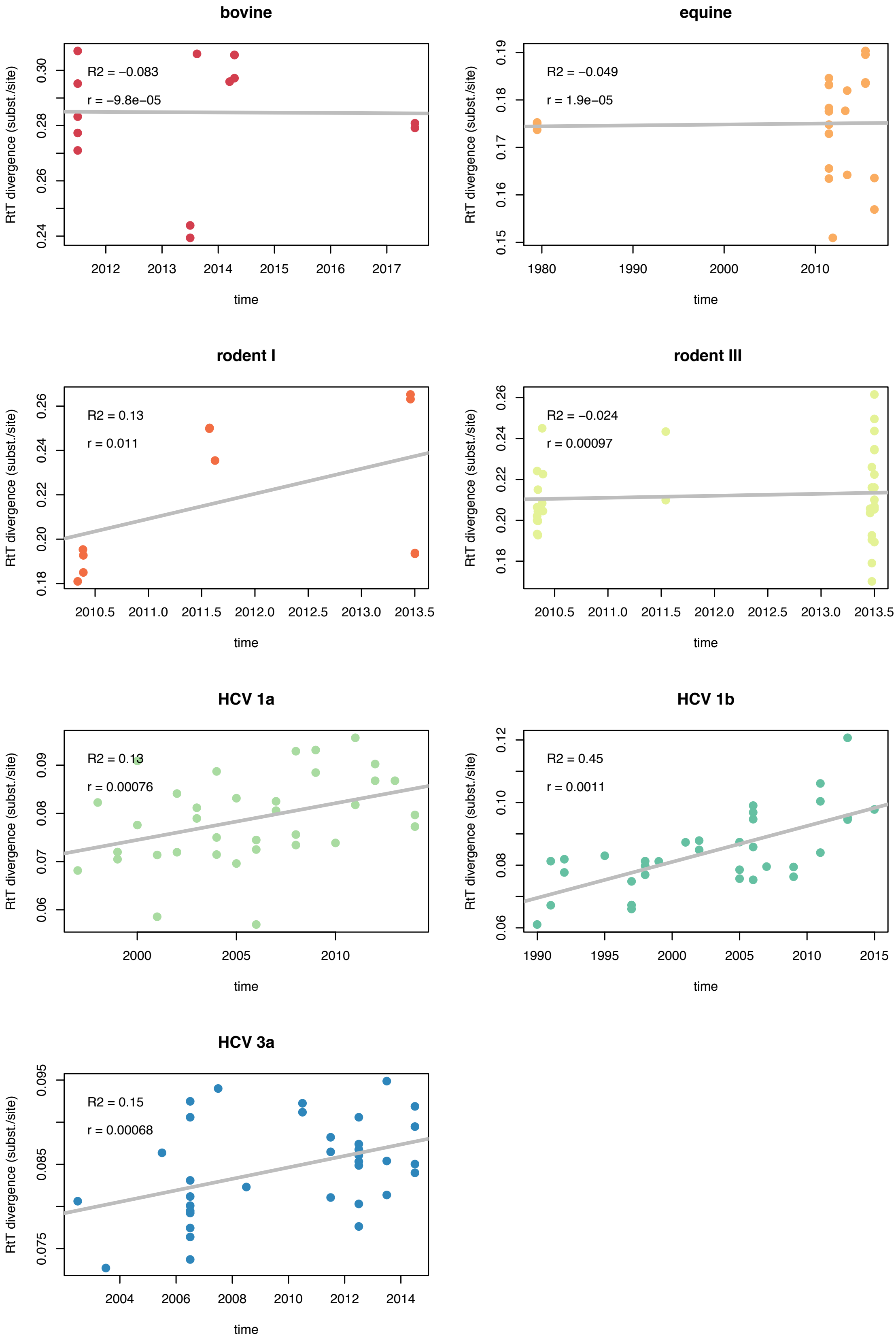


FIG. 6 **Temporal signal analysis of hepaciviruses.** For specific host lineages with limited diversity we show root-to-tip divergence as a function of sampling year. Co-efficients of determination (R2) and slope estimates are indicated in the upper left corner of the regression plots. Data points are coloured according to the specific host lineages: bovine (red), equine (orange), rodent I (coral), rodent III (lime green), HCV1a (light green), HCV1b (green), HCV3a (blue).

- Supplementary Figure 1:

  Outer and inner primer pairs were designed targeting the most variable region of the rodent hepacivirus genome, as illustrated in Suppl. Figure 1.

- Supplementary Figure 2:

  To test for recombination, we selected lineage-specific data sets with limited and shallow diversity. Suppl. Figure 2 demonstrates the lineages used in our recombination and selection analyses, while the species composition of these viral genomes is detailed here:

  – Rodent hepaciviruses from four *Lophuromys* species were used: *L. dudui*, *L. laticeps*, *L. machangui* and *L. stanleyi*.

  – Equine hepaciviruses from two *Equus* species were used: *E. ferus* and *E. asinus*.

  – Bovine hepaciviruses included sequences only from the *Bos taurus* species.

  – Hepatitis C virus datasets were obviously restricted to the *Homo sapiens* species.

  A substantial number of lineage-specific recombinants were identified in rodents, with the highest proportion in strains circulating in the rodent III lineage. Furthermore, three hepaciviruses present in Ghanaian cattle (accession numbers: KP2655943, KP2655948, KP2655950) appeared to share a recombinant part between positions 4953 and 5064 with an ancestor of the German cattle hepacivirus MH027948. For the equine lineage, recombination events were detected between strains circulating in Europe (and the UK) with those circulating in the USA. The best supported recombination breakpoint was located between positions 6472 and 7608.

  We assume that the virus population dynamics remained the same within the different host genera. It is, therefore, conceivable that comparison of selective pressure acting on closely related viruses holds a biological explanation even when pathogens from congeneric species are being tested.

- Supplementary Figure 3:

  Panel a demonstrates the species range of our current rodent sampling, while panel b summarizes our screening results per African country along with the composition of our sample collection. For a detailed list of all hepaci-positive specimens we refer to Supplementary Table 5.

- Supplementary Figure 4:

  In an attempt to position our novel sequences within the tremendous heterogeneity of hepaciviruses, we followed the classification proposal by Smith *et al.* (2016). For this analysis we used a reduced data set of 60 genomes differing over their complete coding sequence by amino acid $p$-distances greater than 0.1 (Supplementary Table 6). This assignment resulted in 46 *Hepacivirus* species for the amino acid region 1123 - 1566 and in 31 *Hepacivirus* species for the amino acid region 2536 - 2959 (Supplementary Figure 2). Only 18 genomes were classified in groups of more than one representatives in the NS3 region, while 34 genomes were assigned in lineages with multiple hepaciviruses in the NS5B fragment. Although the tree topology remains congruent between those regions, the number of assigned *Hepacivirus* species differs dramatically. The current demarcation criteria not only fail to classify viruses from host-specific lineages into one single species, but also they provide erratic assignments between the proposed conserved positions. For example, major inconsistencies appear to exist in the rodent I cluster, which is divided into 15 distinct *Hepacivirus* species for amino acid region 1123 - 1566 compared to the two well-defined species for the other region.

- Supplementary Figure 5:

  Rodent hepacivirus phylogeny annotated with GenBank accession numbers, specimen IDs and their associated RHV strains, the exact rodent species and family and the country of origin.

- Supplementary Figure 6:

  Root-to-tip divergence of host-specific lineages as a function of their sampling year.

**Supplementary table S1:** Complete list of specimens that have been screened for the presence of hepaciviruses.

| Order | Family | Species | No. of samples | No. of positives | Sampling location | Sampling date |
|---|---|---|---|---|---|---|
| Afrosoricida | Chrysochloridae | *Chrysochloris stuhlmanni* | 2 | | CD | 2010 |
| | **Subtotal** | **1 species** | **2 samples** | **0 positives** | | |
| Carnivora | Herpestidae | *Atilax paludinosus* | 1 | | TZ | 2007 |
| | Viverridae | *Genetta angolensis* | 1 | | TZ | 2011 |
| | **Subtotal** | **2 species** | **2 samples** | **0 positives** | | |
| Chiroptera | Molossidae | *Chaerephon pumilus* | 3 | | ZM/TZ | 2010/2013 |
| | | *Mops condylurus* | 4 | | MZ | 2011 |
| | Pteropodidae | *Epomophorus gambianus* | 3 | | ZM | 2010 |
| | | *Epomophorus labiatus* | 8 | | MZ | 2011 |
| | | *Lissonycteris goliath* | 1 | | MZ | 2011 |
| | | *Rousettus aegyptiacus* | 3 | | KE | 2010 |
| | Rhinolophidae | *Rhinolphus hilebrandtii* | 2 | | MZ | 2011 |
| | Vespertilionidae | *Chalinolobus variegatus* | 1 | | ZM | 2010 |
| | | *Glauconycteris atra* | 6 | 2 | TZ | 2013 |
| | | *Pipistrellus* sp. | 5 | | ZM/KE/MZ | 2010/2010/2011 |
| | **Subtotal** | **10 species** | **36 samples** | **2 positives** | | |
| Eulipotyphla | Erinaceidae | *Atelerix* sp. | 1 | | TZ | 2013 |
| | Soricidae | *Crocidura caliginea* | 15 | | CD | 2012,2013 |
| | | *Crocidura* cf. *denti* | 34 | | CD | 2012,2013 |
| | | *Crocidura* cf. *flavescens* | 4 | | TZ | 2013 |
| | | *Crocidura* cf. *littoralis* | 18 | | CD | 2012,2013 |
| | | *Crocidura* cf. *ludia* | 68 | | CD | 2012,2013 |
| | | *Crocidura dolichura* | 10 | | CD | 2012,2013 |
| | | *Crocidura fumosa* | 4 | | KE | 2010 |
| | | *Crocidura hildegardeae* | 5 | | TZ | 2013 |
| | | *Crocidura hirta* | 22 | | ZM/MZ | 2009,2010/2011 |
| | | *Crocidura lamottei-parvipes group* | 3 | | TZ | 2013 |
| | | *Crocidura luna* | 9 | | ZM/TZ | 2010/2013 |
| | | *Crocidura montis* | 22 | | TZ | 2013 |
| | | *Crocidura olivieri* | 46 | | ZM/ET/KE/CD/TZ | 2009/2010/2010/2010,2012,2013/2013 |
| | | *Crocidura silacea-mariquensis* | 1 | | ZM | 2010 |
| | | *Crocidura* sp. | 149 | | ZM/MZ/TZ/CD | 2010/2011/2007-2013/2007,2010-2013 |
| | | *Crocidura* sp. *Chipata* | 5 | | ZM | 2010 |
| | | *Crocidura* sp. *Kaoma* | 1 | | ZM | 2010 |
| | | *Crocidura turba* | 8 | | ZM/KE/TZ | 2009/2010/2013 |
| | | *Paracrocidura schoutedeni* | 5 | | CD | 2012,2013 |
| | | *Scutisorex congicus* | 1 | | CD | 2013 |
| | | *Scutisorex* sp. | 5 | | CD | 2012 |
| | | *Suncus* sp. | 9 | | CD | 2013 |
| | | *Surdisorex polulus* | 1 | | KE | 2010 |
| | | *Sylvisorex akaibei* | 9 | | CD | 2013 |
| | | *Sylvisorex* sp. | 4 | | TZ/CD | 2006/2013 |
| | **Subtotal** | **26 species** | **459 samples** | **0 positives** | | |
| Hyracoidea | Procaviidae | *Dendrohyrax dorsalis* | 1 | | CD | 2012 |
| | **Subtotal** | **1 species** | **1 sample** | **0 positives** | | |

| | | | | | | |
|---|---|---|---|---|---|---|
| Macroscelidae | Macroscelididae | *Elephantulus brachyrhynchus* | 6 | | ZM/KE/TZ | 2009/2010,2011/2013 |
| | | *Elephantulus rufescens* | 1 | | KE | 2010 |
| | | *Petrodromus tetradactylus* | 1 | | MZ | 2011 |
| | | *Petrodromus tordayi* | 6 | | CD | 2013 |
| | | *Rhynchocyon cirnei* | 1 | | MZ | 2011 |
| | **Subtotal** | **5 species** | **15 samples** | **0 positives** | | |
| Rodentia | Bathyergidae | *Fukomys anselli* | 3 | | ZM | 2010 |
| | | *Fukomys bocagei* | 1 | | ZM | 2010 |
| | | *Fukomys mechowii* | 4 | | ZM | 2009 |
| | | *Fukomys whytei* | 1 | | ZM | 2009 |
| | | *Heliophobius argenteocinereus* | 29 | 1 | TZ/ZM/KE/MZ | 2008/2010/2010/2011 |
| | Gliridae | *Graphiurus* sp. * | 25 | 2 | TZ/KE/CD/ZM/MZ | 2007,2008,2010,2013/2010/2010/2010/2011 |
| | Muridae | *Acomys* aff. *percivali* | 4 | | KE | 2010 |
| | | *Acomys ignitus* | 13 | | KE | 2010 |
| | | *Acomys kempi* | 21 | 2 | KE | 2010 |
| | | *Acomys muzei* | 17 | | ZM/TZ | 2009,2010/2013 |
| | | *Acomys ngurui* | 82 | | MZ/TZ | 2011/2009-2011 |
| | | *Acomys percivali* | 6 | | KE | 2010 |
| | | *Acomys* sp. | 20 | 1 | TZ | 2008-2010/2013 |
| | | *Acomys spinosissimus* | 31 | | MZ | 2011 |
| | | *Acomys wilsoni* | 8 | 2 | KE | 2010,2011 |
| | | *Aethomys chrysophilus* | 74 | | ZM/KE/MZ/TZ | 2009,2010/2010/2011/2007,2011,2013 |
| | | *Aethomys hindei* | 6 | | KE/MZ | 2010,2011/2011 |
| | | *Aethomys kaiseri* | 30 | | KE/TZ | 2010,2011/2013 |
| | | *Aethomys silindensis* | 7 | | MZ | 2011 |
| | | *Arvicanthis niloticus* | 25 | | ET | 2010 |
| | | *Arvicanthis nairobae* | 1 | | TZ | 2013 |
| | | *Arvicanthis neumanni* | 9 | | TZ | 2012 |
| | | *Arvicanthis* aff. *niloticus* | 2 | | KE | 2011 |
| | | *Arvicanthis somalicus* | 6 | | KE | 2010,2011 |
| | | *Arvicanthis* sp. | 55 | | CD/TZ | 2007/2007-2009 |
| | | *Deomys ferrugineus* | 1 | | TZ | 2013 |
| | | *Gerbilliscus* cf. *bayeri* | 2 | | KE | 2011 |
| | | *Gerbilliscus* cf. *cosensi* | 2 | | ZM/MZ | 2009/2011 |
| | | *Gerbilliscus* cf. *taborae* | 3 | | TZ | 2013 |
| | | *Gerbilliscus kempi* | 2 | | TZ | 2013 |
| | | *Gerbilliscus leucogaster* | 4 | | ZM/TZ | 2009/2013 |
| | | *Gerbilliscus nigricaudus* | 2 | | KE | 2011 |
| | | *Gerbilliscus phillipsi* | 1 | | KE | 2011 |
| | | *Gerbilliscus* sp. | 34 | 1 | TZ/MZ | 2007-2010,2012/2011 |
| | | *Gerbilliscus vicinus* | 14 | 2 | KE/TZ | 2010,2011/2013 |
| | | *Grammomys* cf. *gazellae* | 1 | | KE | 2010 |
| | | *Grammomys* cf. *kuru* | 3 | | CD/TZ | 2013 |
| | | *Grammomys ibeanus* | 1 | | TZ | 2007 |
| | | *Grammomys macmillani* | 16 | | KE | 2010,2011 |
| | | *Grammomys* sp. | 83 | 1 | CD/TZ | 2010/2006,2007,2009-2011 |
| | | *Grammomys surdaster* | 9 | | ZM/KE/TZ | 2009/2010/2013 |
| | | *Hybomys* cf. *univitattus* | 14 | | CD/TZ | 2013 |
| | | *Hybomys lunaris* | 1 | | CD | 2010 |
| | | *Hybomys* sp. | 40 | 1 | CD | 2010,2012,2013 |
| | | *Hylomyscus aeta* | 3 | | CD | 2012 |
| | | *Hylomyscus anselli* | 1 | | ZM | 2009 |

| | | | | |
|---|---|---|---|---|
| *Hylomyscus endorobae* | 5 | | KE | 2010 |
| *Hylomyscus* sp. | 45 | | CD | 2010-2013 |
| *Hylomyscus stella* | 26 | 1 | KE/CD/TZ | 2010/2012/2013 |
| *Lemniscomys rosalia* | 13 | | ZM/TZ | 2009/2007,2008,2013 |
| *Lemniscomys* sp. | 34 | 1 | TZ/CD/MZ | 2007,2008,2011/2010/2011 |
| *Lemniscomys striatus* | 50 | 1 | KE/TZ/CD | 2010,2011/2009,2013/2010 |
| *Lemniscomys zebra* | 43 | | TZ | 2008,2009,2011-2013 |
| *Lophuromys ansorgei* | 2 | | KE | 2010 |
| *Lophuromys dudui* | 10 | 8 | CD | 2012,2013 |
| *Lophuromys kilonzoi* | 33 | 5 | TZ | 2007 |
| *Lophuromys laticeps* | 13 | 3 | TZ | 2013 |
| *Lophuromys luteogaster* | 6 | | CD | 2012,2013 |
| *Lophuromys machangui* | 30 | 21 | MZ/TZ | 2011/2013 |
| *Lophuromys* sp. | 118 | 1 | CD/TZ | 2007,2010,2012,2013/2006,2009,2010,2011 |
| *Lophuromys stanleyi* | 31 | 4 | TZ | 2013 |
| *Lophuromys zena* | 60 | 2 | KE | 2010,2011 |
| *Malacomys longipes* | 25 | | ZM/CD | 2009/2010, 2012,2013 |
| *Mastomys awashensis* | 28 | 2 | ET | 2010/2012 |
| *Mastomys erythroleucus* | 53 | | KE/ET | 2011/2012 |
| *Mastomys kollmannspergeri* | 1 | | ET | 2012 |
| *Mastomys natalensis* | 643 | 1 | ZM/KE/MZ/ET/TZ | 2009,2010/2010,2011/2011/2012/2007-2009,2011-2013 |
| *Mastomys pernanus* | 4 | | KE | 2010 |
| *Mastomys* sp. | 159 | | CD/TZ | 2007,2010/2006,2007,2009-2011,2013 |
| *Micaelamys namaquensis* | 26 | 1 | TZ | 2011,2012 |
| *Mus bufo* | 17 | | KE | 2010 |
| *Mus* cf. *gerbillus* | 1 | | KE | 2010 |
| *Mus* cf. *gratus* | 18 | | KE/CD/TZ | 2010/2012/2013 |
| *Mus mahomet* | 33 | | ET | 2010 |
| *Mus minutoides* | 35 | | TZ/ZM/KE/MZ | 2007,2013/2009,2010/2010/2011 |
| *Mus musculus* | 2 | | KE | 2010 |
| *Mus* sp. | 103 | | TZ/CD/MG | 2008-2010/2007,2010,2012,2013/2010 |
| *Mus triton* | 23 | | ZM/KE/TZ | 2009/2011/2013 |
| *Myomyscus brockmani* | 36 | | KE | 2010,2011 |
| *Oenomys hypoxanthus* | 11 | | CD/KE/TZ | 2010/2010/2013 |
| *Oenomys* sp. | 3 | | CD | 2010 |
| *Otomys* sp. | 6 | | CD/TZ/KE | 2007/2007,2009,2010/2010 |
| *Otomys tropicalis* | 4 | | KE | 2010 |
| *Praomys delectorum* | 103 | | KE/TZ | 2010/2007,2013 |
| *Praomys jacksoni* | 140 | 7 | ZM/KE/CD/TZ | 2009/2010,2011/2010,2012/2013 |
| *Praomys lukoleae* | 6 | | CD | 2013 |
| *Praomys minor* | 2 | | ZM | 2009 |
| *Praomys* sp. | 390 | 2 | TZ/KE/CD | 2006,2008-2011/2010/2010,2012,2013 |
| *Rattus rattus* | 250 | 1 | CD/TZ/ZM/ET/KE/MG/MZ | 2007,2012,2013/2006,2007,2009/2009,2010/2010/2010/2010/2011 |
| *Rattus* sp. | 84 | | TZ/MZ | 2007,2009,2010,2013/2011 |
| *Rhabdomys dilectus* | 27 | | KE/TZ | 2010/2008,2009,2013 |
| *Stenocephalemys albipes* | 178 | 2 | ET | 2010/2012 |
| *Stenocephalemys albocaudatus* | 10 | | ET | 2012 |
| *Stenocephalemys griseicauda* | 10 | | ET | 2012 |
| *Stenocephalemys* sp. | 9 | | ET | 2012 |
| *Stochomys longicaudatus* | 5 | | CD | 2010,2012,2013 |
| *Uranomys ruddi* | 9 | | MZ/TZ | 2011 |
| *Zelotomys hildegardeae* | 2 | | TZ | 2013 |

| | | | | | |
|---|---|---|---|---|---|
| Nesomyidae | *Beamys hindei* | 4 | | MZ | 2011 |
| | *Beamys major* | 6 | | TZ | 2007,2011 |
| | *Cricetomys gambianus* | 6 | | TZ | 2007 |
| | *Cricetomys* sp. | 5 | | CD/MZ | 2010/2011 |
| | *Dendromus insignis* | 8 | | KE | 2010,2011 |
| | *Dendromus nyikae* | 8 | | TZ | 2007 |
| | *Eliurus myoxinus* | 5 | | MG | 2010 |
| | *Saccostomus campestris* | 13 | 1 | ZM/MZ | 2009,2010/2011 |
| | *Saccostomus mearnsi* | 4 | | KE | 2010, 2011 |
| | *Saccostomus umbriventer* | 2 | | KE | 2010 |
| Sciuridae | *Funisciurus anerythus* | 3 | | CD | 2010 |
| | *Funisciurus pyrrhopus* | 3 | | CD | 2013 |
| | *Funisciurus* sp. | 4 | | CD | 2010 |
| | *Heliosciurus rufobrachium* | 1 | | TZ | 2013 |
| | *Paraxerus ochraceus* | 6 | | TZ/KE | 2007/2010 |
| | *Xerus* sp. | 3 | | TZ | 2013 |
| Spalacidae | *Tachyoryctes splendens* | 13 | 1 | KE/TZ | 2010,2011/2008,2013 |
| Thryonomyidae | *Thryonomys* sp. | 1 | | | |
| **Subtotal** | **116 species** | **3788 samples** | **78 positives** | | |
| **Total** | **161 species** | **4303** | **80** | | |

* The current nomenclature does not reliably delineate the genetic lineages in the *Graphiurus* genus. Based on the experts' opinion that co-authored this manuscript, *Graphiurus* individuals from those countries are genetically different and most likely represent multiple species. Although we cannot update the rodent species taxonomy within the scope of this manuscript, we will use the *Graphiurus* sp. name and count them all together as a single species.

**Supplementary table S2:** Primers used in this study.

| Name | Sequence (5'-3') | Amplicon size (bp) | Study |
|---|---|---|---|
| AK4340F1 | GTACTTGCTACTGCNACNCC | 299 | Kapoor et al., 2013 |
| AK4360R1 | TACCCTGTCATAAGGGCRTC | | Kapoor et al., 2013 |
| AK4340F2 | CTTGCTACTGCNACNCCWCC | 296 | Kapoor et al., 2013 |
| AK4360R2 | TACCCTGTCATAAGGGCRTCNGT | | Kapoor et al., 2013 |
| MOZ094-5110F | ACTGGCTTAGGCTTAGGA | 287 | This study |
| MOZ094-5438R | AAGCAAGCCAAGCCAGCG | | This study |
| MOZ133-582F | CTACAAGCCTATCCCTCTCA | 223 | This study |
| MOZ133-978R | ACTAGCGAGCCAACCAAA | | This study |
| MOZ133-1741F | ACATTGTATCGCGGTTCG | 172 | This study |
| MOZ133-1918R | GTATTGCCATAATTACAATGTGG | | This study |
| MOZ133-2085F | CTTAGGTTGGCTGCTCTGTA | 211 | This study |
| MOZ133-2301R | AGTGGTGACCTCCAGTTG | | This study |
| MOZ133-2492F | TTGCTCGTCTGGCGGAGAA | 133 | This study |
| MOZ133-2837R | ACATCTAGGTCTTCATAGCC | | This study |
| MOZ133-3159F | CTCATCAGGCGCTCCGTT | 195 | This study |
| MOZ133-3365R | GCAACAAGGATTTCGATGG | | This study |
| MOZ133-3393F | ATTACCCATGGAATACTATAAACA | 161 | This study |
| MOZ133-3556R | AATACGTAAGCCGCGAGC | | This study |
| MOZ133-4078F | TTCTCAATTTCATTGACCACA | 162 | This study |
| MOZ133-4410R | AACGCGTTGTTTCATGGTA | | This study |
| MOZ133-4452F | TCTAGAGGCTAAAGCAGG | 137 | This study |
| MOZ133-4690R | CAAAGAGATCAATGGCCAC | | This study |
| MOZ133-5248F | TCTATGTCAGAGTGGTCTCA | 174 | This study |
| MOZ133-5460R | AAGTAGCCTGATGATGGTTG | | This study |
| MOZ133-5699F | TTGTACGTTCCACTTGTGG | 129 | This study |
| MOZ133-5920R | CTCCTCTGAGCTCATACTTT | | This study |
| MOZ133-7434F | CCGTAACTGTAGAGCATCG | 93 | This study |
| MOZ133-7889R | AAGTGTACTGCTATCACCC | | This study |
| MOZ133-7978F | CCTTACATAATCCAGGCCC | 170 | This study |
| MOZ133-8183R | CAGCTCAATAAGGTCTTTCC | | This study |
| MOZ329-7156P1F1 | GTGATGAATGAGATCGAAAGTG | 385 | This study |
| MOZ329-7737P1R1 | GGATCTCAGCGCTGCTCT | | This study |
| MOZ329-7156P2F1 | GGGTTGTATGAAAATGGTTCCT | 364 | This study |
| MOZ329-7737P2R1 | GGACTGTCTCCTCCTTCTCG | | This study |
| MOZ329-7540P3F1 | GGGTCCTGTTGGGGAAAG | 119 | This study |
| MOZ329-8129P3R1† | CGCTCCCGTCTGTCAAGC | | This study |
| MOZ329-7205P1F2 | CAGCAAGCCGGGCGTCCAG | 223 | This study |
| MOZ329-7513P1R2 | TCTTCGAGGAGCCTCTTGAT | | This study |
| MOZ329-7178P2F2 | TCGATCCACCAACCATGC | 184 | This study |
| MOZ329-7490P2R2 | GCAGACACTGGTTCCGGTT | | This study |
| MOZ329-7553P3F2 | GAAAGAAACCCGAGGATCCG | 106 | This study |
| MOZ329-8129P3R2† | CGCTCCCGTCTGTCAAGC | | This study |
| TA100-7156P1F1 | AAACAGGATCAGGGGGTTG | 571 | This study |
| TA100-7737P1R1 | AGATTGCGTTCTCCACCTC | | This study |
| TA100-7156P2F1 | GGTTTGTATGAGAATGGGTCC | 367 | This study |
| TA100-7737P2R1 | GGATTGTCTCCTCTTTCTCG | | This study |
| TA100-7414P3F1 | AGACTGGACCCTGGCAAG | 185 | This study |
| TA100-8108P3R1 | CTGATCGAGCATGTCCATCA | | This study |
| TA100-7205P1F2 | AACAGCTGAAGCCAGGGG | 300 | This study |
| TA100-7513P1R2 | CTGCGAGCATGTCCTTGAAT | | This study |
| TA100-7178P2F2 | TCGATCCACCATCTATGCCT | 184 | This study |
| TA100-7490P2R2 | GCAGCTACTGGTTCTGGC | | This study |
| TA100-7428P3F2 | CAAGTCACAATGCAGACAGG | 148 | This study |
| TA100-7852P3R2 | CAAGCTGGTACTTTATGTTGTAAT | | This study |
| TA338-1157F | TCAATTGGTCTGTCAGGAAC | 388 | This study |
| TA338-1544R | AGAGTCGCTTTCGCCAAC | | This study |
| TA338-2204F | ACCACTTCAGGGTCGGTG | 349 | This study |
| TA338-2552R | TAAGCCATTACGCCCTTG | | This study |
| TA338-3303F | TCTAGCTCTTACTGGCAGG | 318 | This study |
| TA338-3620R | TGCATAACACCGTCACGC | | This study |

† denotes an identical primer sequence

**Supplementary table S3:** Specimen information of all available and novel hepacivirus genomes used in the phylogenetic analysis of study.

| Accession number | Isolate name | Host family | Host species | Host type | Country | Sampling year |
|---|---|---|---|---|---|---|
| AB863589 | JPN3 | Equidae | *Equus ferus caballus* | Horse | Japan | 2013 |
| AF179612 | GBV-B | Cebidae | *Saguinus mystax* | Primate | NA | 1995 |
| NC_004102 | HCV1 | Hominidae | *Homo sapiens* | Human | NA | NA |
| NC_009823 | HCV2 | Hominidae | *Homo sapiens* | Human | NA | NA |
| NC_009824 | HCV3 | Hominidae | *Homo sapiens* | Human | New Zealand | NA |
| NC_009825 | HCV4 | Hominidae | *Homo sapiens* | Human | Egypt | NA |
| NC_009826 | HCV5 | Hominidae | *Homo sapiens* | Human | United Kingdom | NA |
| NC_009827 | HCV6 | Hominidae | *Homo sapiens* | Human | NA | NA |
| EF108306 | HCV7 | Hominidae | *Homo sapiens* | Human | Canada | 2003 |
| JF744991 | NA | Canidae | *Canis lupus familiaris* | Dog | USA | 2011 |
| JQ434001 | NZP-1 | Equidae | *Equus ferus caballus* | Horse | USA | 2011 |
| JQ434002 | G1-073 | Equidae | *Equus ferus caballus* | Horse | USA | 2011 |
| JQ434003 | A6-006 | Equidae | *Equus ferus caballus* | Horse | USA | 2011 |
| JQ434004 | B10-022 | Equidae | *Equus ferus caballus* | Horse | USA | 2011 |
| JQ434005 | F8-068 | Equidae | *Equus ferus caballus* | Horse | USA | 2011 |
| JQ434006 | G5-077 | Equidae | *Equus ferus caballus* | Horse | USA | 2011 |
| JQ434007 | H10-094 | Equidae | *Equus ferus caballus* | Horse | USA | 2011 |
| JQ434008 | H3-011 | Equidae | *Equus ferus caballus* | Horse | UK | 2011 |
| JX948116 | EF369_11J | Equidae | *Equus ferus caballus* | Horse | UK | 2011 |
| KC411777 | RMU10-3382 | Cricetidae | *Myodes glareolus* | Rodent | Germany | 2010 |
| KC411784 | NLR07-oct70 | Cricetidae | *Myodes glareolus* | Rodent | Netherlands | 2007 |
| KC411796 | NLR08-365 | Cricetidae | *Myodes glareolus* | Rodent | Netherlands | 2008 |
| KC411806 | SAR-3 | Muridae | *Rhabdomys pumilio* | Rodent | South Africa | 2008 |
| KC411807 | SAR-46 | Muridae | *Rhabdomys pumilio* | Rodent | South Africa | 2008 |
| KC551800 | GHV-1_BWC08 | Cercopithecidae | *Colobus guereza* | Primate | Uganda | 2010 |
| KC551801 | GHV-1_BWC05 | Cercopithecidae | *Colobus guereza* | Primate | Uganda | 2010 |
| KC551802 | GHV-2_BWC04 | Cercopithecidae | *Colobus guereza* | Primate | Uganda | 2010 |
| KC796074 | PDB-829 | Hipposideridae | *Hipposideros vittatus* | Bat | Kenya | 2011 |
| KC796077 | PDB-112 | Hipposideridae | *Hipposideros vittatus* | Bat | Kenya | 2010 |
| KC796078 | PDB-491.1 | Molossidae | *Otomops martiensseni* | Bat | Kenya | 2011 |
| KC796090 | PDB-452 | Molossidae | *Otomops martiensseni* | Bat | Kenya | 2010 |
| KC796091 | PDB-445 | Molossidae | *Otomops martiensseni* | Bat | Kenya | 2010 |
| KC815310 | RHV-339 | Cricetidae | *Peromyscus maniculatus* | Rodent | USA | 2008 |
| KC815312 | RHV-089 | Cricetidae | *Peromyscus maniculatus* | Rodent | USA | 2008 |
| KF177391 | DH1 | Equidae | *Equus ferus caballus* | Horse | Hungary | 2013 |
| KJ472766 | WSU-2013 | Equidae | *Equus ferus caballus* | Horse | USA | 2013 |
| KJ950938 | NrHV-1_NYC-C12 | Muridae | *Rattus norvegicus* | Rodent | USA | 2013 |
| KJ950939 | NrHV-2_NYC-E43 | Muridae | *Rattus norvegicus* | Rodent | USA | 2012 |
| KP265943 | GHC25 | Bovidae | *Bos taurus* | Cattle | Ghana | 2011 |
| KP265946 | GHC52 | Bovidae | *Bos taurus* | Cattle | Ghana | 2011 |
| KP265947 | GHC55 | Bovidae | *Bos taurus* | Cattle | Ghana | 2011 |
| KP265948 | GHC85 | Bovidae | *Bos taurus* | Cattle | Ghana | 2011 |
| KP265950 | GHC100 | Bovidae | *Bos taurus* | Cattle | Ghana | 2011 |
| KP325401 | NZP-1 | Equidae | *Equus ferus caballus* | Horse | USA | 2011 |
| KP641123 | B1 | Bovidae | *Bos taurus* | Cattle | Germany | 2013 |
| KP641124 | 209 | Bovidae | *Bos taurus* | Cattle | Germany | 2014 |
| KP641125 | 379 | Bovidae | *Bos taurus* | Cattle | Germany | 2014 |
| KP641126 | 438 | Bovidae | *Bos taurus* | Cattle | Germany | 2014 |
| KP641127 | 463 | Bovidae | *Bos taurus* | Cattle | Germany | 2014 |
| MK737639 | HCL-1 | Anatidae | *Anas platyrhynchos domesticus* | Bird | China | 2018 |
| MK737640 | HCL-2 | Anatidae | *Anas platyrhynchos domesticus* | Bird | China | 2018 |
| MK737641 | HCL-3 | Anatidae | *Anas platyrhynchos domesticus* | Bird | China | 2018 |
| MF775364 | SZCDC70 | Soricidae | *Suncus murinus* | Shrew | China | 2015 |
| MH824541 | H2-L41 | Indriidae | *Propithecus diadema* | Primate | Madagascar | 2011 |
| MH824540 | H2-L40 | Indriidae | *Propithecus diadema* | Primate | Madagascar | 2011 |
| MH824539 | H2-L25 | Indriidae | *Propithecus diadema* | Primate | Madagascar | 2011 |
| MH824542 | H6-L83 | Indriidae | *Propithecus diadema* | Primate | Madagascar | 2013 |
| MH824543 | H5-L75 | Indriidae | *Propithecus diadema* | Primate | Madagascar | 2012 |
| MH027948 | BH181 | Bovidae | *Bos taurus* | Cattle | Germany | NA |
| MH027953 | BH204 | Bovidae | *Bos taurus* | Cattle | Germany | NA |
| MG781019 | BR_RN034B019 | Bovidae | *Bos taurus* | Cattle | Brazil | 2013 |
| MG781018 | BR_MA236B017 | Bovidae | *Bos taurus* | Cattle | Brazil | 2013 |
| MG257793 | BovHepV/GD/01 | Bovidae | *Bos taurus* | Cattle | China | 2017 |
| MG257794 | BovHepV/GD/02 | Bovidae | *Bos taurus* | Cattle | China | 2017 |
| MH027992 | H56 | Equidae | *Equus ferus caballus* | Horse | Germany | NA |
| MH028007 | H628 | Equidae | *Equus ferus caballus* | Horse | Germany | NA |
| MH028000 | H268 | Equidae | *Equus ferus caballus* | Horse | Germany | NA |
| MH027993 | H57 | Equidae | *Equus ferus caballus* | Horse | Germany | NA |
| MH028004 | H581 | Equidae | *Equus ferus caballus* | Horse | Germany | NA |

| | | | | | | |
|---|---|---|---|---|---|---|
| MH028005 | H593 | Equidae | *Equus ferus caballus* | Horse | Germany | NA |
| MH027998 | H170 | Equidae | *Equus ferus caballus* | Horse | Germany | NA |
| KX056116 | K-061 | Equidae | *Equus ferus caballus* | Horse | South Korea | 2015 |
| KX056117 | K-062 | Equidae | *Equus ferus caballus* | Horse | South Korea | 2015 |
| MH027995 | H105 | Equidae | *Equus ferus caballus* | Horse | Germany | NA |
| MH027999 | H179 | Equidae | *Equus ferus caballus* | Horse | Germany | NA |
| MH028001 | H285 | Equidae | *Equus ferus caballus* | Horse | Germany | NA |
| MH028002 | H286 | Equidae | *Equus ferus caballus* | Horse | Germany | NA |
| MH027996 | H143 | Equidae | *Equus ferus caballus* | Horse | Germany | NA |
| KX421286 | B82 | Equidae | *Equus asinus asinus* | Donkey | Bulgaria | 2015 |
| KX421287 | B89 | Equidae | *Equus asinus asinus* | Donkey | Bulgaria | 2015 |
| KT880191 | R09-249 | Equidae | *Equus asinus asinus* | Donkey | France | 1979 |
| KT880192 | R09-250 | Equidae | *Equus asinus asinus* | Donkey | France | 1979 |
| KT880193 | R09-251 | Equidae | *Equus asinus asinus* | Donkey | France | 1979 |
| MF152651 | Guangzhou/6 | Equidae | *Equus ferus caballus* | Horse | China | 2016 |
| MF152652 | Guangzhou/33 | Equidae | *Equus ferus caballus* | Horse | China | 2016 |
| KY370095 | IM2014 | Dipodidae | *Dipus sagitta* | Rodent | China | 2014 |
| KY370094 | Tibet2014 | Cricetidae | *Neodon clarkei* | Rodent | China | 2014 |
| KY370092 | IM2014 | Muridae | *Meriones meridianus* | Rodent | China | 2014 |
| KX905133 | rn-1 | Muridae | *Rattus norvegicus* | Rodent | USA | 2015 |
| MF113386 | SD-1 | Muridae | *Rattus norvegicus* | Rodent | USA | 2014 |
| MH370348 | On/2012 | Cricetidae | *Oligoryzomys nigripes* | Rodent | Brazil | 2012 |
| MG211815 | GS2015 | Sciuridae | *Spermophilus dauricus* | Rodent | China | 2015 |
| MG600414 | 05VZ-14-118 | Spalacidae | *Rhizomys pruinosus* | Rodent | Vietnam | 2015 |
| MG600412 | 05VZ-14-104 | Spalacidae | *Rhizomys pruinosus* | Rodent | Vietnam | 2015 |
| MG600413 | 05VZ-14-103 | Spalacidae | *Rhizomys pruinosus* | Rodent | Vietnam | 2015 |
| MG600415 | 05VZ-14-119 | Spalacidae | *Rhizomys pruinosus* | Rodent | Vietnam | 2015 |
| MG822666 | B349/PAN/2014 | Echimyidae | *Proechimys semispinosus* | Rodent | Panama | 2014 |
| KY370091 | IM2014 | Dipodidae | *Allactaga sibirica* | Rodent | China | 2014 |
| MG599988 | PXJHG3419 | Eublepharidae | *Goniurosaurus luii* | Lizard | China | NA |
| MG599987 | YLSHG5584 | Sphaerodactylidae | *Teratoscincus roborowskii* | Lizard | China | NA |
| MG599992 | DHHHBHGS10983 | Urolophidae | *Urolophus aurantiacus* | Cartilaginous fish | China | NA |
| MG599997 | NHYJG60710 | Chimaeridae | *Chimaera sp* | Cartilaginous fish | China | NA |
| MG599989 | LPXYG13170 | Gekkonidae | *Hemidactylus bowringii* | Lizard | China | NA |
| MG599993 | FZFYG124617 | Protopteridae | *Protopterus annectens* | Lungfish | Nigeria | NA |
| MG599991 | BWLTYG5315 | Rhinobatidae | *Rhinobatos hynnicephalus* | Cartilaginous fish | China | NA |
| MG599994 | NHJSG30635 | Triakidae | *Mustelus manazo* | Cartilaginous fish | China | NA |
| MG599998 | RBCSG7845 | Triakidae | *Mustelus manazo* | Cartilaginous fish | China | NA |
| MG334001 | Hepacivirus sp. | Emydidae | *Trachemys scripta elegans* | Turtle | USA | 2008 |
| MG599999 | WHJYGF75270 | Trionychidae | *Pelodiscus sinensis* | Turtle | China | NA |
| MG599995 | NHJSG30261 | Squalidae | *Squalus brevirostris* | Cartilaginous fish | China | NA |
| MG599996 | NHYJG60722 | Chimaeridae | *Chimaera sp* | Cartilaginous fish | China | NA |
| MG600000 | WHWGGF64311 | Geoemydinae | *Mauremys megalocephala* | Turtle | China | NA |
| MG599990 | XMLMGHepa10640 | Muraenidae | *Gymnothorax reticularis* | Ray-finned fish | China | NA |
| MN635447 | VERT31 | Phalangeridae | *Trichosurus vulpecula* | Possum | Australia | NA |
| MN062427 | NA03-001 | Accipitridae | *Haliaeetus leucocephalus* | Bird | USA | 2002 |
| **Subtotal** | **115 available genomes** | | | | | |
| MN587650 | CRT125-A/COD/2010 | Muridae | *Lophuromys dudui* | Rodent | DRC | 2010 |
| MN587651 | CRT125-B/COD/2010 | Muridae | *Lophuromys dudui* | Rodent | DRC | 2010 |
| MN587652 | CRT125-C/COD/2010 | Muridae | *Lophuromys dudui* | Rodent | DRC | 2010 |
| MN587653 | CRT125-D/COD/2010 | Muridae | *Lophuromys dudui* | Rodent | DRC | 2010 |
| MN587654 | CRT352-A/COD/2010 | Muridae | *Lophuromys dudui* | Rodent | DRC | 2010 |
| MN587655 | CRT352-B/COD/2010 | Muridae | *Lophuromys dudui* | Rodent | DRC | 2010 |
| MN587656 | CRT382/COD/2010 | Muridae | *Lophuromys dudui* | Rodent | DRC | 2010 |
| MN587657 | CRT471/COD/2010 | Muridae | *Lophuromys dudui* | Rodent | DRC | 2010 |
| MN587658 | CRT490-A/COD/2010 | Muridae | *Lophuromys dudui* | Rodent | DRC | 2010 |
| MN587659 | CRT490-B/COD/2010 | Muridae | *Lophuromys dudui* | Rodent | DRC | 2010 |
| MN587660 | CRT490-C/COD/2010 | Muridae | *Lophuromys dudui* | Rodent | DRC | 2010 |
| MN587661 | CRT64-A/COD/2010 | Muridae | *Lophuromys dudui* | Rodent | DRC | 2010 |
| MN587662 | CRT64-B/COD/2010 | Muridae | *Lophuromys dudui* | Rodent | DRC | 2010 |
| MN587663 | CRT64-C/COD/2010 | Muridae | *Lophuromys dudui* | Rodent | DRC | 2010 |
| MN587664 | CRT64-D/COD/2010 | Muridae | *Lophuromys dudui* | Rodent | DRC | 2010 |
| MN587665 | CRT64-E/COD/2010 | Muridae | *Lophuromys dudui* | Rodent | DRC | 2010 |
| MN564789 | CRT682/COD/2010 | Gliridae | *Graphiurus kelleni* | Rodent | DRC | 2010 |
| MN587666 | CRT74/COD/2010 | Muridae | *Lophuromys dudui* | Rodent | DRC | 2010 |
| MN564790 | ETH674/ETH/2012 | Muridae | *Stenocephalemys albipes* | Rodent | Ethiopia | 2012 |
| MN587667 | MOZ094/MOZ/2011 | Muridae | *Lophuromys machangui* | Rodent | Mozambique | 2011 |
| MN587668 | MOZ133/MOZ/2011 | Muridae | *Lophuromys machangui* | Rodent | Mozambique | 2011 |
| MN587669 | MOZ329-A/MOZ/2011 | Muridae | *Lophuromys machangui* | Rodent | Mozambique | 2011 |
| MN587670 | MOZ329-B/MOZ/2011 | Muridae | *Lophuromys machangui* | Rodent | Mozambique | 2011 |
| MN587671 | MOZ329-C/MOZ/2011 | Muridae | *Lophuromys machangui* | Rodent | Mozambique | 2011 |
| MN587695 | TA085/TZA/2013 | Muridae | *Lophuromys stanleyi* | Rodent | Tanzania | 2013 |
| MN587696 | TA100-A/TZA/2013 | Muridae | *Lophuromys stanleyi* | Rodent | Tanzania | 2013 |
| MN587697 | TA100-B/TZA/2013 | Muridae | *Lophuromys stanleyi* | Rodent | Tanzania | 2013 |
| MN587698 | TA100-C/TZA/2013 | Muridae | *Lophuromys stanleyi* | Rodent | Tanzania | 2013 |
| MN564792 | TA132/TZA/2013 | Muridae | *Praomys jacksoni* | Rodent | Tanzania | 2013 |
| MN564793 | TA142/TZA/2013 | Muridae | *Praomys jacksoni* | Rodent | Tanzania | 2013 |

| | | | | | | |
|---|---|---|---|---|---|---|
| MN564794 | TA152/TZA/2013 | Muridae | *Praomys jacksoni* | Rodent | Tanzania | 2013 |
| MN587672 | TA275/TZA/2013 | Muridae | *Lophuromys laticeps* | Rodent | Tanzania | 2013 |
| MN587673 | TA293-A/TZA/2013 | Muridae | *Lophuromys laticeps* | Rodent | Tanzania | 2013 |
| MN587674 | TA293-B/TZA/2013 | Muridae | *Lophuromys laticeps* | Rodent | Tanzania | 2013 |
| MN587675 | TA293-C/TZA/2013 | Muridae | *Lophuromys laticeps* | Rodent | Tanzania | 2013 |
| MN587676 | TA293-D/TZA/2013 | Muridae | *Lophuromys laticeps* | Rodent | Tanzania | 2013 |
| MN587677 | TA293-E/TZA/2013 | Muridae | *Lophuromys laticeps* | Rodent | Tanzania | 2013 |
| MN564791 | TA338/TZA/2013 | Muridae | *Mastomys natalensis* | Rodent | Tanzania | 2013 |
| MN587678 | TA498-A/TZA/2013 | Muridae | *Lophuromys machangui* | Rodent | Tanzania | 2013 |
| MN587679 | TA498-B/TZA/2013 | Muridae | *Lophuromys machangui* | Rodent | Tanzania | 2013 |
| MN587680 | TA498-C/TZA/2013 | Muridae | *Lophuromys machangui* | Rodent | Tanzania | 2013 |
| MN587681 | TA528/TZA/2013 | Muridae | *Lophuromys machangui* | Rodent | Tanzania | 2013 |
| MN587682 | TA529-A/TZA/2013 | Muridae | *Lophuromys machangui* | Rodent | Tanzania | 2013 |
| MN587683 | TA529-B/TZA/2013 | Muridae | *Lophuromys machangui* | Rodent | Tanzania | 2013 |
| MN587684 | TA529-C/TZA/2013 | Muridae | *Lophuromys machangui* | Rodent | Tanzania | 2013 |
| MN587685 | TA529-D/TZA/2013 | Muridae | *Lophuromys machangui* | Rodent | Tanzania | 2013 |
| MN587686 | TA531-A/TZA/2013 | Muridae | *Lophuromys machangui* | Rodent | Tanzania | 2013 |
| MN587687 | TA531-B/TZA/2013 | Muridae | *Lophuromys machangui* | Rodent | Tanzania | 2013 |
| MN587688 | TA531-C/TZA/2013 | Muridae | *Lophuromys machangui* | Rodent | Tanzania | 2013 |
| MN587689 | TA531-D/TZA/2013 | Muridae | *Lophuromys machangui* | Rodent | Tanzania | 2013 |
| MN587690 | TA531-E/TZA/2013 | Muridae | *Lophuromys machangui* | Rodent | Tanzania | 2013 |
| MN587691 | TA532-A/TZA/2013 | Muridae | *Lophuromys machangui* | Rodent | Tanzania | 2013 |
| MN587692 | TA532-B/TZA/2013 | Muridae | *Lophuromys machangui* | Rodent | Tanzania | 2013 |
| MN587693 | TA532-C/TZA/2013 | Muridae | *Lophuromys machangui* | Rodent | Tanzania | 2013 |
| MN587694 | TA532-D/TZA/2013 | Muridae | *Lophuromys machangui* | Rodent | Tanzania | 2013 |
| MN555567 | TZ25757/TZA/2011 | Muridae | *Acomys wilsoni* | Rodent | Tanzania | 2011 |
| **Subtotal** | **56 novel genomes** | | | | | |
| **Total** | **171 genomes** | | | | | |

**Supplementary table S4:** Cytochrome b haplotype information of all available and novel rodent hepacivirus hosts species.

| Accession number | Host species | Country | Sampling year |
|---|---|---|---|
| MN616976 | Lophuromys dudui | Democratic Republic of the Congo | 2010 |
| MN616984 | Graphiurus kelleni | Democratic Republic of the Congo | 2010 |
| MN616986 | Stenocephalemys albipes | Ethiopia | 2010 |
| MN616989 | Lophuromys machangui | Mozambique | 2011 |
| MN616994 | Lophuromys stanleyi | Tanzania | 2013 |
| MN616998 | Praomys jacksoni | Tanzania | 2013 |
| KY595882 | Lophuromys laticeps | Tanzania | 2013 |
| MN617006 | Mastomys natalensis | Tanzania | 2013 |
| MN617014 | Acomys wilsoni | Tanzania | 2011 |
| KY753939 | Allactaga sibirica | China | 2014 |
| KX399741 | Dipus sagitta | China | 2014 |
| JQ065613 | Meriones meridianus | China | 2014 |
| KJ612494 | Myodes glareolus | Netherlands | 2007 |
| KP190220 | Neodon clarkei | China | 2014 |
| GU126530 | Oligoryzomys nigripes | Brazil | 2012 |
| DQ385827 | Peromyscus maniculatus | USA | 2008 |
| NC_039103 | Proechimys semispinosus | Panama | 2014 |
| NC_001665 | Rattus norvegicus | USA | 2013 |
| AF533116 | Rhabdomys pumilio | South Africa | 2008 |
| NC_021478 | Rhizomys pruinosus | Vietnam | 2015 |
| NC_027283 | Spermophilus dauricus | China | 2015 |

**Supplementary table S5:** Specimen characteristics of all hepacivirus positive cases detected.

| Sample | Species | Family | Locality | Country | Year | Latitude | Longitude |
|---|---|---|---|---|---|---|---|
| 2240 | *Mastomys awashensis* | Muridae | Aroresha | Ethiopia | 2010 | 12.4166667 | 39.55 |
| ETH019 | *Stenocephalemys albipes* | Muridae | Menangasha | Ethiopia | 2012 | 8.9661 | 38.5495 |
| ETH505 | *Mastomys awashensis* | Muridae | Lake Hashenge | Ethiopia | 2012 | 12.6394 | 39.5383 |
| ETH674 | *Stenocephalemys albipes* | Muridae | Mizan Tefari | Ethiopia | 2012 | 7.06036 | 35.67192 |
| KE107 | *Gerbilliscus vicinus* | Muridae | Wamba | Kenya | 2010 | 0.9795 | 37.3272 |
| KE118 | *Graphiurus* sp. 1* | Gliridae | Marsabit | Kenya | 2010 | 2.3092 | 37.9659 |
| KE142 | *Acomys kempi* | Muridae | South Horr | Kenya | 2010 | 2.1038 | 36.8931 |
| KE153 | *Acomys kempi* | Muridae | South Horr | Kenya | 2010 | 2.0973 | 36.8992 |
| KE389 | *Hylomyscus stella* | Muridae | Kakamega | Kenya | 2010 | 0.2382 | 34.8647 |
| KE645 | *Gerbilliscus vicinus* | Muridae | Kapiti plains | Kenya | 2011 | -1.4842 | 37.0552 |
| KE839 | *Lophuromys zena* | Muridae | Nyahururu forest | Kenya | 2011 | 0.0449 | 36.3734 |
| KE840 | *Lophuromys zena* | Muridae | Nyahururu forest | Kenya | 2011 | 0.0449 | 36.3734 |
| TA085 | *Lophuromys stanleyi* | Muridae | Minziro FR | Tanzania | 2013 | -1.031 | 31.572 |
| TA094 | *Praomys jacksoni* | Muridae | Minziro FR | Tanzania | 2013 | -1.031 | 31.572 |
| TA100 | *Lophuromys stanleyi* | Muridae | Minziro FR | Tanzania | 2013 | -1.031 | 31.572 |
| TA106 | *Lophuromys stanleyi* | Muridae | Minziro FR | Tanzania | 2013 | -1.031 | 31.572 |
| TA109 | *Lophuromys stanleyi* | Muridae | Minziro FR | Tanzania | 2013 | -1.031 | 31.572 |
| TA132 | *Praomys jacksoni* | Muridae | Minziro FR | Tanzania | 2013 | -1.031 | 31.572 |
| TA142 | *Praomys jacksoni* | Muridae | Minziro FR | Tanzania | 2013 | -1.031 | 31.572 |
| TA151 | *Praomys jacksoni* | Muridae | Minziro FR | Tanzania | 2013 | -1.031 | 31.572 |
| TA152 | *Praomys jacksoni* | Muridae | Minziro FR | Tanzania | 2013 | -1.031 | 31.572 |
| TA156 | *Praomys jacksoni* | Muridae | Minziro FR | Tanzania | 2013 | -1.031 | 31.572 |
| TA166 | *Glauconycteris atra* | Vespertilionidae | Minziro FR | Tanzania | 2013 | -1.031 | 31.572 |
| TA168 | *Glauconycteris atra* | Vespertilionidae | Minziro FR | Tanzania | 2013 | -1.031 | 31.572 |
| TA275 | *Lophuromys laticeps* | Muridae | Bitale (North of Kigoma) | Tanzania | 2013 | -4.731659 | 29.706526 |
| TA289 | *Lophuromys laticeps* | Muridae | Bitale (North of Kigoma) | Tanzania | 2013 | -4.731659 | 29.706526 |
| TA293 | *Lophuromys laticeps* | Muridae | Bitale (North of Kigoma) | Tanzania | 2013 | -4.731659 | 29.706526 |
| TA338 | *Mastomys natalensis* | Muridae | Makongoro | Tanzania | 2013 | -6.451433 | 31.115017 |
| TA497 | *Lophuromys machangui* | Muridae | Mt Ngozi (Poroto Range) | Tanzania | 2013 | -9.04075 | 33.5732 |
| TA498 | *Lophuromys machangui* | Muridae | Mt Ngozi (Poroto Range) | Tanzania | 2013 | -9.04075 | 33.5732 |
| TA502 | *Lophuromys machangui* | Muridae | Mt Ngozi (Poroto Range) | Tanzania | 2013 | -9.04075 | 33.5732 |
| TA503 | *Lophuromys machangui* | Muridae | Mt Ngozi (Poroto Range) | Tanzania | 2013 | -9.04075 | 33.5732 |
| TA504 | *Lophuromys machangui* | Muridae | Mt Ngozi (Poroto Range) | Tanzania | 2013 | -9.04075 | 33.5732 |
| TA527 | *Lophuromys machangui* | Muridae | Mt Ngozi (Poroto Range) | Tanzania | 2013 | -9.04075 | 33.5732 |
| TA528 | *Lophuromys machangui* | Muridae | Mt Ngozi (Poroto Range) | Tanzania | 2013 | -9.04075 | 33.5732 |
| TA529 | *Lophuromys machangui* | Muridae | Mt Ngozi (Poroto Range) | Tanzania | 2013 | -9.04075 | 33.5732 |
| TA530 | *Lophuromys machangui* | Muridae | Mt Ngozi (Poroto Range) | Tanzania | 2013 | -9.04075 | 33.5732 |
| TA531 | *Lophuromys machangui* | Muridae | Mt Ngozi (Poroto Range) | Tanzania | 2013 | -9.04075 | 33.5732 |
| TA532 | *Lophuromys machangui* | Muridae | Mt Ngozi (Poroto Range) | Tanzania | 2013 | -9.04075 | 33.5732 |
| TA533 | *Lophuromys machangui* | Muridae | Mt Ngozi (Poroto Range) | Tanzania | 2013 | -9.04075 | 33.5732 |
| VM067 | *Saccostomus campestris* | Nesomyidae | Chipata - Mamarula Camp | Zambia | 2010 | -13.5823 | 32.6099 |
| CRT112 | *Lophuromys dudui* | Muridae | Yaikela Ligne Pitfall Kaswera | DRC | 2010 | 0.81361 | 24.27913 |
| CRT125 | *Lophuromys dudui* | Muridae | Yaikela Ligne 2 | DRC | 2010 | 0.82099 | 24.27678 |
| CRT352 | *Lophuromys dudui* | Muridae | Bomane Ligne 1 KGB | DRC | 2010 | 1.2 | 23.7 |
| CRT353 | *Hybomys* sp. | Muridae | Bomane Ligne 2 KGB | DRC | 2010 | 1.2 | 23.7 |
| CRT382 | *Lophuromys dudui* | Muridae | Bomane Ligne 2 KGB | DRC | 2010 | 1.2 | 23.7 |
| CRT471 | *Lophuromys dudui* | Muridae | Bomane Champ Nic2 | DRC | 2010 | 1.27374 | 23.72831 |
| CRT490 | *Lophuromys dudui* | Muridae | Bomane Champ Nic2 | DRC | 2010 | 1.27374 | 23.72831 |
| CRT502 | *Lemniscomys striatus* | Muridae | Bomane Champ Nic2 | DRC | 2010 | 1.27374 | 23.72831 |
| CRT518 | *Praomys* sp. | Muridae | Bomane Ligne 1 Dudu ile | DRC | 2010 | 1.26534 | 23.74014 |
| CRT548 | *Praomys* sp. | Muridae | Bomane Ligne 1 Dudu ile | DRC | 2010 | 1.26534 | 23.74014 |
| CRT551 | *Lemniscomys striatus* | Muridae | | DRC | 2010 | | |
| CRT558 | *Praomys jacksoni* | Muridae | Bomane Ligne 2 Dudu ile | DRC | 2010 | 1.26573 | 23.74016 |
| CRT64 | *Lophuromys dudui* | Muridae | Yaikela Ligne 2 | DRC | 2010 | 0.82099 | 24.27678 |
| CRT682 | *Graphiurus* sp. 2* | Gliridae | Lieki paysans RD Dudu | DRC | 2010 | 0.67661 | 24.22865 |
| CRT74 | *Lophuromys dudui* | Muridae | Yaikela Ligne 2 | DRC | 2010 | 0.82099 | 24.27678 |
| EPU7 | *Lophuromys* sp. | Muridae | Mayali mingi | DRC | 2012 | 1.40646 | 28.56027 |
| KRT-210 | *Acomys* sp. | Muridae | Karatu | Tanzania | 2010 | -3.3 | 35.6 |
| TE5836 | *Lophuromys kilonzoi* | Muridae | Magamba | Tanzania | 2006 | -4.75 | 38.29 |
| MOZ002 | *Micaelamys namaquensis* | Muridae | Chimanimani | Mozambique | 2011 | -19.7105 | 33.0069 |
| MOZ068 | *Lophuromys machangui* | Muridae | Mt Mabu | Mozambique | 2011 | -16.3057 | 36.4241 |
| MOZ091 | *Lophuromys machangui* | Muridae | Mt Mabu | Mozambique | 2011 | -16.3057 | 36.4241 |
| MOZ094 | *Lophuromys machangui* | Muridae | Mt Mabu | Mozambique | 2011 | -16.3057 | 36.4241 |
| MOZ133 | *Lophuromys machangui* | Muridae | Mt Mabu | Mozambique | 2011 | -16.3086 | 36.4245 |
| MOZ134 | *Lophuromys machangui* | Muridae | Mt Mabu | Mozambique | 2011 | -16.3086 | 36.4245 |
| MOZ135 | *Lophuromys machangui* | Muridae | Mt Mabu | Mozambique | 2011 | -16.3086 | 36.4245 |
| MOZ241 | *Lophuromys machangui* | Muridae | Gile NP | Mozambique | 2011 | -16.7018 | 38.7984 |
| MOZ295 | *Lophuromys machangui* | Muridae | Mt Jesi | Mozambique | 2011 | -12.8675 | 35.185 |
| MOZ329 | *Lophuromys machangui* | Muridae | Gurue | Mozambique | 2011 | -15.7366 | 37.2063 |
| T8_345 | *Tachyoryctes splendens* | Spalacidae | Kilimanjaro-Tarakea | Tanzania | 2008 | -3.0569 | 37.5321 |
| T8_359 | *Heliophobius argenteocinereus* | Bathyergidae | Handei forest | Tanzania | 2008 | -5.02948 | 38.60332 |

| | | | | | | | |
|---|---|---|---|---|---|---|---|
| T8_510 | *Gerbilliscus vicinus* | Muridae | Shinyanga-Lubaga | Tanzania | 2008 | -3.6375 | 33.41723 |
| TZ20496 | *Lophuromys kilonzoi* | Muridae | Gologolo | Tanzania | 2007 | -4.8 | 38.26666667 |
| TZ20565 | *Rattus rattus* | Muridae | Emao | Tanzania | 2007 | -4.63 | 38.27 |
| TZ20612 | *Lophuromys kilonzoi* | Muridae | Kiranga | Tanzania | 2007 | -4.58 | 38.27 |
| TZ20620 | *Lophuromys kilonzoi* | Muridae | Kiranga | Tanzania | 2007 | -4.58 | 38.27 |
| TZ21205 | *Grammomys* sp. | Muridae | Magamba | Tanzania | 2009 | -4.75 | 38.28333333 |
| TZ21837 | *Lophuromys kilonzoi* | Muridae | Lushoto | Tanzania | 2009 | -4.8 | 38.3 |
| TZ25717 | *Acomys wilsoni* | Muridae | Mbwewe_site 1 | Tanzania | 2011 | -5.99493 | 38.23189 |
| TZ25757 | *Acomys wilsoni* | Muridae | Mihuga_site 2 | Tanzania | 2011 | -6.20882 | 38.53539 |

* The current nomenclature does not reliably delineate the genetic lineages in the *Graphiurus* genus. Based on the experts' opinion that co-authored this manuscript, *Graphiurus* individuals from Kenya are genetically different from those in the DRC. Although we cannot update the rodent species taxonomy within the scope of this manuscript, we will correspond *Graphiurus* sp. 1 to the Kenyan specimen and *Graphiurus* sp. 2 to the specimen originating from the DRC.

| | 1 | 2 | 3 | 4 | 5 | 6 | 7 | 8 | 9 | 10 | 11 | 12 | 13 | 14 | 15 | 16 | 17 | 18 | 19 | 20 | 21 | 22 | 23 | 24 | 25 | 26 | 27 | 28 | 29 | 30 | 31 | 32 | 33 | 34 | 35 | 36 | 37 | 38 | 39 | 40 | 41 | 42 | 43 | 44 | 45 | 46 | 47 | 48 | 49 | 50 | 51 | 52 | 53 | 54 | 55 | 56 | 57 | 58 | 59 | 60 |
|---|---|---|---|---|---|---|---|---|---|---|---|---|---|---|---|---|---|---|---|---|---|---|---|---|---|---|---|---|---|---|---|---|---|---|---|---|---|---|---|---|---|---|---|---|---|---|---|---|---|---|---|---|---|---|---|---|---|---|---|---|
| 1 MG599990_fish/rept/turtle_Gymnothorax_reticularis_China |
| 2 MG334001_fish/rept/turtle_Trachemys_scripta_USA_2008 | 0.629 |
| 3 MK737640_duck_Anas_platyrhynchos_China_2018 | 0.605 | 0.524 |
| 4 MN133813_mosquito_Culex_annulirostris_Australia_2018 | 0.626 | 0.58 | 0.531 |
| 5 MN062427_eagle_Haliaeetus_leucocephalus_USA_2002 | 0.598 | 0.584 | 0.517 | 0.451 |
| 6 MG599993_fish/rept/turtle_Protopterus_annectens_Nigeria | 0.622 | 0.524 | 0.563 | 0.601 | 0.556 |
| 7 MG599988_fish/rept/turtle_Goniurosaurus_luii_China | 0.605 | 0.605 | 0.598 | 0.587 | 0.573 | 0.563 |
| 8 MG599987_fish/rept/turtle_Teratoscincus_roborowskii_China | 0.636 | 0.626 | 0.584 | 0.626 | 0.619 | 0.612 | 0.476 |
| 9 MG599989_fish/rept/turtle_Hemidactylus_bowringii_China | 0.612 | 0.594 | 0.573 | 0.601 | 0.549 | 0.584 | 0.455 | 0.458 |
| 10 MG599991_fish/rept/turtle_Rhinobatos_hynnicephalus_China | 0.591 | 0.507 | 0.528 | 0.545 | 0.521 | 0.472 | 0.601 | 0.608 | 0.563 |
| 11 MG599992_fish/rept/turtle_Urolophus_aurantiacus_China | 0.647 | 0.524 | 0.538 | 0.577 | 0.549 | 0.483 | 0.594 | 0.612 | 0.577 | 0.437 |
| 12 KP641127_cattle_Bos_taurus_Germany_2014 | 0.587 | 0.517 | 0.503 | 0.535 | 0.542 | 0.521 | 0.563 | 0.566 | 0.524 | 0.503 | 0.517 |
| 13 KC551800_primate_Colobus_guereza_Uganda_2010 | 0.605 | 0.545 | 0.521 | 0.566 | 0.524 | 0.524 | 0.566 | 0.552 | 0.563 | 0.5 | 0.556 | 0.42 |
| 14 MH370348_rodent_Oligoryzomys_nigripes_Brazil_2012 | 0.608 | 0.563 | 0.556 | 0.584 | 0.563 | 0.514 | 0.601 | 0.573 | 0.57 | 0.542 | 0.545 | 0.455 | 0.451 |
| 15 KC411806_rodent_Rhabdomys_pumilio_South_Africa_2008 | 0.629 | 0.556 | 0.51 | 0.584 | 0.563 | 0.521 | 0.598 | 0.57 | 0.563 | 0.531 | 0.549 | 0.49 | 0.472 | 0.378 |
| 16 KY370092_rodent_Meriones_meridianus_China_2014 | 0.612 | 0.556 | 0.521 | 0.563 | 0.542 | 0.514 | 0.556 | 0.57 | 0.591 | 0.517 | 0.545 | 0.465 | 0.469 | 0.381 | 0.329 |
| 17 AF179612_primate_Saguinus_mystax_South_America_1995 | 0.584 | 0.552 | 0.549 | 0.545 | 0.507 | 0.545 | 0.556 | 0.57 | 0.531 | 0.503 | 0.514 | 0.472 | 0.465 | 0.517 | 0.549 | 0.524 |
| 18 MF775364_shrew_Suncus_murinus_China_2015 | 0.664 | 0.566 | 0.559 | 0.577 | 0.57 | 0.556 | 0.573 | 0.587 | 0.58 | 0.549 | 0.542 | 0.479 | 0.524 | 0.531 | 0.531 | 0.566 | 0.521 |
| 19 KC796077_bat_Hipposideros_vittatus_Kenya_2010 | 0.612 | 0.542 | 0.483 | 0.517 | 0.493 | 0.507 | 0.545 | 0.549 | 0.521 | 0.476 | 0.514 | 0.392 | 0.378 | 0.451 | 0.483 | 0.455 | 0.413 | 0.514 |
| 20 MN635449_possum_Trichosurus_vulpecula_Sydney | 0.601 | 0.573 | 0.531 | 0.577 | 0.556 | 0.556 | 0.594 | 0.587 | 0.57 | 0.514 | 0.542 | 0.441 | 0.476 | 0.528 | 0.524 | 0.545 | 0.444 | 0.517 | 0.451 |
| 21 KJ950939_rodent_Rattus_norvegicus_USA_2012 | 0.594 | 0.559 | 0.528 | 0.573 | 0.556 | 0.535 | 0.577 | 0.573 | 0.559 | 0.531 | 0.542 | 0.448 | 0.448 | 0.517 | 0.521 | 0.517 | 0.448 | 0.5 | 0.413 | 0.416 |
| 22 MH824542_sifaka_Propithecus_diadema_Madagascar_2013 | 0.598 | 0.535 | 0.528 | 0.566 | 0.507 | 0.514 | 0.573 | 0.552 | 0.549 | 0.514 | 0.514 | 0.444 | 0.43 | 0.493 | 0.514 | 0.524 | 0.43 | 0.542 | 0.406 | 0.402 | 0.448 |
| 23 MG211815_rodent_Spermophilus_dauricus_China_2015 | 0.629 | 0.563 | 0.51 | 0.577 | 0.528 | 0.507 | 0.573 | 0.566 | 0.552 | 0.507 | 0.524 | 0.434 | 0.437 | 0.469 | 0.51 | 0.479 | 0.451 | 0.497 | 0.388 | 0.43 | 0.371 | 0.364 |
| 24 MN564789_CRT682_rodent_Graphiurus_cf._kelleni_DRC_2010 | 0.633 | 0.573 | 0.538 | 0.591 | 0.531 | 0.545 | 0.605 | 0.584 | 0.577 | 0.521 | 0.535 | 0.462 | 0.479 | 0.479 | 0.535 | 0.497 | 0.497 | 0.528 | 0.451 | 0.465 | 0.43 | 0.42 | 0.388 |
| 25 MN587695_TA085_rodent_Lophuromys_stanleyi_Tanzania_2013 | 0.619 | 0.545 | 0.503 | 0.58 | 0.566 | 0.483 | 0.58 | 0.552 | 0.545 | 0.497 | 0.517 | 0.476 | 0.444 | 0.465 | 0.472 | 0.486 | 0.476 | 0.472 | 0.448 | 0.469 | 0.434 | 0.458 | 0.367 | 0.409 |
| 26 MN587668_MOZ133_rodent_Lophuromys_machangui_Mozambique_2011 | 0.626 | 0.556 | 0.514 | 0.58 | 0.573 | 0.49 | 0.58 | 0.563 | 0.542 | 0.528 | 0.521 | 0.479 | 0.437 | 0.465 | 0.486 | 0.503 | 0.497 | 0.483 | 0.444 | 0.472 | 0.441 | 0.469 | 0.392 | 0.423 | 0.101 |
| 27 KJ950938_rodent_Rattus_norvegicus_USA_2013 | 0.619 | 0.559 | 0.542 | 0.566 | 0.538 | 0.531 | 0.577 | 0.58 | 0.535 | 0.524 | 0.545 | 0.448 | 0.416 | 0.476 | 0.521 | 0.51 | 0.444 | 0.51 | 0.402 | 0.413 | 0.385 | 0.374 | 0.353 | 0.434 | 0.409 | 0.416 |
| 28 KY370095_rodent_Dipus_sagitta_China_2014 | 0.629 | 0.584 | 0.535 | 0.587 | 0.556 | 0.538 | 0.584 | 0.587 | 0.58 | 0.514 | 0.538 | 0.458 | 0.409 | 0.51 | 0.486 | 0.465 | 0.479 | 0.51 | 0.441 | 0.444 | 0.395 | 0.42 | 0.388 | 0.423 | 0.399 | 0.427 | 0.381 |
| 29 MN555567_TZ25757_rodent_Acomys_wilsoni_Tanzania_2011 | 0.636 | 0.577 | 0.531 | 0.598 | 0.545 | 0.535 | 0.601 | 0.584 | 0.563 | 0.51 | 0.549 | 0.451 | 0.462 | 0.514 | 0.51 | 0.479 | 0.469 | 0.521 | 0.462 | 0.462 | 0.416 | 0.413 | 0.381 | 0.409 | 0.448 | 0.465 | 0.378 | 0.357 |
| 30 MG822666_rodent_Proechimys_semispinosus_Panama_2014 | 0.664 | 0.605 | 0.559 | 0.615 | 0.531 | 0.573 | 0.605 | 0.605 | 0.566 | 0.559 | 0.552 | 0.493 | 0.476 | 0.497 | 0.535 | 0.521 | 0.5 | 0.517 | 0.479 | 0.476 | 0.42 | 0.448 | 0.43 | 0.406 | 0.469 | 0.472 | 0.381 | 0.395 | 0.322 |
| 31 MN564790_ETH674_rodent_Stenocephalemys_albipes_Ethiopia_2012 | 0.643 | 0.584 | 0.57 | 0.598 | 0.542 | 0.552 | 0.619 | 0.601 | 0.584 | 0.542 | 0.535 | 0.493 | 0.462 | 0.503 | 0.514 | 0.49 | 0.49 | 0.538 | 0.451 | 0.469 | 0.385 | 0.42 | 0.402 | 0.381 | 0.444 | 0.441 | 0.371 | 0.357 | 0.273 | 0.318 |
| 32 MG600413_rodent_Rhizomys_pruinosus_Vietnam_2015 | 0.643 | 0.591 | 0.549 | 0.601 | 0.566 | 0.559 | 0.598 | 0.573 | 0.563 | 0.559 | 0.552 | 0.476 | 0.472 | 0.493 | 0.507 | 0.49 | 0.476 | 0.535 | 0.458 | 0.472 | 0.395 | 0.427 | 0.378 | 0.413 | 0.43 | 0.448 | 0.364 | 0.36 | 0.287 | 0.322 | 0.311 |
| 33 KC815310_rodent_Peromyscus_maniculatus_USA_2008 | 0.633 | 0.587 | 0.542 | 0.598 | 0.566 | 0.535 | 0.573 | 0.573 | 0.538 | 0.538 | 0.514 | 0.458 | 0.465 | 0.5 | 0.51 | 0.503 | 0.462 | 0.521 | 0.441 | 0.451 | 0.381 | 0.406 | 0.367 | 0.395 | 0.43 | 0.441 | 0.364 | 0.346 | 0.255 | 0.35 | 0.266 | 0.287 |
| 34 KY370094_rodent_Neodon_clarkei_China_2014 | 0.65 | 0.556 | 0.524 | 0.594 | 0.521 | 0.531 | 0.605 | 0.594 | 0.566 | 0.51 | 0.514 | 0.455 | 0.437 | 0.49 | 0.497 | 0.493 | 0.444 | 0.528 | 0.434 | 0.434 | 0.395 | 0.406 | 0.416 | 0.402 | 0.437 | 0.458 | 0.371 | 0.336 | 0.262 | 0.301 | 0.287 | 0.301 | 0.266 |
| 35 KC411784_rodent_Myodes_glareolus_Netherlands_2007 | 0.633 | 0.559 | 0.517 | 0.612 | 0.531 | 0.549 | 0.615 | 0.612 | 0.58 | 0.507 | 0.531 | 0.465 | 0.462 | 0.503 | 0.507 | 0.514 | 0.486 | 0.542 | 0.455 | 0.43 | 0.42 | 0.402 | 0.413 | 0.406 | 0.441 | 0.469 | 0.378 | 0.374 | 0.259 | 0.322 | 0.304 | 0.325 | 0.269 | 0.22 |
| 36 KC796074_bat_Hipposideros_vittatus_Kenya_2011 | 0.629 | 0.49 | 0.517 | 0.545 | 0.563 | 0.503 | 0.587 | 0.563 | 0.542 | 0.483 | 0.503 | 0.486 | 0.472 | 0.514 | 0.497 | 0.507 | 0.507 | 0.528 | 0.462 | 0.538 | 0.514 | 0.517 | 0.49 | 0.51 | 0.472 | 0.472 | 0.49 | 0.503 | 0.531 | 0.535 | 0.524 | 0.517 | 0.524 | 0.507 | 0.524 |
| 37 KC796078_bat_Otomops_martiensseni_Kenya_2011 | 0.615 | 0.497 | 0.493 | 0.545 | 0.573 | 0.493 | 0.577 | 0.528 | 0.528 | 0.507 | 0.514 | 0.469 | 0.507 | 0.542 | 0.514 | 0.524 | 0.517 | 0.51 | 0.479 | 0.545 | 0.472 | 0.49 | 0.483 | 0.503 | 0.462 | 0.469 | 0.486 | 0.486 | 0.476 | 0.514 | 0.5 | 0.486 | 0.486 | 0.483 | 0.497 | 0.353 |
| 38 KP325401_horse_Equus_ferus_USA_2011 | 0.587 | 0.497 | 0.503 | 0.556 | 0.58 | 0.514 | 0.573 | 0.559 | 0.528 | 0.483 | 0.507 | 0.483 | 0.479 | 0.503 | 0.503 | 0.531 | 0.483 | 0.514 | 0.448 | 0.524 | 0.465 | 0.486 | 0.476 | 0.528 | 0.469 | 0.458 | 0.493 | 0.483 | 0.503 | 0.524 | 0.524 | 0.51 | 0.497 | 0.486 | 0.503 | 0.357 | 0.364 |
| 39 EF108306_human_Homo_sapiens | 0.619 | 0.514 | 0.542 | 0.535 | 0.556 | 0.472 | 0.58 | 0.577 | 0.552 | 0.486 | 0.556 | 0.493 | 0.455 | 0.552 | 0.556 | 0.552 | 0.476 | 0.545 | 0.451 | 0.535 | 0.5 | 0.476 | 0.5 | 0.566 | 0.486 | 0.497 | 0.514 | 0.514 | 0.517 | 0.549 | 0.552 | 0.549 | 0.528 | 0.5 | 0.535 | 0.416 | 0.42 | 0.336 |
| 40 NC_009823_human_Homo_sapiens | 0.622 | 0.517 | 0.538 | 0.538 | 0.556 | 0.486 | 0.573 | 0.58 | 0.552 | 0.493 | 0.559 | 0.476 | 0.472 | 0.528 | 0.514 | 0.542 | 0.479 | 0.542 | 0.448 | 0.524 | 0.514 | 0.49 | 0.493 | 0.563 | 0.493 | 0.507 | 0.514 | 0.521 | 0.521 | 0.552 | 0.549 | 0.514 | 0.531 | 0.528 | 0.517 | 0.406 | 0.427 | 0.329 | 0.136 |
| 41 NC_009824_human_Homo_sapiens | 0.615 | 0.51 | 0.521 | 0.531 | 0.552 | 0.49 | 0.58 | 0.584 | 0.563 | 0.49 | 0.549 | 0.507 | 0.465 | 0.524 | 0.524 | 0.542 | 0.472 | 0.542 | 0.448 | 0.549 | 0.517 | 0.493 | 0.5 | 0.556 | 0.469 | 0.486 | 0.517 | 0.507 | 0.507 | 0.559 | 0.545 | 0.524 | 0.528 | 0.528 | 0.521 | 0.402 | 0.413 | 0.308 | 0.164 | 0.154 |
| 42 NC_009826_human_Homo_sapiens | 0.633 | 0.517 | 0.528 | 0.545 | 0.556 | 0.507 | 0.58 | 0.587 | 0.58 | 0.507 | 0.545 | 0.5 | 0.462 | 0.535 | 0.524 | 0.528 | 0.486 | 0.538 | 0.455 | 0.545 | 0.514 | 0.486 | 0.493 | 0.57 | 0.497 | 0.503 | 0.521 | 0.514 | 0.51 | 0.542 | 0.531 | 0.514 | 0.521 | 0.5 | 0.507 | 0.406 | 0.448 | 0.343 | 0.143 | 0.143 | 0.154 |
| 43 NC_009827_human_Homo_sapiens | 0.626 | 0.535 | 0.535 | 0.552 | 0.552 | 0.483 | 0.594 | 0.566 | 0.556 | 0.503 | 0.559 | 0.5 | 0.476 | 0.531 | 0.514 | 0.521 | 0.497 | 0.535 | 0.458 | 0.528 | 0.503 | 0.479 | 0.493 | 0.542 | 0.483 | 0.483 | 0.517 | 0.503 | 0.503 | 0.549 | 0.521 | 0.517 | 0.514 | 0.507 | 0.517 | 0.434 | 0.416 | 0.332 | 0.154 | 0.157 | 0.189 | 0.147 |
| 44 NC_009825_human_Homo_sapiens | 0.622 | 0.51 | 0.524 | 0.556 | 0.563 | 0.497 | 0.584 | 0.577 | 0.552 | 0.507 | 0.542 | 0.49 | 0.472 | 0.517 | 0.524 | 0.524 | 0.479 | 0.542 | 0.451 | 0.535 | 0.5 | 0.483 | 0.486 | 0.556 | 0.479 | 0.479 | 0.507 | 0.507 | 0.497 | 0.524 | 0.521 | 0.503 | 0.5 | 0.507 | 0.497 | 0.413 | 0.437 | 0.329 | 0.143 | 0.154 | 0.161 | 0.143 | 0.133 |
| 45 M62321_HCV_reference | 0.626 | 0.521 | 0.524 | 0.542 | 0.556 | 0.472 | 0.58 | 0.566 | 0.549 | 0.479 | 0.528 | 0.493 | 0.451 | 0.51 | 0.497 | 0.51 | 0.476 | 0.535 | 0.423 | 0.521 | 0.493 | 0.462 | 0.476 | 0.542 | 0.469 | 0.486 | 0.493 | 0.5 | 0.493 | 0.521 | 0.507 | 0.507 | 0.497 | 0.486 | 0.5 | 0.378 | 0.395 | 0.308 | 0.136 | 0.143 | 0.171 | 0.14 | 0.122 | 0.129 |
| 46 MG599997_fish/rept/turtle_Chimaera_sp_China | 0.594 | 0.507 | 0.497 | 0.57 | 0.559 | 0.483 | 0.608 | 0.615 | 0.577 | 0.43 | 0.448 | 0.483 | 0.528 | 0.517 | 0.528 | 0.514 | 0.514 | 0.559 | 0.5 | 0.517 | 0.524 | 0.49 | 0.514 | 0.5 | 0.517 | 0.538 | 0.517 | 0.528 | 0.514 | 0.538 | 0.521 | 0.528 | 0.507 | 0.503 | 0.531 | 0.493 | 0.5 | 0.497 | 0.51 | 0.493 | 0.503 | 0.493 | 0.5 | 0.5 | 0.483 |
| 47 MG599996_fish/rept/turtle_Chimaera_sp_China | 0.619 | 0.51 | 0.497 | 0.556 | 0.545 | 0.493 | 0.587 | 0.619 | 0.58 | 0.451 | 0.476 | 0.455 | 0.531 | 0.497 | 0.486 | 0.51 | 0.521 | 0.535 | 0.462 | 0.528 | 0.521 | 0.476 | 0.503 | 0.521 | 0.542 | 0.538 | 0.538 | 0.531 | 0.51 | 0.538 | 0.542 | 0.521 | 0.528 | 0.51 | 0.5 | 0.497 | 0.507 | 0.503 | 0.514 | 0.5 | 0.507 | 0.503 | 0.514 | 0.514 | 0.507 | 0.371 |
| 48 MG599995_fish/rept/turtle_Squalus_brevirostris_China | 0.594 | 0.535 | 0.5 | 0.573 | 0.531 | 0.503 | 0.598 | 0.619 | 0.549 | 0.476 | 0.458 | 0.497 | 0.57 | 0.563 | 0.552 | 0.545 | 0.51 | 0.556 | 0.503 | 0.531 | 0.57 | 0.521 | 0.549 | 0.538 | 0.545 | 0.563 | 0.531 | 0.58 | 0.521 | 0.57 | 0.573 | 0.549 | 0.521 | 0.538 | 0.531 | 0.49 | 0.497 | 0.507 | 0.535 | 0.524 | 0.524 | 0.531 | 0.521 | 0.535 | 0.521 | 0.448 | 0.437 |
| 49 MG599994_fish/rept/turtle_Mustelus_manazo_China | 0.629 | 0.524 | 0.507 | 0.57 | 0.566 | 0.514 | 0.594 | 0.615 | 0.598 | 0.43 | 0.479 | 0.531 | 0.521 | 0.566 | 0.559 | 0.552 | 0.528 | 0.535 | 0.521 | 0.514 | 0.545 | 0.507 | 0.517 | 0.559 | 0.531 | 0.545 | 0.535 | 0.531 | 0.535 | 0.58 | 0.559 | 0.57 | 0.535 | 0.528 | 0.542 | 0.483 | 0.479 | 0.476 | 0.507 | 0.524 | 0.51 | 0.531 | 0.528 | 0.521 | 0.497 | 0.413 | 0.437 | 0.451 |
| 50 MG599999_fish/rept/turtle_Pelodiscus_sinensis_China | 0.619 | 0.493 | 0.5 | 0.535 | 0.531 | 0.503 | 0.591 | 0.591 | 0.584 | 0.451 | 0.493 | 0.49 | 0.514 | 0.559 | 0.521 | 0.51 | 0.517 | 0.552 | 0.493 | 0.542 | 0.535 | 0.503 | 0.507 | 0.542 | 0.514 | 0.521 | 0.521 | 0.514 | 0.524 | 0.545 | 0.545 | 0.556 | 0.542 | 0.51 | 0.507 | 0.458 | 0.458 | 0.469 | 0.497 | 0.503 | 0.486 | 0.5 | 0.479 | 0.507 | 0.479 | 0.476 | 0.448 | 0.483 | 0.462 |
| 51 MG600000_fish/rept/turtle_Mauremys_megalocephala_China | 0.591 | 0.524 | 0.497 | 0.57 | 0.524 | 0.497 | 0.563 | 0.601 | 0.563 | 0.507 | 0.514 | 0.486 | 0.5 | 0.545 | 0.559 | 0.535 | 0.497 | 0.573 | 0.465 | 0.538 | 0.545 | 0.5 | 0.462 | 0.538 | 0.483 | 0.507 | 0.486 | 0.514 | 0.528 | 0.556 | 0.552 | 0.549 | 0.521 | 0.51 | 0.51 | 0.469 | 0.486 | 0.497 | 0.465 | 0.476 | 0.441 | 0.469 | 0.469 | 0.472 | 0.469 | 0.479 | 0.476 | 0.483 | 0.479 | 0.308 |
| 52 KY370091_rodent_Allactaga_sibirica_China_2014 | 0.584 | 0.479 | 0.486 | 0.528 | 0.545 | 0.514 | 0.577 | 0.556 | 0.531 | 0.514 | 0.493 | 0.483 | 0.49 | 0.535 | 0.542 | 0.514 | 0.535 | 0.545 | 0.465 | 0.528 | 0.507 | 0.503 | 0.507 | 0.531 | 0.51 | 0.51 | 0.472 | 0.514 | 0.517 | 0.563 | 0.535 | 0.521 | 0.517 | 0.49 | 0.524 | 0.458 | 0.441 | 0.458 | 0.469 | 0.462 | 0.486 | 0.465 | 0.455 | 0.465 | 0.462 | 0.462 | 0.476 | 0.486 | 0.462 | 0.458 | 0.465 |
| 53 KC411777_rodent_Myodes_glareolus_Germany_2010 | 0.577 | 0.476 | 0.493 | 0.549 | 0.51 | 0.493 | 0.573 | 0.57 | 0.549 | 0.462 | 0.493 | 0.448 | 0.493 | 0.503 | 0.497 | 0.493 | 0.469 | 0.552 | 0.448 | 0.517 | 0.51 | 0.451 | 0.479 | 0.462 | 0.469 | 0.5 | 0.493 | 0.5 | 0.483 | 0.531 | 0.5 | 0.507 | 0.517 | 0.472 | 0.497 | 0.43 | 0.416 | 0.448 | 0.423 | 0.43 | 0.423 | 0.437 | 0.434 | 0.437 | 0.423 | 0.434 | 0.455 | 0.5 | 0.486 | 0.42 | 0.406 | 0.388 |
| 54 MN564792_TA132_rodent_Praomys_jacksoni_Tanzania_2013 | 0.654 | 0.472 | 0.5 | 0.552 | 0.556 | 0.486 | 0.584 | 0.612 | 0.58 | 0.472 | 0.538 | 0.486 | 0.5 | 0.524 | 0.521 | 0.503 | 0.514 | 0.545 | 0.493 | 0.552 | 0.552 | 0.507 | 0.486 | 0.545 | 0.497 | 0.517 | 0.517 | 0.514 | 0.524 | 0.57 | 0.552 | 0.542 | 0.542 | 0.521 | 0.531 | 0.448 | 0.472 | 0.479 | 0.472 | 0.476 | 0.465 | 0.462 | 0.458 | 0.465 | 0.458 | 0.483 | 0.479 | 0.542 | 0.472 | 0.444 | 0.458 | 0.451 | 0.308 |
| 55 MN587672_TA275_rodent_Lophuromys_laticeps_Tanzania_2013 | 0.643 | 0.49 | 0.528 | 0.587 | 0.559 | 0.5 | 0.584 | 0.605 | 0.556 | 0.49 | 0.552 | 0.49 | 0.51 | 0.549 | 0.563 | 0.549 | 0.507 | 0.584 | 0.497 | 0.531 | 0.545 | 0.493 | 0.483 | 0.559 | 0.531 | 0.535 | 0.51 | 0.517 | 0.524 | 0.556 | 0.563 | 0.542 | 0.556 | 0.493 | 0.531 | 0.486 | 0.472 | 0.486 | 0.465 | 0.476 | 0.462 | 0.462 | 0.441 | 0.469 | 0.472 | 0.5 | 0.5 | 0.535 | 0.5 | 0.462 | 0.434 | 0.455 | 0.318 | 0.217 |
| 56 MN587655_CRT352-B_rodent_Lophuromys_dudui_DRC_2010 | 0.629 | 0.486 | 0.535 | 0.58 | 0.556 | 0.51 | 0.591 | 0.619 | 0.552 | 0.503 | 0.552 | 0.507 | 0.51 | 0.545 | 0.552 | 0.552 | 0.517 | 0.57 | 0.507 | 0.542 | 0.556 | 0.49 | 0.486 | 0.559 | 0.528 | 0.538 | 0.517 | 0.521 | 0.535 | 0.566 | 0.559 | 0.556 | 0.559 | 0.521 | 0.556 | 0.469 | 0.479 | 0.479 | 0.462 | 0.469 | 0.465 | 0.455 | 0.434 | 0.465 | 0.465 | 0.486 | 0.5 | 0.528 | 0.493 | 0.462 | 0.43 | 0.441 | 0.315 | 0.206 | 0.077 |
| 57 MN587683_TA529-B_rodent_Lophuromys_machangui_Tanzania_2013 | 0.636 | 0.493 | 0.535 | 0.58 | 0.552 | 0.517 | 0.587 | 0.605 | 0.563 | 0.497 | 0.57 | 0.493 | 0.51 | 0.552 | 0.552 | 0.549 | 0.51 | 0.58 | 0.493 | 0.538 | 0.556 | 0.5 | 0.49 | 0.566 | 0.528 | 0.538 | 0.51 | 0.514 | 0.528 | 0.566 | 0.559 | 0.552 | 0.563 | 0.521 | 0.545 | 0.479 | 0.486 | 0.483 | 0.472 | 0.462 | 0.455 | 0.462 | 0.444 | 0.465 | 0.465 | 0.514 | 0.514 | 0.538 | 0.5 | 0.465 | 0.444 | 0.448 | 0.322 | 0.21 | 0.084 | 0.059 |
| 58 MN587662_CRT64-B_rodent_Lophuromys_dudui_DRC_2010 | 0.647 | 0.493 | 0.531 | 0.57 | 0.559 | 0.503 | 0.591 | 0.612 | 0.549 | 0.503 | 0.556 | 0.493 | 0.51 | 0.552 | 0.556 | 0.552 | 0.514 | 0.58 | 0.5 | 0.531 | 0.563 | 0.493 | 0.497 | 0.57 | 0.517 | 0.521 | 0.503 | 0.521 | 0.531 | 0.577 | 0.566 | 0.556 | 0.566 | 0.521 | 0.538 | 0.49 | 0.465 | 0.479 | 0.465 | 0.462 | 0.451 | 0.462 | 0.427 | 0.462 | 0.462 | 0.497 | 0.503 | 0.538 | 0.49 | 0.455 | 0.434 | 0.434 | 0.329 | 0.224 | 0.077 | 0.063 | 0.066 |
| 59 MN587670_MOZ329-B_rodent_Lophuromys_machangui_Mozambique_201[illegible] | 0.64 | 0.469 | 0.538 | 0.573 | 0.563 | 0.497 | 0.584 | 0.619 | 0.556 | 0.49 | 0.545 | 0.493 | 0.521 | 0.549 | 0.552 | 0.556 | 0.521 | 0.57 | 0.507 | 0.545 | 0.566 | 0.503 | 0.493 | 0.556 | 0.521 | 0.524 | 0.517 | 0.521 | 0.531 | 0.573 | 0.573 | 0.552 | 0.559 | 0.51 | 0.549 | 0.462 | 0.476 | 0.479 | 0.472 | 0.469 | 0.455 | 0.465 | 0.451 | 0.469 | 0.476 | 0.486 | 0.503 | 0.528 | 0.483 | 0.451 | 0.448 | 0.43 | 0.322 | 0.196 | 0.073 | 0.063 | 0.07 | 0.066 |
| 60 MN587666_CRT74_rodent_Lophuromys_dudui_DRC_2010 | 0.636 | 0.493 | 0.535 | 0.577 | 0.563 | 0.507 | 0.584 | 0.612 | 0.552 | 0.497 | 0.552 | 0.5 | 0.517 | 0.559 | 0.559 | 0.556 | 0.507 | 0.573 | 0.497 | 0.545 | 0.559 | 0.493 | 0.5 | 0.563 | 0.535 | 0.538 | 0.51 | 0.524 | 0.528 | 0.559 | 0.559 | 0.556 | 0.552 | 0.517 | 0.552 | 0.472 | 0.465 | 0.472 | 0.472 | 0.469 | 0.451 | 0.469 | 0.441 | 0.465 | 0.469 | 0.493 | 0.5 | 0.521 | 0.486 | 0.458 | 0.434 | 0.444 | 0.329 | 0.217 | 0.077 | 0.052 | 0.066 | 0.059 | 0.059 |

Estimates of Evolutionary Divergence between Sequences, NS3 region

| | 1 | 2 | 3 | 4 | 5 | 6 | 7 | 8 | 9 | 10 | 11 | 12 | 13 | 14 | 15 | 16 | 17 | 18 | 19 | 20 | 21 | 22 | 23 | 24 | 25 | 26 | 27 | 28 | 29 | 30 | 31 | 32 | 33 | 34 | 35 | 36 | 37 | 38 | 39 | 40 | 41 | 42 | 43 | 44 | 45 | 46 | 47 | 48 | 49 | 50 | 51 | 52 | 53 | 54 | 55 | 56 | 57 | 58 | 59 | 60 |
|---|---|---|---|---|---|---|---|---|---|---|---|---|---|---|---|---|---|---|---|---|---|---|---|---|---|---|---|---|---|---|---|---|---|---|---|---|---|---|---|---|---|---|---|---|---|---|---|---|---|---|---|---|---|---|---|---|---|---|---|---|
| 1 MG599990_fish/rept/turtle_Gymnothorax_reticularis_China | | | | | | | | | | | | | | | | | | | | | | | | | | | | | | | | | | | | | | | | | | | | | | | | | | | | | | | | | | | | |
| 2 MG334001_fish/rept/turtle_Trachemys_scripta_USA_2008 | 0.714 | | | | | | | | | | | | | | | | | | | | | | | | | | | | | | | | | | | | | | | | | | | | | | | | | | | | | | | | | | | |
| 3 MK737640_duck_Anas_platyrhynchos_China_2018 | 0.729 | 0.617 | | | | | | | | | | | | | | | | | | | | | | | | | | | | | | | | | | | | | | | | | | | | | | | | | | | | | | | | | | |
| 4 MN133813_mosquito_Culex_annulirostris_Australia_2018 | 0.752 | 0.624 | 0.489 | | | | | | | | | | | | | | | | | | | | | | | | | | | | | | | | | | | | | | | | | | | | | | | | | | | | | | | | |
| 5 MN062427_eagle_Haliaeetus_leucocephalus_USA_2002 | 0.737 | 0.699 | 0.444 | 0.489 | | | | | | | | | | | | | | | | | | | | | | | | | | | | | | | | | | | | | | | | | | | | | | | | | | | | | | | |
| 6 MG599993_fish/rept/turtle_Protopterus_annectens_Nigeria | 0.684 | 0.639 | 0.586 | 0.579 | 0.602 | | | | | | | | | | | | | | | | | | | | | | | | | | | | | | | | | | | | | | | | | | | | | | | | | | | | | |
| 7 MG599988_fish/rept/turtle_Goniurosaurus_luii_China | 0.722 | 0.632 | 0.526 | 0.586 | 0.549 | 0.564 | | | | | | | | | | | | | | | | | | | | | | | | | | | | | | | | | | | | | | | | | | | | | | | | | | | |
| 8 MG599987_fish/rept/turtle_Teratoscincus_roborowskii_China | 0.752 | 0.594 | 0.489 | 0.511 | 0.534 | 0.556 | 0.444 | | | | | | | | | | | | | | | | | | | | | | | | | | | | | | | | | | | | | | | | | | | | | | | | | |
| 9 MG599989_fish/rept/turtle_Hemidactylus_bowringii_China | 0.729 | 0.632 | 0.526 | 0.564 | 0.541 | 0.549 | 0.361 | 0.429 | | | | | | | | | | | | | | | | | | | | | | | | | | | | | | | | | | | | | | | | | | | | | | | | |
| 10 MG599991_fish/rept/turtle_Rhinobatos_hynnicephalus_China | 0.707 | 0.586 | 0.586 | 0.609 | 0.594 | 0.526 | 0.511 | 0.564 | 0.489 | | | | | | | | | | | | | | | | | | | | | | | | | | | | | | | | | | | | | | | | | | | | | | | |
| 11 MG599992_fish/rept/turtle_Urolophus_aurantiacus_China | 0.744 | 0.632 | 0.594 | 0.586 | 0.594 | 0.586 | 0.556 | 0.579 | 0.549 | 0.489 | | | | | | | | | | | | | | | | | | | | | | | | | | | | | | | | | | | | | | | | | | | | | | |
| 12 KP641127_cattle_Bos_taurus_Germany_2014 | 0.669 | 0.586 | 0.594 | 0.602 | 0.594 | 0.504 | 0.571 | 0.564 | 0.617 | 0.564 | 0.579 | | | | | | | | | | | | | | | | | | | | | | | | | | | | | | | | | | | | | | | | | | | | | |
| 13 KC551800_primate_Colobus_guereza_Uganda_2010 | 0.707 | 0.617 | 0.541 | 0.602 | 0.594 | 0.496 | 0.534 | 0.534 | 0.556 | 0.556 | 0.571 | 0.466 | | | | | | | | | | | | | | | | | | | | | | | | | | | | | | | | | | | | | | | | | | | | |
| 14 MH370348_rodent_Oligoryzomys_nigripes_Brazil_2012 | 0.722 | 0.677 | 0.526 | 0.579 | 0.534 | 0.466 | 0.556 | 0.556 | 0.564 | 0.534 | 0.586 | 0.451 | 0.451 | | | | | | | | | | | | | | | | | | | | | | | | | | | | | | | | | | | | | | | | | | | |
| 15 KC411806_rodent_Rhabdomys_pumilio_South_Africa_2008 | 0.662 | 0.639 | 0.556 | 0.571 | 0.496 | 0.496 | 0.534 | 0.549 | 0.549 | 0.541 | 0.556 | 0.444 | 0.421 | 0.293 | | | | | | | | | | | | | | | | | | | | | | | | | | | | | | | | | | | | | | | | | | |
| 16 KY370092_rodent_Meriones_meridianus_China_2014 | 0.684 | 0.639 | 0.549 | 0.564 | 0.519 | 0.504 | 0.541 | 0.526 | 0.586 | 0.534 | 0.594 | 0.459 | 0.481 | 0.301 | 0.293 | | | | | | | | | | | | | | | | | | | | | | | | | | | | | | | | | | | | | | | | | |
| 17 AF179612_primate_Saguinus_mystax_South_America_1995 | 0.722 | 0.662 | 0.602 | 0.594 | 0.564 | 0.624 | 0.594 | 0.586 | 0.594 | 0.564 | 0.647 | 0.534 | 0.496 | 0.481 | 0.466 | 0.481 | | | | | | | | | | | | | | | | | | | | | | | | | | | | | | | | | | | | | | | | |
| 18 MF775364_shrew_Suncus_murinus_China_2015 | 0.722 | 0.639 | 0.564 | 0.602 | 0.564 | 0.564 | 0.579 | 0.556 | 0.586 | 0.549 | 0.579 | 0.466 | 0.346 | 0.429 | 0.436 | 0.459 | 0.489 | | | | | | | | | | | | | | | | | | | | | | | | | | | | | | | | | | | | | | | |
| 19 KC796077_bat_Hipposideros_vittatus_Kenya_2010 | 0.714 | 0.617 | 0.579 | 0.594 | 0.594 | 0.586 | 0.549 | 0.586 | 0.579 | 0.586 | 0.609 | 0.534 | 0.504 | 0.489 | 0.489 | 0.526 | 0.504 | 0.526 | | | | | | | | | | | | | | | | | | | | | | | | | | | | | | | | | | | | | | |
| 20 MN635449_possum_Trichosurus_vulpecula_Sydney | 0.662 | 0.639 | 0.556 | 0.602 | 0.571 | 0.511 | 0.564 | 0.526 | 0.564 | 0.556 | 0.564 | 0.436 | 0.459 | 0.368 | 0.376 | 0.398 | 0.496 | 0.436 | 0.541 | | | | | | | | | | | | | | | | | | | | | | | | | | | | | | | | | | | | | |
| 21 KJ950939_rodent_Rattus_norvegicus_USA_2012 | 0.654 | 0.662 | 0.586 | 0.586 | 0.586 | 0.504 | 0.571 | 0.526 | 0.579 | 0.534 | 0.564 | 0.391 | 0.421 | 0.429 | 0.414 | 0.459 | 0.504 | 0.436 | 0.504 | 0.353 | | | | | | | | | | | | | | | | | | | | | | | | | | | | | | | | | | | | |
| 22 MH824542_sifaka_Propithecus_diadema_Madagascar_2013 | 0.714 | 0.662 | 0.586 | 0.586 | 0.586 | 0.496 | 0.602 | 0.594 | 0.609 | 0.541 | 0.632 | 0.421 | 0.451 | 0.429 | 0.436 | 0.451 | 0.526 | 0.414 | 0.511 | 0.376 | 0.353 | | | | | | | | | | | | | | | | | | | | | | | | | | | | | | | | | | | |
| 23 MG211815_rodent_Spermophilus_dauricus_China_2015 | 0.684 | 0.617 | 0.534 | 0.564 | 0.549 | 0.489 | 0.586 | 0.504 | 0.579 | 0.556 | 0.586 | 0.414 | 0.414 | 0.331 | 0.383 | 0.406 | 0.451 | 0.368 | 0.451 | 0.346 | 0.316 | 0.361 | | | | | | | | | | | | | | | | | | | | | | | | | | | | | | | | | | |
| 24 MN564789_CRT682_rodent_Graphiurus_cf._kelleni_DRC_2010 | 0.707 | 0.654 | 0.609 | 0.609 | 0.586 | 0.444 | 0.541 | 0.556 | 0.564 | 0.504 | 0.579 | 0.406 | 0.421 | 0.368 | 0.406 | 0.421 | 0.496 | 0.414 | 0.496 | 0.323 | 0.286 | 0.346 | 0.293 | | | | | | | | | | | | | | | | | | | | | | | | | | | | | | | | | |
| 25 MN587695_TA085_rodent_Lophuromys_stanleyi_Tanzania_2013 | 0.669 | 0.647 | 0.571 | 0.609 | 0.556 | 0.541 | 0.579 | 0.534 | 0.617 | 0.511 | 0.602 | 0.391 | 0.459 | 0.451 | 0.444 | 0.474 | 0.504 | 0.414 | 0.534 | 0.376 | 0.301 | 0.353 | 0.323 | 0.338 | | | | | | | | | | | | | | | | | | | | | | | | | | | | | | | | |
| 26 MN587668_MOZ133_rodent_Lophuromys_machangui_Mozambique_2011 | 0.662 | 0.624 | 0.564 | 0.617 | 0.564 | 0.534 | 0.571 | 0.541 | 0.617 | 0.519 | 0.594 | 0.406 | 0.436 | 0.459 | 0.429 | 0.489 | 0.519 | 0.414 | 0.549 | 0.383 | 0.301 | 0.376 | 0.338 | 0.346 | 0.038 | | | | | | | | | | | | | | | | | | | | | | | | | | | | | | | |
| 27 KJ950938_rodent_Rattus_norvegicus_USA_2013 | 0.684 | 0.639 | 0.594 | 0.579 | 0.564 | 0.459 | 0.564 | 0.534 | 0.594 | 0.534 | 0.571 | 0.406 | 0.429 | 0.421 | 0.398 | 0.459 | 0.429 | 0.421 | 0.474 | 0.368 | 0.293 | 0.376 | 0.271 | 0.301 | 0.301 | 0.308 | | | | | | | | | | | | | | | | | | | | | | | | | | | | | | |
| 28 KY370095_rodent_Dipus_sagitta_China_2014 | 0.692 | 0.617 | 0.586 | 0.586 | 0.594 | 0.534 | 0.556 | 0.549 | 0.594 | 0.534 | 0.594 | 0.398 | 0.444 | 0.406 | 0.429 | 0.459 | 0.444 | 0.429 | 0.511 | 0.346 | 0.271 | 0.361 | 0.286 | 0.248 | 0.278 | 0.308 | 0.286 | | | | | | | | | | | | | | | | | | | | | | | | | | | | | |
| 29 MN555567_TZ25757_rodent_Acomys_wilsoni_Tanzania_2011 | 0.729 | 0.647 | 0.586 | 0.602 | 0.571 | 0.474 | 0.549 | 0.556 | 0.617 | 0.549 | 0.594 | 0.421 | 0.383 | 0.414 | 0.436 | 0.436 | 0.474 | 0.414 | 0.519 | 0.338 | 0.323 | 0.361 | 0.286 | 0.271 | 0.301 | 0.301 | 0.248 | 0.271 | | | | | | | | | | | | | | | | | | | | | | | | | | | | |
| 30 MG822666_rodent_Proechimys_semispinosus_Panama_2014 | 0.684 | 0.647 | 0.556 | 0.586 | 0.564 | 0.526 | 0.549 | 0.504 | 0.586 | 0.594 | 0.594 | 0.414 | 0.436 | 0.406 | 0.414 | 0.451 | 0.474 | 0.429 | 0.481 | 0.338 | 0.331 | 0.398 | 0.316 | 0.353 | 0.293 | 0.323 | 0.316 | 0.301 | 0.308 | | | | | | | | | | | | | | | | | | | | | | | | | | | |
| 31 MN564790_ETH674_rodent_Stenocephalemys_albipes_Ethiopia_2012 | 0.684 | 0.617 | 0.594 | 0.586 | 0.564 | 0.481 | 0.571 | 0.541 | 0.624 | 0.526 | 0.602 | 0.376 | 0.414 | 0.383 | 0.414 | 0.436 | 0.421 | 0.406 | 0.481 | 0.331 | 0.278 | 0.338 | 0.263 | 0.271 | 0.256 | 0.278 | 0.211 | 0.241 | 0.226 | 0.256 | | | | | | | | | | | | | | | | | | | | | | | | | | |
| 32 MG600413_rodent_Rhizomys_pruinosus_Vietnam_2015 | 0.677 | 0.647 | 0.571 | 0.586 | 0.586 | 0.549 | 0.564 | 0.511 | 0.624 | 0.564 | 0.586 | 0.451 | 0.436 | 0.421 | 0.421 | 0.459 | 0.459 | 0.406 | 0.489 | 0.346 | 0.316 | 0.368 | 0.278 | 0.331 | 0.278 | 0.286 | 0.316 | 0.286 | 0.323 | 0.278 | 0.271 | | | | | | | | | | | | | | | | | | | | | | | | | |
| 33 KC815310_rodent_Peromyscus_maniculatus_USA_2008 | 0.669 | 0.617 | 0.564 | 0.609 | 0.579 | 0.489 | 0.556 | 0.541 | 0.609 | 0.534 | 0.594 | 0.391 | 0.421 | 0.398 | 0.414 | 0.451 | 0.474 | 0.414 | 0.511 | 0.368 | 0.286 | 0.383 | 0.293 | 0.293 | 0.263 | 0.256 | 0.263 | 0.278 | 0.241 | 0.256 | 0.173 | 0.301 | | | | | | | | | | | | | | | | | | | | | | | | |
| 34 KY370094_rodent_Neodon_clarkei_China_2014 | 0.684 | 0.624 | 0.586 | 0.594 | 0.571 | 0.504 | 0.556 | 0.511 | 0.609 | 0.534 | 0.602 | 0.391 | 0.414 | 0.429 | 0.383 | 0.421 | 0.459 | 0.406 | 0.459 | 0.353 | 0.278 | 0.376 | 0.301 | 0.263 | 0.278 | 0.286 | 0.256 | 0.248 | 0.241 | 0.263 | 0.203 | 0.233 | 0.218 | | | | | | | | | | | | | | | | | | | | | | | |
| 35 KC411784_rodent_Myodes_glareolus_Netherlands_2007 | 0.699 | 0.647 | 0.579 | 0.602 | 0.579 | 0.549 | 0.579 | 0.526 | 0.624 | 0.586 | 0.609 | 0.436 | 0.406 | 0.451 | 0.429 | 0.466 | 0.474 | 0.414 | 0.489 | 0.376 | 0.286 | 0.376 | 0.278 | 0.316 | 0.286 | 0.301 | 0.286 | 0.263 | 0.248 | 0.256 | 0.218 | 0.256 | 0.256 | 0.173 | | | | | | | | | | | | | | | | | | | | | | |
| 36 KC796074_bat_Hipposideros_vittatus_Kenya_2011 | 0.699 | 0.647 | 0.617 | 0.617 | 0.571 | 0.586 | 0.556 | 0.594 | 0.609 | 0.571 | 0.624 | 0.541 | 0.526 | 0.534 | 0.459 | 0.534 | 0.564 | 0.534 | 0.579 | 0.571 | 0.519 | 0.526 | 0.481 | 0.541 | 0.541 | 0.541 | 0.526 | 0.526 | 0.549 | 0.541 | 0.541 | 0.526 | 0.549 | 0.504 | 0.489 | | | | | | | | | | | | | | | | | | | | | |
| 37 KC796078_bat_Otomops_martiensseni_Kenya_2011 | 0.722 | 0.639 | 0.579 | 0.602 | 0.571 | 0.564 | 0.571 | 0.571 | 0.556 | 0.549 | 0.594 | 0.519 | 0.474 | 0.436 | 0.391 | 0.436 | 0.526 | 0.474 | 0.534 | 0.474 | 0.474 | 0.466 | 0.421 | 0.459 | 0.481 | 0.466 | 0.436 | 0.474 | 0.481 | 0.459 | 0.459 | 0.466 | 0.451 | 0.429 | 0.489 | 0.368 | | | | | | | | | | | | | | | | | | | | |
| 38 KP325401_horse_Equus_ferus_USA_2011 | 0.707 | 0.632 | 0.639 | 0.632 | 0.617 | 0.632 | 0.617 | 0.639 | 0.639 | 0.632 | 0.632 | 0.609 | 0.526 | 0.549 | 0.541 | 0.519 | 0.586 | 0.541 | 0.549 | 0.579 | 0.571 | 0.579 | 0.534 | 0.564 | 0.579 | 0.586 | 0.586 | 0.556 | 0.556 | 0.579 | 0.579 | 0.564 | 0.594 | 0.556 | 0.564 | 0.489 | 0.489 | | | | | | | | | | | | | | | | | | | |
| 39 EF108306_human_Homo_sapiens | 0.714 | 0.617 | 0.632 | 0.624 | 0.647 | 0.602 | 0.647 | 0.647 | 0.677 | 0.624 | 0.617 | 0.647 | 0.571 | 0.579 | 0.549 | 0.541 | 0.609 | 0.541 | 0.571 | 0.564 | 0.556 | 0.564 | 0.534 | 0.571 | 0.586 | 0.571 | 0.564 | 0.579 | 0.571 | 0.602 | 0.579 | 0.541 | 0.586 | 0.549 | 0.579 | 0.534 | 0.519 | 0.331 | | | | | | | | | | | | | | | | | | |
| 40 NC_009823_human_Homo_sapiens | 0.722 | 0.632 | 0.662 | 0.624 | 0.639 | 0.617 | 0.639 | 0.632 | 0.662 | 0.639 | 0.647 | 0.654 | 0.586 | 0.586 | 0.571 | 0.549 | 0.602 | 0.549 | 0.571 | 0.586 | 0.594 | 0.586 | 0.556 | 0.586 | 0.609 | 0.602 | 0.556 | 0.594 | 0.579 | 0.624 | 0.594 | 0.571 | 0.602 | 0.579 | 0.594 | 0.549 | 0.541 | 0.353 | 0.18 | | | | | | | | | | | | | | | | | | | | | |
| 41 NC_009824_human_Homo_sapiens | 0.729 | 0.669 | 0.647 | 0.632 | 0.632 | 0.632 | 0.684 | 0.639 | 0.654 | 0.632 | 0.639 | 0.662 | 0.586 | 0.594 | 0.556 | 0.571 | 0.609 | 0.571 | 0.579 | 0.579 | 0.571 | 0.564 | 0.556 | 0.571 | 0.579 | 0.571 | 0.564 | 0.579 | 0.579 | 0.624 | 0.609 | 0.541 | 0.602 | 0.586 | 0.586 | 0.556 | 0.526 | 0.353 | 0.188 | 0.226 | | | | | | | | | | | | | | | | | | | | |
| 42 NC_009826_human_Homo_sapiens | 0.744 | 0.639 | 0.647 | 0.609 | 0.602 | 0.609 | 0.669 | 0.632 | 0.647 | 0.609 | 0.624 | 0.647 | 0.571 | 0.586 | 0.556 | 0.541 | 0.617 | 0.556 | 0.571 | 0.579 | 0.586 | 0.571 | 0.564 | 0.586 | 0.624 | 0.617 | 0.586 | 0.617 | 0.617 | 0.632 | 0.617 | 0.579 | 0.624 | 0.594 | 0.594 | 0.541 | 0.549 | 0.353 | 0.226 | 0.203 | 0.226 | | | | | | | | | | | | | | | | | | | |
| 43 NC_009827_human_Homo_sapiens | 0.714 | 0.654 | 0.632 | 0.624 | 0.617 | 0.617 | 0.647 | 0.617 | 0.654 | 0.617 | 0.624 | 0.624 | 0.571 | 0.556 | 0.541 | 0.519 | 0.586 | 0.541 | 0.526 | 0.571 | 0.556 | 0.556 | 0.519 | 0.549 | 0.564 | 0.564 | 0.564 | 0.564 | 0.571 | 0.594 | 0.586 | 0.549 | 0.594 | 0.579 | 0.579 | 0.519 | 0.511 | 0.331 | 0.195 | 0.18 | 0.158 | 0.218 | | | | | | | | | | | | | | | | | | |
| 44 NC_009825_human_Homo_sapiens | 0.714 | 0.669 | 0.632 | 0.647 | 0.617 | 0.594 | 0.654 | 0.632 | 0.639 | 0.609 | 0.647 | 0.639 | 0.579 | 0.579 | 0.556 | 0.556 | 0.594 | 0.549 | 0.586 | 0.571 | 0.556 | 0.564 | 0.549 | 0.564 | 0.579 | 0.571 | 0.579 | 0.571 | 0.586 | 0.609 | 0.579 | 0.571 | 0.594 | 0.571 | 0.602 | 0.541 | 0.534 | 0.368 | 0.218 | 0.226 | 0.218 | 0.233 | 0.248 | | | | | | | | | | | | | | | | | |
| 45 M62321_HCV_reference | 0.737 | 0.632 | 0.639 | 0.609 | 0.639 | 0.639 | 0.669 | 0.624 | 0.647 | 0.624 | 0.654 | 0.662 | 0.594 | 0.594 | 0.579 | 0.571 | 0.609 | 0.556 | 0.564 | 0.602 | 0.602 | 0.602 | 0.571 | 0.579 | 0.617 | 0.617 | 0.594 | 0.609 | 0.609 | 0.632 | 0.609 | 0.594 | 0.602 | 0.609 | 0.617 | 0.534 | 0.549 | 0.376 | 0.226 | 0.195 | 0.241 | 0.226 | 0.188 | 0.226 | | | | | | | | | | | | | | | | |
| 46 MG599997_fish/rept/turtle_Chimaera_sp_China | 0.759 | 0.639 | 0.564 | 0.541 | 0.541 | 0.511 | 0.534 | 0.556 | 0.504 | 0.481 | 0.489 | 0.571 | 0.579 | 0.526 | 0.541 | 0.541 | 0.564 | 0.586 | 0.549 | 0.586 | 0.541 | 0.586 | 0.556 | 0.549 | 0.579 | 0.571 | 0.571 | 0.571 | 0.571 | 0.541 | 0.571 | 0.549 | 0.571 | 0.556 | 0.571 | 0.579 | 0.541 | 0.609 | 0.609 | 0.609 | 0.647 | 0.624 | 0.609 | 0.624 | 0.602 | | | | | | | | | | | | | | | |
| 47 MG599996_fish/rept/turtle_Chimaera_sp_China | 0.759 | 0.617 | 0.504 | 0.541 | 0.556 | 0.519 | 0.519 | 0.526 | 0.489 | 0.451 | 0.496 | 0.579 | 0.571 | 0.534 | 0.564 | 0.549 | 0.579 | 0.564 | 0.571 | 0.624 | 0.519 | 0.556 | 0.541 | 0.564 | 0.571 | 0.571 | 0.564 | 0.564 | 0.586 | 0.526 | 0.564 | 0.549 | 0.534 | 0.556 | 0.564 | 0.519 | 0.504 | 0.609 | 0.609 | 0.632 | 0.639 | 0.624 | 0.602 | 0.639 | 0.632 | 0.263 | | | | | | | | | | | | | | |
| 48 MG599995_fish/rept/turtle_Squalus_brevirostris_China | 0.752 | 0.647 | 0.556 | 0.579 | 0.602 | 0.579 | 0.571 | 0.534 | 0.541 | 0.504 | 0.504 | 0.586 | 0.586 | 0.549 | 0.564 | 0.549 | 0.602 | 0.609 | 0.571 | 0.564 | 0.549 | 0.632 | 0.586 | 0.579 | 0.586 | 0.586 | 0.586 | 0.564 | 0.571 | 0.556 | 0.564 | 0.556 | 0.541 | 0.549 | 0.556 | 0.632 | 0.579 | 0.639 | 0.617 | 0.602 | 0.639 | 0.609 | 0.609 | 0.632 | 0.602 | 0.406 | 0.444 | | | | | | | | | | | | | |
| 49 MG599994_fish/rept/turtle_Mustelus_manazo_China | 0.759 | 0.571 | 0.586 | 0.579 | 0.624 | 0.579 | 0.571 | 0.564 | 0.602 | 0.519 | 0.459 | 0.564 | 0.556 | 0.579 | 0.564 | 0.594 | 0.602 | 0.564 | 0.617 | 0.594 | 0.534 | 0.594 | 0.519 | 0.556 | 0.556 | 0.549 | 0.564 | 0.534 | 0.541 | 0.564 | 0.541 | 0.541 | 0.534 | 0.526 | 0.526 | 0.594 | 0.556 | 0.624 | 0.564 | 0.617 | 0.617 | 0.602 | 0.617 | 0.617 | 0.632 | 0.474 | 0.459 | 0.459 | | | | | | | | | | | | |
| 50 MG599999_fish/rept/turtle_Pelodiscus_sinensis_China | 0.707 | 0.617 | 0.534 | 0.541 | 0.564 | 0.511 | 0.579 | 0.579 | 0.571 | 0.579 | 0.579 | 0.549 | 0.534 | 0.511 | 0.459 | 0.504 | 0.549 | 0.534 | 0.519 | 0.564 | 0.526 | 0.571 | 0.511 | 0.541 | 0.549 | 0.526 | 0.541 | 0.549 | 0.534 | 0.519 | 0.519 | 0.519 | 0.511 | 0.504 | 0.519 | 0.564 | 0.526 | 0.609 | 0.579 | 0.602 | 0.602 | 0.602 | 0.602 | 0.602 | 0.609 | 0.511 | 0.489 | 0.556 | 0.594 | | | | | | | | | | | |
| 51 MG600000_fish/rept/turtle_Mauremys_megalocephala_China | 0.729 | 0.677 | 0.564 | 0.519 | 0.549 | 0.571 | 0.571 | 0.534 | 0.586 | 0.602 | 0.571 | 0.549 | 0.564 | 0.489 | 0.489 | 0.504 | 0.549 | 0.549 | 0.526 | 0.534 | 0.541 | 0.541 | 0.496 | 0.571 | 0.534 | 0.541 | 0.556 | 0.534 | 0.549 | 0.511 | 0.534 | 0.504 | 0.556 | 0.549 | 0.519 | 0.556 | 0.519 | 0.594 | 0.586 | 0.624 | 0.617 | 0.632 | 0.602 | 0.609 | 0.639 | 0.511 | 0.534 | 0.564 | 0.579 | 0.248 | | | | | | | | | | |
| 52 KY370091_rodent_Allactaga_sibirica_China_2014 | 0.684 | 0.609 | 0.541 | 0.609 | 0.579 | 0.496 | 0.579 | 0.556 | 0.564 | 0.541 | 0.579 | 0.541 | 0.444 | 0.444 | 0.451 | 0.519 | 0.549 | 0.429 | 0.474 | 0.459 | 0.474 | 0.496 | 0.444 | 0.459 | 0.481 | 0.459 | 0.489 | 0.459 | 0.451 | 0.474 | 0.466 | 0.474 | 0.481 | 0.466 | 0.436 | 0.534 | 0.541 | 0.579 | 0.549 | 0.579 | 0.586 | 0.594 | 0.579 | 0.549 | 0.602 | 0.489 | 0.534 | 0.549 | 0.541 | 0.398 | 0.451 | | | | | | | | | |
| 53 KC411777_rodent_Myodes_glareolus_Germany_2010 | 0.722 | 0.639 | 0.564 | 0.541 | 0.549 | 0.474 | 0.579 | 0.541 | 0.594 | 0.549 | 0.556 | 0.481 | 0.504 | 0.429 | 0.481 | 0.489 | 0.579 | 0.474 | 0.526 | 0.519 | 0.504 | 0.541 | 0.459 | 0.466 | 0.481 | 0.481 | 0.481 | 0.504 | 0.474 | 0.504 | 0.474 | 0.504 | 0.504 | 0.474 | 0.474 | 0.564 | 0.541 | 0.594 | 0.594 | 0.617 | 0.624 | 0.579 | 0.602 | 0.586 | 0.624 | 0.504 | 0.526 | 0.526 | 0.526 | 0.414 | 0.451 | 0.338 | | | | | | | | |
| 54 MN564792_TA132_rodent_Praomys_jacksoni_Tanzania_2013 | 0.699 | 0.609 | 0.526 | 0.564 | 0.526 | 0.519 | 0.556 | 0.526 | 0.594 | 0.549 | 0.541 | 0.459 | 0.481 | 0.444 | 0.466 | 0.519 | 0.556 | 0.466 | 0.541 | 0.496 | 0.466 | 0.511 | 0.436 | 0.459 | 0.451 | 0.436 | 0.481 | 0.474 | 0.459 | 0.489 | 0.459 | 0.459 | 0.459 | 0.451 | 0.444 | 0.549 | 0.549 | 0.579 | 0.594 | 0.617 | 0.617 | 0.602 | 0.594 | 0.579 | 0.624 | 0.519 | 0.519 | 0.519 | 0.511 | 0.421 | 0.444 | 0.316 | 0.173 | | | | | | | |
| 55 MN587672_TA275_rodent_Lophuromys_laticeps_Tanzania_2013 | 0.669 | 0.647 | 0.564 | 0.579 | 0.534 | 0.511 | 0.541 | 0.556 | 0.594 | 0.564 | 0.579 | 0.459 | 0.481 | 0.421 | 0.429 | 0.489 | 0.564 | 0.451 | 0.526 | 0.459 | 0.444 | 0.496 | 0.406 | 0.444 | 0.436 | 0.444 | 0.444 | 0.459 | 0.444 | 0.436 | 0.451 | 0.459 | 0.459 | 0.459 | 0.421 | 0.549 | 0.511 | 0.556 | 0.586 | 0.586 | 0.624 | 0.609 | 0.594 | 0.586 | 0.624 | 0.526 | 0.541 | 0.541 | 0.526 | 0.421 | 0.429 | 0.331 | 0.18 | 0.203 | | | | | | |
| 56 MN587655_CRT352-B_rodent_Lophuromys_dudui_DRC_2010 | 0.692 | 0.647 | 0.571 | 0.579 | 0.519 | 0.511 | 0.534 | 0.556 | 0.602 | 0.571 | 0.579 | 0.474 | 0.474 | 0.406 | 0.436 | 0.489 | 0.556 | 0.451 | 0.549 | 0.459 | 0.451 | 0.489 | 0.406 | 0.436 | 0.436 | 0.436 | 0.451 | 0.459 | 0.421 | 0.451 | 0.421 | 0.444 | 0.444 | 0.444 | 0.406 | 0.556 | 0.519 | 0.556 | 0.571 | 0.571 | 0.609 | 0.602 | 0.594 | 0.571 | 0.617 | 0.526 | 0.549 | 0.541 | 0.519 | 0.421 | 0.436 | 0.323 | 0.195 | 0.211 | 0.053 | | | | | |
| 57 MN587683_TA529-B_rodent_Lophuromys_machangui_Tanzania_2013 | 0.677 | 0.647 | 0.564 | 0.586 | 0.519 | 0.511 | 0.549 | 0.564 | 0.594 | 0.571 | 0.564 | 0.481 | 0.466 | 0.406 | 0.429 | 0.496 | 0.549 | 0.444 | 0.549 | 0.451 | 0.444 | 0.481 | 0.406 | 0.429 | 0.429 | 0.429 | 0.451 | 0.451 | 0.421 | 0.436 | 0.429 | 0.444 | 0.444 | 0.451 | 0.406 | 0.541 | 0.504 | 0.549 | 0.571 | 0.579 | 0.594 | 0.602 | 0.586 | 0.571 | 0.617 | 0.526 | 0.549 | 0.541 | 0.519 | 0.429 | 0.436 | 0.316 | 0.195 | 0.203 | 0.053 | 0.038 | | | | |
| 58 MN587662_CRT64-B_rodent_Lophuromys_dudui_DRC_2010 | 0.684 | 0.639 | 0.564 | 0.579 | 0.511 | 0.519 | 0.541 | 0.564 | 0.594 | 0.571 | 0.579 | 0.489 | 0.481 | 0.414 | 0.436 | 0.496 | 0.556 | 0.444 | 0.549 | 0.451 | 0.436 | 0.481 | 0.406 | 0.436 | 0.436 | 0.436 | 0.451 | 0.466 | 0.421 | 0.444 | 0.429 | 0.451 | 0.444 | 0.444 | 0.398 | 0.541 | 0.534 | 0.564 | 0.579 | 0.579 | 0.609 | 0.609 | 0.586 | 0.579 | 0.617 | 0.519 | 0.549 | 0.541 | 0.519 | 0.436 | 0.451 | 0.331 | 0.188 | 0.203 | 0.06 | 0.053 | 0.038 | | | |
| 59 MN587670_MOZ329-B_rodent_Lophuromys_machangui_Mozambique_201 | 0.677 | 0.639 | 0.564 | 0.579 | 0.526 | 0.519 | 0.541 | 0.564 | 0.586 | 0.564 | 0.564 | 0.489 | 0.466 | 0.421 | 0.421 | 0.496 | 0.549 | 0.436 | 0.549 | 0.451 | 0.444 | 0.489 | 0.414 | 0.444 | 0.436 | 0.429 | 0.459 | 0.459 | 0.429 | 0.444 | 0.436 | 0.459 | 0.451 | 0.459 | 0.414 | 0.541 | 0.511 | 0.556 | 0.571 | 0.586 | 0.602 | 0.609 | 0.586 | 0.571 | 0.617 | 0.519 | 0.541 | 0.541 | 0.519 | 0.429 | 0.436 | 0.323 | 0.195 | 0.211 | 0.053 | 0.068 | 0.03 | 0.03 | | |
| 60 MN587666_CRT74_rodent_Lophuromys_dudui_DRC_2010 | 0.677 | 0.639 | 0.564 | 0.586 | 0.526 | 0.511 | 0.549 | 0.564 | 0.594 | 0.571 | 0.571 | 0.481 | 0.474 | 0.414 | 0.421 | 0.496 | 0.541 | 0.436 | 0.549 | 0.444 | 0.444 | 0.481 | 0.406 | 0.436 | 0.429 | 0.429 | 0.451 | 0.451 | 0.421 | 0.436 | 0.429 | 0.451 | 0.444 | 0.451 | 0.406 | 0.541 | 0.511 | 0.556 | 0.579 | 0.586 | 0.609 | 0.609 | 0.586 | 0.579 | 0.617 | 0.526 | 0.549 | 0.541 | 0.519 | 0.436 | 0.436 | 0.323 | 0.195 | 0.211 | 0.038 | 0.045 | 0.023 | 0.038 | 0.023 | |

Estimates of Evolutionary Divergence between Sequences, NS5B region

**Supplementary table S7:** Complete list of all rodent hepacivirus co-infections identified in this study.

| Accession number | Specimen voucher | Co-infections | Host family | Host species | Country | Sampling year |
|---|---|---|---|---|---|---|
| MN587650 | CRT125 | CRT125, strain 1 | Muridae | *Lophuromys dudui* | DRC | 2010 |
| MN587651 | | CRT125, strain 2 | Muridae | *Lophuromys dudui* | DRC | 2010 |
| MN587652 | | CRT125, strain 3 | Muridae | *Lophuromys dudui* | DRC | 2010 |
| MN587653 | | CRT125, strain 4 | Muridae | *Lophuromys dudui* | DRC | 2010 |
| MN587654 | CRT352 | CRT352, strain 1 | Muridae | *Lophuromys dudui* | DRC | 2010 |
| MN587655 | | CRT352, strain 2 | Muridae | *Lophuromys dudui* | DRC | 2010 |
| MN587658 | CRT490 | CRT490, strain 1 | Muridae | *Lophuromys dudui* | DRC | 2010 |
| MN587659 | | CRT490, strain 2 | Muridae | *Lophuromys dudui* | DRC | 2010 |
| MN587660 | | CRT490, strain 3 | Muridae | *Lophuromys dudui* | DRC | 2010 |
| MN587661 | CRT64 | CRT64, strain 1 | Muridae | *Lophuromys dudui* | DRC | 2010 |
| MN587662 | | CRT64, strain 2 | Muridae | *Lophuromys dudui* | DRC | 2010 |
| MN587663 | | CRT64, strain 3 | Muridae | *Lophuromys dudui* | DRC | 2010 |
| MN587664 | | CRT64, strain 4 | Muridae | *Lophuromys dudui* | DRC | 2010 |
| MN587665 | | CRT64, strain 5 | Muridae | *Lophuromys dudui* | DRC | 2010 |
| MN587669 | MOZ329 | MOZ329, strain 1 | Muridae | *Lophuromys machangui* | Mozambique | 2011 |
| MN587670 | | MOZ329, strain 2 | Muridae | *Lophuromys machangui* | Mozambique | 2011 |
| MN587671 | | MOZ329, strain 3 | Muridae | *Lophuromys machangui* | Mozambique | 2011 |
| MN587696 | TA100 | TA100, strain 1 | Muridae | *Lophuromys stanleyi* | Tanzania | 2013 |
| MN587697 | | TA100, strain 2 | Muridae | *Lophuromys stanleyi* | Tanzania | 2013 |
| MN587698 | | TA100, strain 3 | Muridae | *Lophuromys stanleyi* | Tanzania | 2013 |
| MN587673 | TA293 | TA293, strain 1 | Muridae | *Lophuromys laticeps* | Tanzania | 2013 |
| MN587674 | | TA293, strain 2 | Muridae | *Lophuromys laticeps* | Tanzania | 2013 |
| MN587675 | | TA293, strain 3 | Muridae | *Lophuromys laticeps* | Tanzania | 2013 |
| MN587676 | | TA293, strain 4 | Muridae | *Lophuromys laticeps* | Tanzania | 2013 |
| MN587677 | | TA293, strain 5 | Muridae | *Lophuromys laticeps* | Tanzania | 2013 |
| MN587678 | TA498 | TA498, strain 1 | Muridae | *Lophuromys machangui* | Tanzania | 2013 |
| MN587679 | | TA498, strain 2 | Muridae | *Lophuromys machangui* | Tanzania | 2013 |
| MN587680 | | TA498, strain 3 | Muridae | *Lophuromys machangui* | Tanzania | 2013 |
| MN587682 | TA529 | TA529, strain 1 | Muridae | *Lophuromys machangui* | Tanzania | 2013 |
| MN587683 | | TA529, strain 2 | Muridae | *Lophuromys machangui* | Tanzania | 2013 |
| MN587684 | | TA529, strain 3 | Muridae | *Lophuromys machangui* | Tanzania | 2013 |
| MN587685 | | TA529, strain 4 | Muridae | *Lophuromys machangui* | Tanzania | 2013 |
| MN587686 | TA531 | TA531, strain 1 | Muridae | *Lophuromys machangui* | Tanzania | 2013 |
| MN587687 | | TA531, strain 2 | Muridae | *Lophuromys machangui* | Tanzania | 2013 |
| MN587688 | | TA531, strain 3 | Muridae | *Lophuromys machangui* | Tanzania | 2013 |
| MN587689 | | TA531, strain 4 | Muridae | *Lophuromys machangui* | Tanzania | 2013 |
| MN587690 | | TA531, strain 5 | Muridae | *Lophuromys machangui* | Tanzania | 2013 |
| MN587691 | TA532 | TA532, strain 1 | Muridae | *Lophuromys machangui* | Tanzania | 2013 |
| MN587692 | | TA532, strain 2 | Muridae | *Lophuromys machangui* | Tanzania | 2013 |
| MN587693 | | TA532, strain 3 | Muridae | *Lophuromys machangui* | Tanzania | 2013 |
| MN587694 | | TA532, strain 4 | Muridae | *Lophuromys machangui* | Tanzania | 2013 |
